# Toward Best Practice in Identifying Subtle Differential Expression with RNA-seq: A Real-World Multi-Center Benchmarking Study Using Quartet and MAQC Reference Materials

**DOI:** 10.1101/2023.12.09.570956

**Authors:** Duo Wang, Yaqing Liu, Yuanfeng Zhang, Qingwang Chen, Yanxi Han, Wanwan Hou, Cong Liu, Yin Yu, Ziyang Li, Ziqiang Li, Jiaxin Zhao, Yuanting Zheng, Leming Shi, Jinming Li, Rui Zhang

## Abstract

Translating RNA-seq into clinical diagnostics requires ensuring the reliability of detecting clinically relevant subtle differential expressions, such as those between different disease subtypes or stages. Moreover, cross-laboratory reproducibility and consistency under diverse experimental and bioinformatics workflows urgently need to be addressed. As part of the Quartet project, we presented a comprehensive RNA-seq benchmarking study utilizing Quartet and MAQC RNA reference samples spiked with ERCC controls in 45 independent laboratories, each employing their in-house RNA-seq workflows. We assessed the data quality, accuracy and reproducibility of gene expression and differential gene expression and compared over 40 experimental processes and 140 combined differential analysis pipelines based on multiple ‘ground truths’. Here we show that real-world RNA-seq exhibited greater inter-laboratory variations when detecting subtle differential expressions between Quartet samples. Experimental factors including mRNA enrichment methods and strandedness, and each bioinformatics step, particularly normalization, emerged as primary sources of variations in gene expression and have a more pronounced impact on the subtle differential expression measurement. We underscored the pivotal role of experimental execution over the choice of experimental protocols, the importance of strategies for filtering low-expression genes, and optimal gene annotation and analysis tools. In summary, this study provided best practice recommendations for the development, optimization, and quality control of RNA-seq for clinical diagnostic purposes.

## Introduction

Transcriptome sequencing (RNA-seq) has expanded new avenues for exploring global expression patterns as well as identifying alternative splicing events ^1^. Differential expression analysis of transcriptomic data enables genome-wide identification of gene or isoform expression changes associated with biological conditions of interest. This contributes significantly to the discovery of biomarkers for disease diagnosis ^2^, prognosis ^3^, and therapeutic selection ^4^. These evidences facilitate the application of RNA-seq in clinical routine. Noticeably, clinically relevant biological differences among study groups are often small, manifested by fewer differentially expressed genes, especially between certain disease and normal tissues ^5, 6^, or between different disease subtypes or stages ^7–11^. Such subtle differential gene expression is typically challenging to distinguish from noises of technical replicates. Therefore, translating RNA-seq into clinical diagnostics poses requirements for more sensitive differential expression analysis, emphasizing the necessity for quality assessment at subtle differential expression levels.

However, over the past decade, quality assessment of RNA-seq in the community has predominantly relied on milestone MAQC reference materials, characterized by significantly large biological differences between samples, which were developed by the MicroArray/Sequencing Quality Control (SEQC/MAQC) Consortium from ten cancer cell lines and brain tissues of 23 donors ^12^. The MAQC Consortium utilized these samples with spike-ins of 92 synthetic RNA from the External RNA Control Consortium (ERCC) to assess RNA-seq performance and demonstrated a high accuracy and reproducibility of relative expression measurements across different sites and platforms under appropriate data processing and analysis conditions ^13, 14^. More large-scale studies have also employed these two RNA reference materials to compare different library preparation protocols and sequencing platforms ^15–17^, and have utilized the MAQC datasets for benchmarking bioinformatics pipelines ^18–21^. Moreover, the Genetic European Variation in Disease, a European Medical Sequencing (GEUVADIS) Consortium sequenced RNA samples from lymphoblastoid cell lines of 465 individuals across seven sites to assess reproducibility across laboratories and examine the sources of inter-laboratory variation under an identical experimental and bioinformatics process ^22^.

Noticeably, quality control based on the MAQC reference materials may not fully ensure the accurate identification of clinically relevant subtle differential expression ^23^. Moreover, in contrast to the rigorously controlled RNA-seq workflows of previous study designs, the real-world scenarios present significant differences in sample processing, experimental protocols, sequencing platforms, and analysis pipelines across laboratories, where confounding factors may compromise the accuracy and reproducibility of RNA-seq ^14, 15, 22^. In the context of such diverse experimental and bioinformatics processes, understanding of the sources of inter-laboratory variation remains limited. Therefore, a detailed quality assessment of the overall performance of real-world RNA-seq in detecting subtle differential expression for clinical diagnostic purposes and of the technical factors affecting diagnostic performance is necessary.

Recently, the Quartet project for quality control and data integration of multi-omics profiling, introduced multi-omics reference materials derived from immortalized B-lymphoblastoid cell lines from a Chinese quartet family of parents and monozygotic twin daughters, and developed ratio-based reference datasets ^24^. These well-characterized, homogenous, and stable Quartet RNA reference materials with small inter-sample biological differences, provided a unique opportunity for the assessment and benchmarking of transcriptome profiling at subtle differential expression levels in a reference-based manner ^23^.

Within the scope of the Quartet project, this study utilized Quartet RNA samples with spike-ins of ERCC controls, and MAQC RNA samples to generate RNA-seq data across 45 independent laboratories, each using its own in-house experimental protocol and analysis pipeline. Overall, approximately 120 billion reads of RNA-seq data were generated and analyzed, representing the most extensive effort to conduct an in-depth exploration of transcriptome data to date. Through the quality assessment based on Quartet and MAQC samples in parallel, this study thoroughly elucidated the performance of real-world RNA-seq, particularly when detecting subtle differential expression levels. Subsequently, we leveraged gene expression data from over 40 different experimental processes and 140 differential analysis pipelines to investigate sources of variation at experimental and bioinformatics aspects, respectively. This study provides best practice recommendations for the experimental and bioinformatics designs of the RNA-seq toward the scientific question addressed, and underscores the necessity of quality controls at subtle differential expression levels through the comparisons of Quartet and MAQC reference materials.

## Results

### Study design

Our multi-center study involved four well-characterized Quartet RNA samples (M8, F7, D5 and D6) with ERCC spike-in RNA controls added to M8 and D6 samples, T1 and T2 samples constructed by mixing M8 and D6 at the defined ratios of 3:1 and 1:3, respectively, and MAQC RNA samples A and B (**Fig. 1a**). The sample panel design introduces various types of ‘ground truth’, encompassing three reference datasets: ratio-based Quartet reference datasets, TaqMan datasets for Quartet and MAQC samples, and ‘built-in truth’ involving ERCC spike-in ratios and known mixing ratios for the T1 and T2 samples **(Supplementary Notes, section 2.1)**. Each sample was provided with three technical replicates, resulting in a total of 24 RNA samples, which were sequenced and analyzed by 45 independent laboratories. Each laboratory employed distinct RNA-seq workflows, involving different RNA processing methods, library preparation protocols, sequencing platforms, and bioinformatics pipelines **(Supplementary Table 1)**. This approach accurately mirrored the actual research practices in real-world scenarios.

**Fig. 1.**
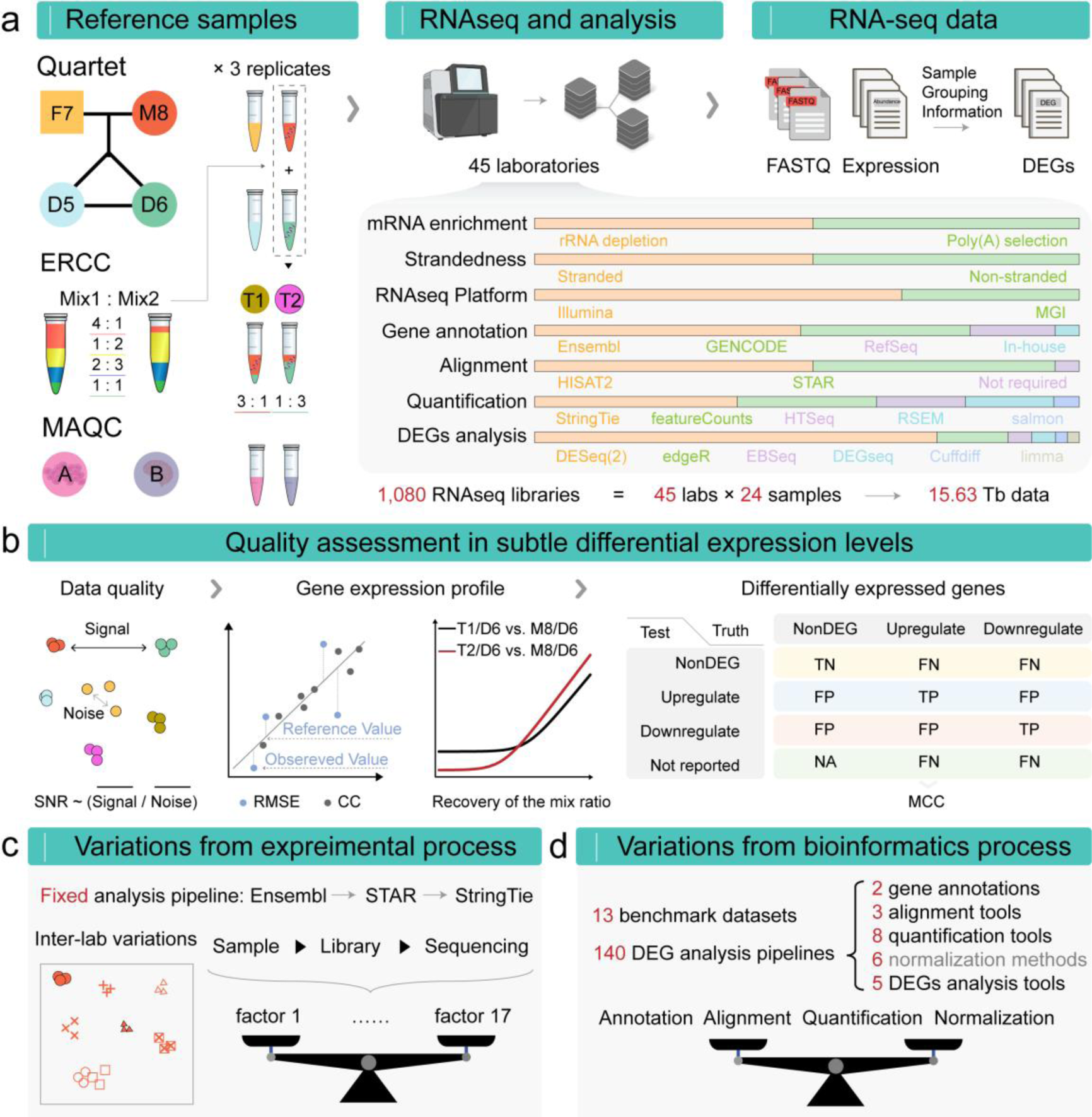
Overview of study design. **(a)** Two MAQC samples (A = Universal Human Reference RNA and B = Human Brain Reference RNA), two ERCC synthetic RNA mix, and Quartet RNA reference materials were utilized to prepare a set of samples. The M8 and D6 samples were combined with ERCC controls at manufacturer recommended amounts, and then mixed at 3:1 and 1:3 ratios to create sample T1 and T2, respectively. Each sample was prepared with three replicates, and tested by 45 laboratories with distinct protocols and analysis pipelines, resulting in a total of 1080 libraries and 15.63 Tb of data generated. All 45 laboratories submitted expression data and differential expression calls at gene and transcript levels, while 42 laboratories submitted complete raw sequencing data. DEG, differential expression gene. **(b)** A comprehensive framework for assessment of real-world RNA-seq data, encompassing assessment of data quality using PCA-based SNR, as well as gene expression profiles and differentially expressed genes by comparing with various ground truths. SNR, Signal-to-Noise Ratio; RMSE, Root Mean Square Error; CC, Correlation Coefficient; MCC, Matthews Correlation Coefficient; TN, True Negative; TP, True Positive; FN, False Negative; FP, False Positive. **(c)** A fixed analysis pipeline was applied to all raw data to exclude the influence of the bioinformatic process. Then the relative contributions of experimental factors to inter-laboratory variations were investigated. **(d)** High-quality data from 13 laboratories were selected for the benchmarking study, and the performance of 140 differential analysis pipelines composed of two gene annotations, three alignment tools, eight quantification tools following six normalization methods, and five differential analysis tools was compared to explore the sources of variations from the bioinformatics process.

Totally, 1,080 RNA-seq libraries were prepared, yielding a dataset of over 120 billion reads (15.63 Tb). Based on these extensive data for Quartet and MAQC samples, this study aimed to provide real-world evidence on the performance of RNA-seq in detecting both subtle and large differential expression by assessing data quality, and the accuracy and reproducibility of gene expression and differential expression calls (**Fig. 1b**). Moreover, a fixed analysis pipeline was applied for all RNA-seq raw data to exclusively investigate the sources of inter-laboratory variation from the experimental processes (**Fig. 1c**). A total of 140 different analysis pipelines consisting of two gene annotations, three alignment tools, eight quantification tools following six normalization methods, and five differential analysis tools were applied for high-quality benchmark datasets selected from 13 laboratories to investigate the sources of variation from the bioinformatics process (**Fig. 1d**).

### Basic quality control for raw reads and read alignment

We first assessed the sequencing quality properties of the RNA-seq data for the Quartet and MAQC samples, including sequencing depth, base quality, GC content, and duplicate rate **(Supplementary Table 2)**. The average sequencing depth ranged from 39.4 Mb to 418.8 Mb for Quartet samples and from 40.9 Mb to 424.2 Mb for MAQC samples across laboratories **(Supplementary Fig. 1)**. Within the same laboratory, different samples exhibited variations in sequencing depth, particularly noticeable for laboratories with higher average sequencing depths. Given that different flowcells or lanes can lead to variations in total reads counts, we compared 15 laboratories that assigned 24 libraries to two or more lanes to other laboratories that assigned libraries to a single lane, and observed no increased variations **(Supplementary Fig. 2)**. Therefore, inter-sample variations were considered to be due to difficulties of equimolar pooling ^22^. Both Quartet and MAQC samples exhibited high Q30 scores, ranging from 88.4% to 96.6% and from 88.3% to 96.7%, respectively, reflecting the high quality of base calling **(Supplementary Fig. 3)**. The base quality distribution of the first about 10 bases was relatively lower than the highest value in most laboratories for the Quartet and MAQC samples **(Supplementary Fig. 4–5**), which was attributed to the reverse transcriptase priming step ^15, 22^. We also observed that the quality scores of reverse reads were generally lower than those of forward reads in most laboratories, which was attributed to the decreased cluster size and higher number of errors due to more amplification steps before sequencing the reverse reads ^25^. GC content bias was found across laboratories, with the average GC content ranging from 42.3% to 54.2% for the Quartet samples and from 42.4% to 52.9% for the MAQC samples. Such laboratory-specific GC content bias, primarily caused by different sites of library preparation ^26^, was more noticeable than the sample-specific GC content bias **(Supplementary Fig. 6–7)**. Unusual GC content presents inherent challenges, as GC-poor genes (< 35%) tended to exhibit more variable expression levels between laboratories than genes with medium or high (> 65%) GC content **(Supplementary Fig. 8)**. The average duplication rates of the sequencing reads varied significantly across laboratories, ranging from 4.2% to 73.4% for the Quartet samples and from 5.0% to 75.5% for the MAQC samples **(Supplementary Fig. 9)**. Nine laboratories exhibited an average duplication rate exceeding 30%, surpassing the typical duplication levels observed in prior research ^15, 27, 28^. These extra duplicate reads may be due to PCR amplification bias rather than highly expressed genes ^29^.

We next assessed the alignment statistics after mapping the raw reads using STAR **(Online Methods)**. All laboratories exhibited a high overall alignment rate, ranging from 90.69% to 98.7% for the Quartet samples and from 92.1% to 98.9% for the MAQC samples **(Supplementary Fig. 10)**. The slightly lower uniquely mapping rate was noticeable in the Quartet samples in comparison to the MAQC samples, with average mapping rates of 89.7% (80.9%–95.4%) and 92.0% (84.1%–96.0%), respectively. This was similar to the common characteristics observed when comparing clinical samples with the MAQC samples ^20^. The multi-mapping rate seemed to be associated with the mRNA enrichment methods. The rRNA depletion method resulted in higher average multi-mapping rates than the poly(A) selection method **(Supplementary Fig. 11)**, possibly due to the capture of a greater number of small non-coding RNAs with high sequence similarity ^30^. Meanwhile, a high multi-mapping rate was consistently correlated with a higher mismatch rate **(Supplementary Fig. 10)**. The percentage of aligned reads mapping to annotated exons is directly related to expression quantification, and is therefore a critical quality metric. The poly(A) selection method consistently showed a higher median percentage of exonic reads at 84.5% and 80.9%, compared to the rRNA depletion method at 46.3% and 44.1% for the Quartet and MAQC samples, respectively **(Supplementary Fig. 12–13)**.

Additionally, the percentage of reads mapped to ERCC reference sequences allowed for the identification of problematic samples or libraries. In four samples (MAQC A, B, and Quartet F7, D5) without ERCC spike-ins, we observed reads counts ranging from 1 to 213,467 mapped to ERCC genes across 38 laboratories **(Supplementary Fig. 14)**. Particularly, Lab10 exhibited an exceptionally high fraction of ERCC reads in the two replicates of MAQC sample A, accounting for 0.8% and 0.06% of the exonic reads. This indicates potential contaminations across RNA samples or libraries ^31^.

### Significant variations in detecting subtle differential expression

We combined multiple metrics for a robust characterization of RNA-seq performance: (i) quality of gene expression data using signal-to-noise ratio (SNR) based on principal component analysis (PCA) ^23^, (ii) the accuracy and reproducibility of absolute and relative gene expression measurements based on several ‘ground truths’, and (iii) the accuracy of differentially expressed genes (DEGs) based on the reference datasets (**Fig. 1b**). These metrics constitutes a comprehensive performance assessment framework that captures different aspects of gene-level transcriptome profiling (**Supplementary Notes, section 2.2**).

PCA-based SNR values based on both the Quartet and MAQC samples discriminated all gene expression data into a wide range of quality levels, reflecting the varying ability to distinguish biological difference signals in different sample groups from technical noises in replicates (**Fig. 2a**). However, smaller intrinsic biological differences appeared to be more challenging to distinguish from noises, indicated by lower average SNR values for Quartet samples among laboratories compared to MAQC samples, at 19.8 (0.3–37.6) and 33.0 (11.2–45.2), respectively **(Supplementary Fig. 15)**. The reduced biological differences among the mixed samples led to a further decrease in the average SNR values to 18.2 (0.2–36.4). Particularly, for different laboratories, the gap between two sets of SNR values, one based on the Quartet and mixed samples and the other based on the MAQC samples, differed from 4.7 to 29.3, suggesting that diagnosing quality issues at subtle differential expression levels was sensitive. Moreover, SNR examinations allowed for the identification of random failures in the technical replicates. The SNR17 values, calculated from any 17 out of the 18 samples (12 Quartet and 6 mixed samples), increased by six decibels compared to the corresponding SNR18 values in six laboratories (**Fig. 2a**).

**Fig. 2.**
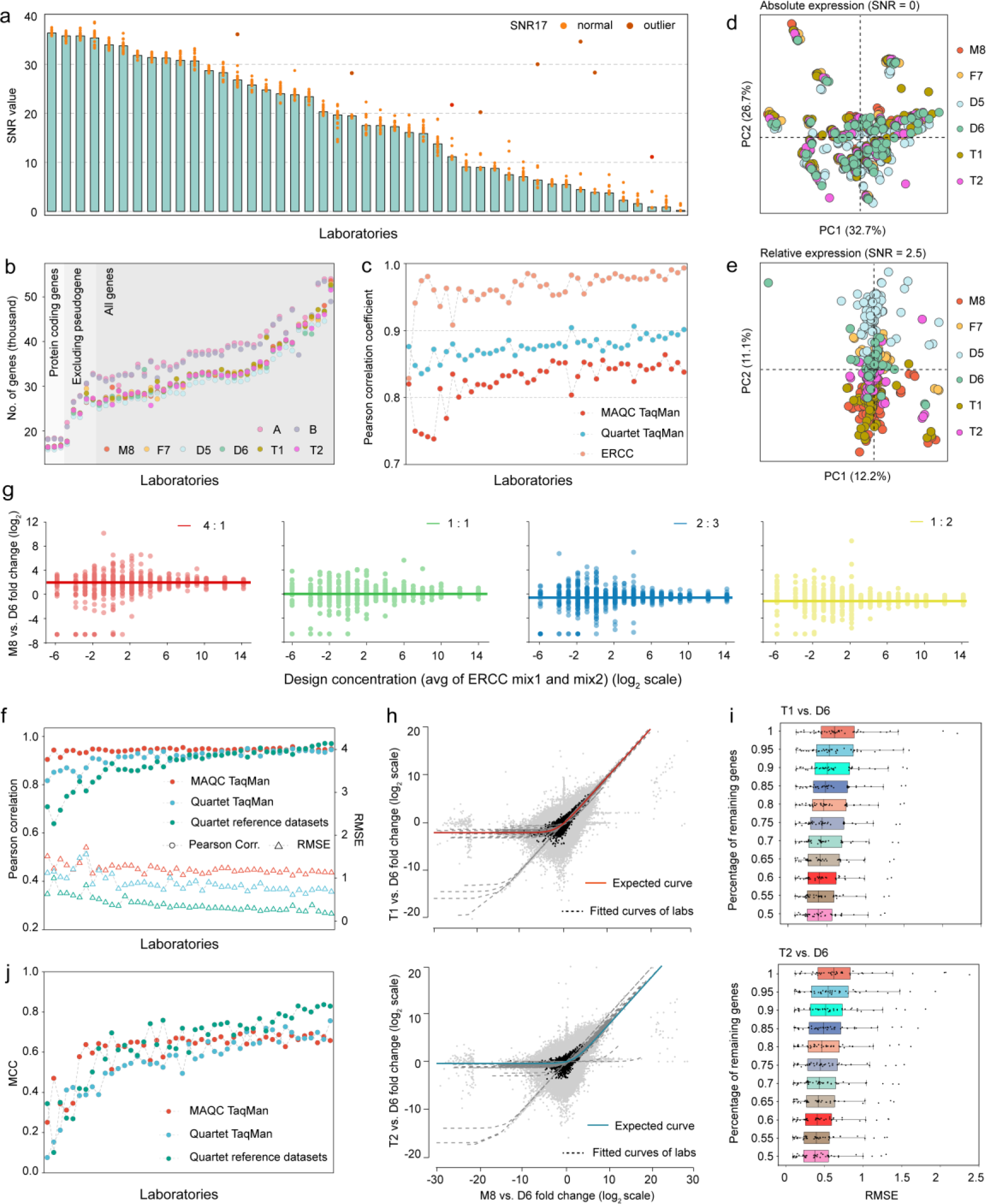
RNA-seq performance metrics for real-world laboratories. **(a)** SNR values across 45 laboratories to measure data quality. Laboratories were ordered by SNR values. Dots represented SNR values based on any 17 of the 18 samples (12 Quartet and 6 mixed samples) in each laboratory. A dot in dark red represented SNR17 value that increased over five decibels compared to its standard SNR (18-sample SNR), when one library in this laboratory was excluded, while a dot in orange represented SNR17 value that decreased or increased less five decibels compared to its standard SNR. **(b)** The gene types of interest for all laboratories and the corresponding number of genes supported by at least one reads for all three replicates **(Supplementary Notes, section 2.4)**. Three laboratories analyzed only protein-coding genes, five laboratories excluded pseudogenes from their analysis, and the remaining 37 laboratories analyzed all gene types. **(c)** Comparison of absolute expression levels to TaqMan datasets and ERCC concentrations on the log2 scale. **(d)** Scatterplots of PCA on RNA-seq data of all laboratories in absolute expression levels, **(e)** and relative expression levels. The circles of the same color represent all replicates across all laboratories for each sample. **(f)** Assessment of relative expression levels using Pearson correlation coefficient and the Root Mean Square Error (RMSE) based on Quartet reference datasets and TaqMan datasets on the log2 scale. **(g)** ERCC spike-in ratios can be recovered increasingly well at higher expression levels. **(h)** A consistency test for recovering the expected sample mixing ratio in samples T1 and T2. The red and cyan solid line traces the expected curve after mRNA/total-RNA shift correction. The grey dashed lines indicate the fitted curves from data of laboratories. The ERCC genes are shown in black, and the other human genes are shown in grey. **(i)** The ability to recover expected mixing ratios was measured using RMSE between the observed expression profiles and the expected expression profiles. As genes with low fold changes were progressively filtered out, the RMSE across all laboratories decreased, indicating an increase of accuracy. The different colors in the box plots represent varying percentage of filtered genes. **(j)** Comparison of differentially expressed genes to Quartet reference datasets and TaqMan datasets using Matthews Correlation Coefficients (MCC).

Gene expression measurements was assessed based on the Quartet reference datasets, TaqMan datasets, and the built-in truths including the ERCC spike-in ratios and mixed ratios of sample T1 and T2. Gene expression exhibited significant inter-laboratory variations, especially in absolute expression. Considering the varying gene types of interest among laboratories (**Fig. 2b**), only protein-coding genes were included to facilitate comparisons between laboratories. All laboratories exhibited lower correlation coefficients at 0.825 (0.738–0.856) with the MAQC TaqMan datasets of 830 protein-coding genes, compared to those at 0.876 (0.835–0.906) with the Quartet TaqMan datasets of 143 protein-coding genes (**Fig. 2c**). Correlations with the nominal concentrations of the 92 ERCC spike-in RNAs were consistently high for all laboratories with an average correlation coefficient of 0.964 (0.828–0.963). More ERCC based assessments are shown in **Supplementary Notes, Section 2.3**. These results indicate that accurate quantification of a broader set of genes is more challenging, highlighting the importance of large-scale reference datasets for performance assessment. We also focused on the absolute expression for other gene types, and observed that small non-coding RNAs exhibited the largest inter-laboratory variations, followed by pseudogenes, long non-coding RNAs, and immunoglobulin/T cell receptor segments **(Supplementary Fig. 16)**, which appeared to be associated with gene features specific to each type, such as gene lengths and gene expression levels **(Supplementary Fig. 17–18)**.

Relative expression measurements are more reliable than absolute expression measurements, but they still present challenges when identifying subtle differential expression. The variations in relative expression across laboratories decreased compared to those in absolute expression, as indicated by that samples tended to cluster based on the source sample rather than the laboratory in PCA analyses. However, laboratories still exhibited considerable variations in relative expression exceeding the small biological difference among the Quartet samples (**Fig. 2d–e and Supplementary Fig. 19)**. Despite the higher accuracy metrics among laboratories compared to absolute expression **(Supplementary Fig. 20)**, relative expression demonstrated lower average correlation coefficients of 0.865 (0.288–0.978) and 0.860 (0.488–0.944) with the Quartet reference datasets of 23790 protein-coding genes and Quartet TaqMan datasets, respectively, compared to the average correlation coefficient of 0.927 (0.778–0.949) with MAQC TaqMan datasets (**Fig. 2f**). It is noteworthy that the Root Mean Square Error (RMSE) values between laboratories and the Quartet reference datasets were the lowest, reflecting the systematic deviations between RNA-seq and TaqMan RT-qPCR assays but not between RNA-seq and the Quartet reference datasets (**Fig. 2f**). In addition, based on the ERCC spike-in ratios and mixing ratios of samples T1 and T2, we complementarily examined the accuracy and reproducibility of the relative expression across 92 ERCC RNAs and all detected genes. Our results revealed the impact of low gene expression and subtle differential expression on relative expression measurements. The expected ERCC spike-in ratios were more accurately recovered for high-concentration ERCC genes compared to low-concentration genes (**Fig. 2g**). The mixing ratios in the mixed samples were recovered well in most laboratories (**Fig. 2h**). Laboratories that failed to recover the mixing ratio demonstrated the presence of outliers **(Supplementary Fig. 21)**, which are typically caused by the erroneous detection or calculation of low-expressed genes **(Supplementary Fig. 22)**. By stepwise filtering of genes with low fold changes, the RMSE values between the observed and expected fold changes decreased, indicating a higher accuracy of gene expression measurements (**Fig. 2i**).

The DEGs calls revealed significant variations across laboratories in terms of both DEGs number and the accuracy of DEGs classification based on the Quartet reference datasets and TaqMan datasets. The number of protein-coding DEGs ranged from 787 to 13,194 for the Quartet and mixed samples, and from 4,275 to 12,773 for the MAQC samples **(Supplementary Fig. 23)**. As a result, true positives ranging from 0.03% to 78.6%, from 1.2% to 82.0%, and from 0.2% to 52.9% of the Quartet reference datasets, Quartet TaqMan, and MAQC TaqMan datasets, respectively, were missed by the laboratories. Consequently, we employed a penalized Matthews Correlation Coefficient (MCC) to assess the accuracy of DEGs calls (**Fig. 1b and Supplementary Notes, section 2.2)**. The MCC values based on the Quartet reference datasets and Quartet TaqMan datasets were more dispersed among laboratories, ranging from 0.100 to 0.837 and from 0.075 to 0.756, respectively (**Fig. 2j**). In contrast, the MCC values based on the MAQC TaqMan datasets ranged from 0.251 to 0.702. Importantly, the relatively low MCC values in certain laboratories could be explained by several factors **(Supplementary Table 3)**. For example, in the case of lab18, the expression data exhibited a SNR of 0.9, indicating that the low-quality library preparation or sequencing processes resulted in unreliable and uninformative RNA-seq data for differential analysis. The lab03 and lab04 demonstrated low accuracy of fold change determination, impacting the reliability of the DEG calls. Additionally, different thresholds to filter low-expression genes and cutoffs for DEGs identification led to variations in the number of DEGs, which collectively contributed to low accuracy.

### Sources of variation from the experimental process

The significant variations, especially at subtle differential expression levels, necessitated investigating the sources of variation. To exclusively focus on variation from the experimental process, we employed a uniform data analysis pipeline for all RNA-seq raw data, involving the use of fastp for data pre-processing, Ensembl gene annotation, STAR for reads alignment, and StringTie for gene quantification. When compared to the original expression data, the SNR values and accuracy metrics for gene expression measurements increased in most laboratories, indicating that the fixed pipeline was reliable for excluding the influence of diverse bioinformatics tools **(Supplementary Fig. 24–25**). These variations arising from different RNA processing methods, library preparation protocols, and sequencing platforms among laboratories represent ‘experimental noise’.

In the presence of significant inter-laboratory variations from the experimental process for both Quartet (**Fig. 3a**) and MAQC samples (**Fig. 3b**), experimental factors had a great impact on subtle differential expression measurement. We quantified the relative contribution of technical and biological factors to the total variations by principal variance component analysis (PVCA) based on absolute expression data from all laboratories for all samples. A total of 17 factors from the experimental process were considered **(Supplementary Table 4)**, and these experimental factors introduced significantly greater variations than biological differences among the Quartet samples (89.4% vs. 9.6%), with mRNA enrichment methods and strandedness as the primary sources (**Fig. 3c**). Additionally, library preparation kits, reads length, and the number of exonic reads also contributed to 17.9% of the variations. In contrast, while MAQC samples revealed similar sources, variations derived from experimental factors were lower than biological differences between the MAQC samples (45.3% vs. 54.7%) (**Fig. 3d**).

**Fig. 3.**
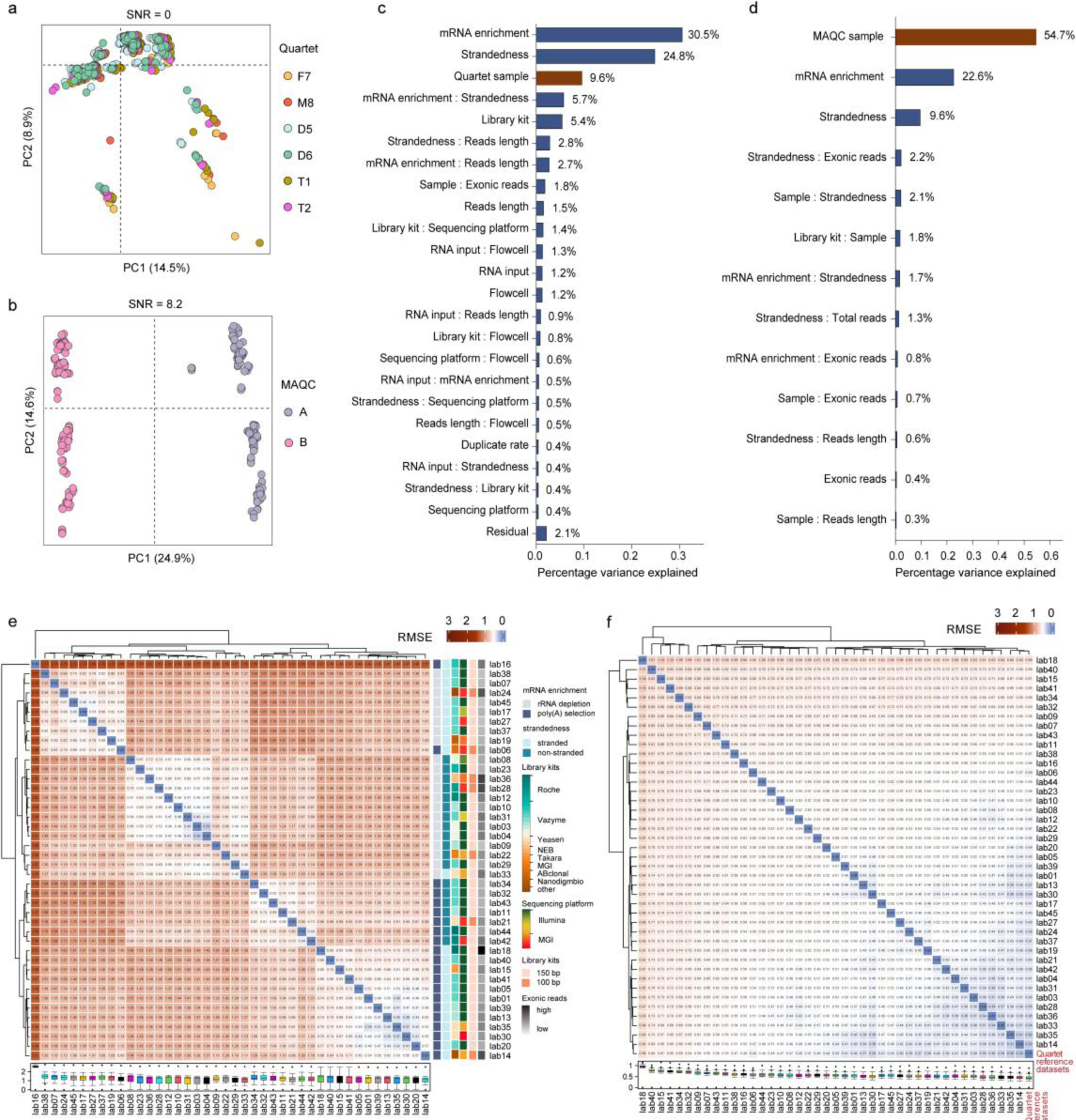
Sources of variation from the experimental process. **(a)** Scatterplots of PCA on RNA-seq data of all laboratories for Quartet samples, **(b)** and MAQC samples after applying fixed analysis pipeline. The circles of the same color represent all replicates across all laboratories for each sample. **(c)** Principal variance component analysis quantifies the proportion of variance explained by each experimental factor for Quartet samples, **(d)** and MAQC samples. **(e)** Heatmap and hierarchical clustering of different laboratories based on the RMSE at absolute expression levels, **(f)** and relative expression levels for Quartet samples. RMSE, Root Mean Square Error.

Relative expression could effectively correct for the influence of experimental factors, as indicated by a significant decrease of over 40% in the relative contribution of experimental factors to the variations for both the Quartet and MAQC samples. **(Supplementary Fig. 26)**. The increased consistency between any two laboratories compared to absolute expression further confirmed this (**Fig. 3e and 3f**). However, Quartet samples demonstrated that there remained 27.5% of unexplained variations that could not be eliminated, implying the presence of additional influencing factors within the complex and diverse experimental process.

## Sources of variation from the bioinformatics process

To assess the sources of variation from the bioinformatics process, high-quality data for Quartet and MAQC samples from 13 laboratories served as benchmark datasets, encompassing 13 different library preparation protocols, seven sequencing platforms, and a wide range of sequencing depths spanning 42.6 Mb to 425.3 Mb to mitigate bias **(Materials and Methods)**. Following commonly used transcriptomic profiling pipelines in real-world settings, two gene annotations, three alignment tools, and eight expression quantification tools were incorporated into the analysis, resulting in 28 combined quantification pipelines. Subsequently, six representative normalization methods were systematically compared **(Supplementary Fig. 27)**. Variations caused by different combinations of analysis tools represent ‘bioinformatics noise’.

Bioinformatics processes introduced variations comparable to those from the experimental processes, and each bioinformatics step also had a greater impact on the subtle differential expression measurement **(compare Fig. 4a with Fig. 3c and Fig. 4b with Fig. 3d**). We quantified the relative contribution of annotation, alignment, quantification, and normalization, to variations using PVCA analysis based on the absolute expression data for different samples from all combined pipelines. For the Quartet samples, different bioinformatics steps collectively introduced significantly greater variations than the intrinsic biological differences (75.1% vs. 5.6%). Normalization methods were the primary source of variations, followed by quantification tools, alignment tools, and gene annotation types (**Fig. 4a**). However, MAQC samples revealed smaller variations introduced from different bioinformatics steps than their biological differences (34.0% vs. 56.7%) (**Fig. 4b**).

**Fig. 4.**
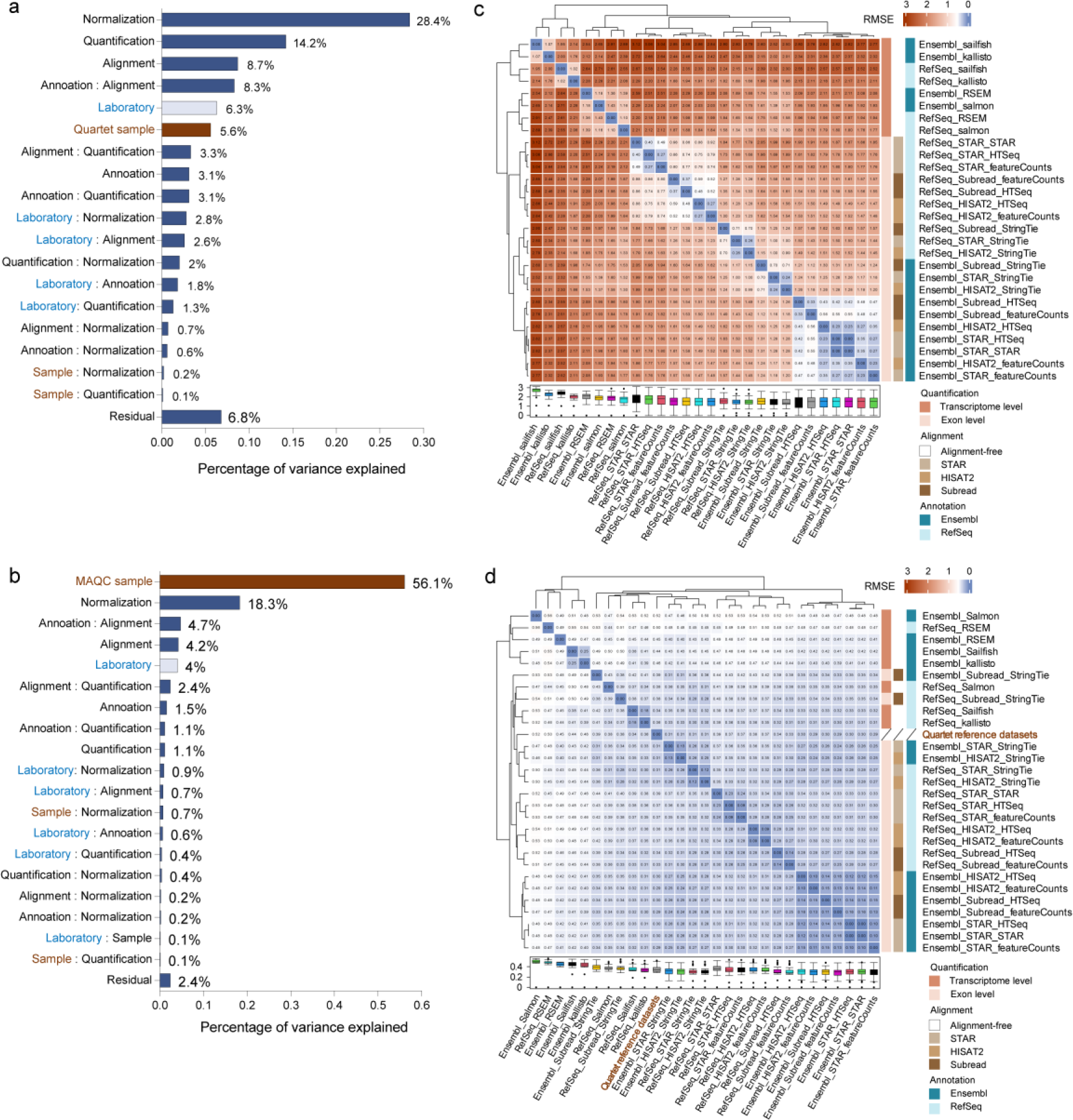
Sources of variation from the bioinformatics process. **(a)** Principal variance component analysis quantifies the proportion of variance explained by each data analysis step for Quartet samples, **(b)** and MAQC samples. **(c)** Heatmap and hierarchical clustering of 28 gene quantification pipelines based on the RMSE at absolute expression levels, **(d)** and relative expression levels. RMSE, Root Mean Square Error.

Noticeably, the calculation of relative expression could help reduce these variations, as indicated by the increased consistency in relative expression levels across different analysis pipelines when compared to absolute expression (**Fig. 4c–d**). Furthermore, the contribution of each bioinformatics step to variations in relative expression levels decreased significantly **(compare Supplementary Fig. 28 with Fig. 4**), suggesting that the relative expression calculations could correct for the influence of different analysis tools. However, similar to the experimental process, 28.4% of the variations from the bioinformatics process remained for the Quartet samples, suggesting inherent performance differences among various analysis tools.

### Best practices for experimental designs

To assess whether experimental factors are related to overall performance, we assessed the accuracy of 42 experimental processes based on the reference datasets under uniform analysis pipeline conditions. We observed that laboratories exhibiting high correlation coefficients for relative expression measurements or high MCC values for DEGs detection dispersed across various experimental protocols **(Supplementary Fig. 29**). Therefore, these results indicate that RNA-seq performance is primarily dependent on experimental quality, with the choice of experimental protocols having a relatively minor impact.

We further filtered out RNA-seq data with low experimental quality and utilized the remaining data from 32 laboratories to evaluate each experimental factor with regard to data quality, and accuracy of absolute expression, relative expression, and differential gene expression **(Materials and Methods)**. All performance metrics demonstrated similar patterns in assessing different protocols within each experimental step (**Supplementary Fig. 30)**, and collectively demonstrated that experimental factors predominantly influence absolute expression measurements rather than relative expression and differential gene expression. Specifically, certain experimental factors related to performance were identified (**Fig. 5**). First, the poly(A) selection method exhibited higher SNR values than the rRNA depletion method, which is associated with the latter capturing more lowly expressed non-protein-coding genes. Second, for absolute expression levels, the rRNA depletion method, strandedness, and 100 bp of read length corresponded to higher accuracy, and exonic coverage also exhibited a significantly positive relationship with the accuracy. Third, exonic coverage was also associated with improved accuracy of relative expression or differential gene expression, likely due to more reliable detection of lowly expressed genes. We also observed significant differences in accuracy associated with different choices of some experimental methods, such as library kit, sequencing platform, and reads length, but these findings were derived solely from comparisons with a single reference dataset.

**Fig. 5.**
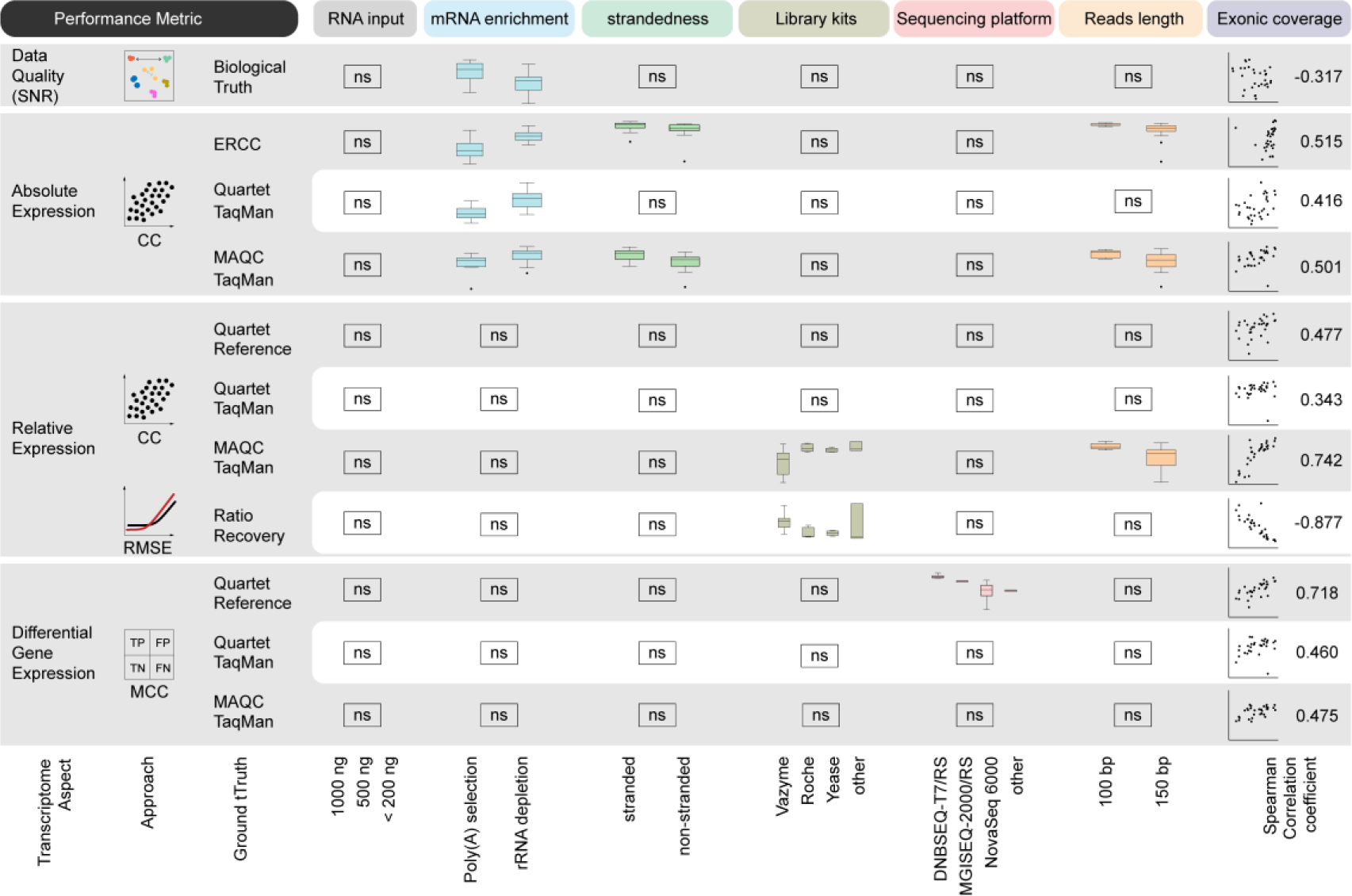
The influence of experimental factors based on different performance metrics. Performance metrics included SNR for data quality, correlation coefficient for accuracy of absolute and relative expression, RMSE for recovery of mixing ratios, and MCC for differential gene expression. The impact of exonic coverage is evaluated by Spearman correlation analyzes. Significance testing was conducted based on normal distribution assumptions using one-way analysis of variance (ANOVA) and paired t-tests, or, in cases where normal distribution was not observed, independent samples were subjected to Kruskal-Wallis test and Mann-Whitney U test. ** indicates a p-value < 0.05. ns, not significant; SNR, Signal-to-Noise Ratio; RMSE, Root Mean Square Error; CC, Correlation Coefficient; MCC, Matthews Correlation Coefficient; TN, True Negative; TP, True Positive; FN, False Negative; FP, False Positive.

### Best practices for bioinformatics designs

To obtain an optimal analysis pipeline for gene-level quantification and differential expression measurements, we sequentially evaluated the performance of 140 combined analysis pipelines with regard to alignment quality, quantification accuracy, normalization effectiveness, low-expression gene filtering efficacy, and accuracy of DEGs identification.

We first evaluated the influence of six alignment approaches combined with two annotations and three alignment tools in terms of sequence alignment and splice junction discovery. In comparison to the RefSeq annotation, the Ensembl consistently resulted in higher uniquely mapping rates and lower multi-mapping rates (**Fig. 6a**). STAR exhibited the highest overall mapping rate as well as uniquely mapping rate. STAR either mapped or discarded the paired reads, avoiding the alignment of unpaired single-end reads (**Fig. 6a**). HISAT2 and Subread had comparable uniquely mapping rates, yet HISAT2 tended to have slightly higher multi-mapping rates in most samples, resulting in higher overall mapping rates. Subread displayed a higher tolerance of accepting mismatch, primarily concentrating in fewer mismatched bases (**Fig. 6b**). Given that Subread did not detect exon-exon junctions, we compared the junctions from STAR and HISAT2. The Ensembl annotation, being more complex, led to the validation of a greater number of junctions (**Fig. 6c and Supplementary Fig. 31)**. For these known junctions, two alignment tools did not exhibit significant differences, whereas HISAT2 identified more completely novel junctions (**Fig. 6c**). Most of novel junctions were not reliable, indicated by significantly decreased number after applying a counts-based threshold **(Supplementary Fig. 32)**. Additionally, we examined the influence of sequencing depth on junction discovery, and observed that even lower sequencing depth was sufficient to detect known junctions, and increasing the sequencing depth further facilitated the identification of more novel junctions **(Supplementary Fig. 33)**.

**Fig. 6.**
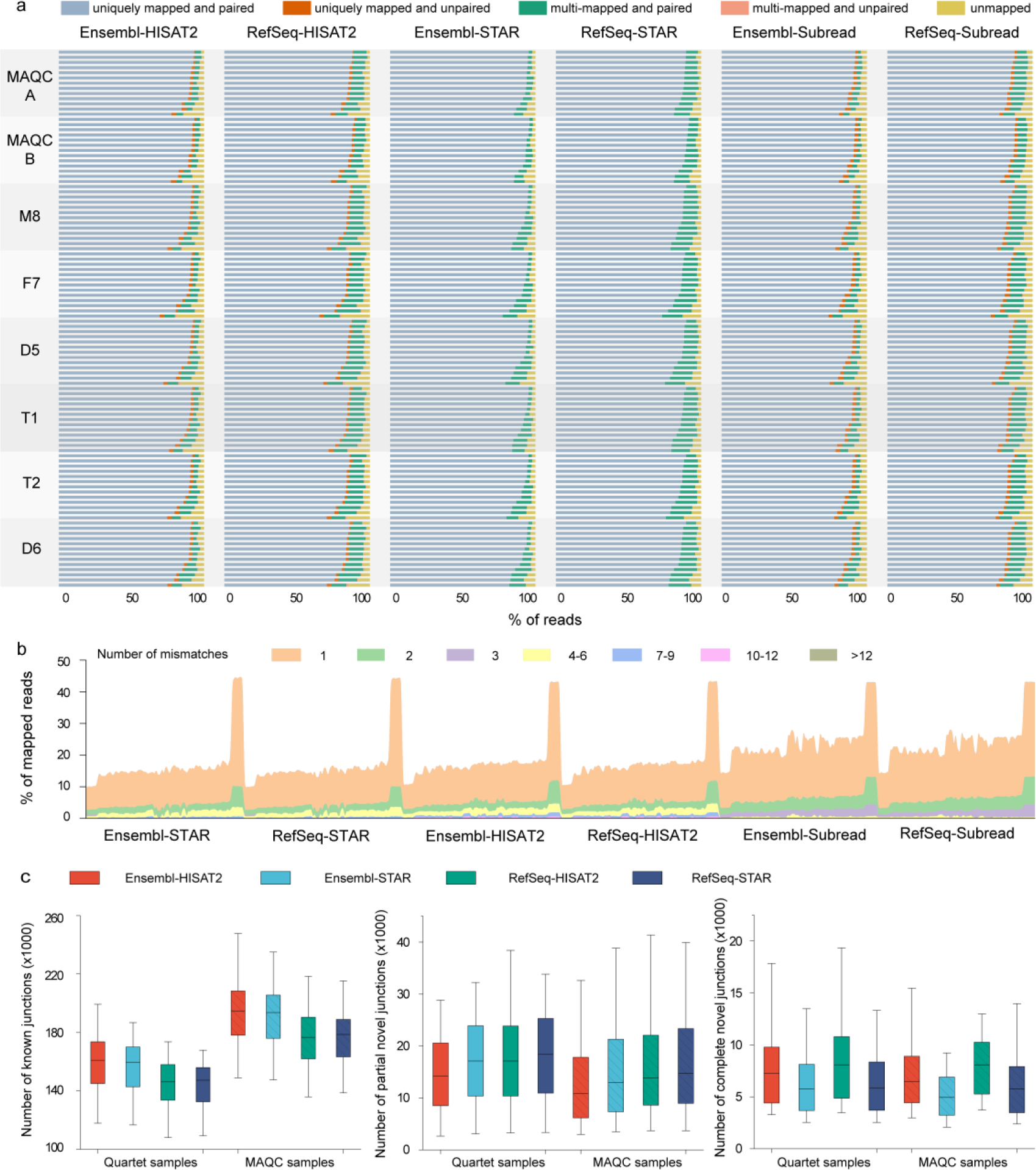
Performance of different alignment schemes. **(a)** Distribution of mapping status of sequenced reads for six combinations of annotation and alignment tools. The 13 benchmark datasets corresponding to each sample are arranged in descending order based on the uniquely mapping rate. **(b)** Distribution of the number of reads with mismatch bases. **(c)** Comparison of known junctions (left), partially novel (middle), and completely novel junctions (right) detected by different alignment approaches in Quartet and MAQC samples. Only junctions supported by at least one reads for all three replicates were included.

We next assessed the performance of 28 gene quantification pipelines, consisting of six alignment approaches and eight quantification tools **(Supplementary Fig. 27)**. These pipelines demonstrated similar clustering patterns at absolute and relative expression levels, primarily divided into two clusters based on quantification principles: exon-level tools (featureCounts, HTSeq, StringTie, and STAR) and transcript-level tools (RSEM, Salmon, kallisto, and Sailfish) (**Fig. 4c–d and Supplementary Fig. 34)**. Gene annotation and alignment tools also contributed to the clustering. In particular, different gene annotations showed a pronounced impact on absolute expression measurement using featureCounts, HTSeq, and STAR, and in relative expression measurement using transcript-level quantification tools. We further examined the impact of different annotations, alignment tools, and quantification tools on accuracy based on three reference datasets, and found that the performance of each step was interdependent. The choice of gene annotation should also consider the quantification tool, as Ensembl annotation exhibited higher or similar accuracy when combined with genome- or transcriptome-alignment quantification tools, whereas RefSeq exhibited higher accuracy when combined with pseudoalignment quantification tools **(Supplementary Fig. 35)**. The impact of different alignment tools was relatively small, but the combination of Subread and StringTie decreased accuracy. **(Supplementary Fig. 36)**. The accuracy also varied among different quantification tools, especially between exon-level and transcript-level quantification tools **(Supplementary Fig. 37)**. Overall, our results, derived from the performance ranking of all quantification pipelines, supported the superior performance of opting for Ensembl gene annotation, any alignment tool, and either featureCounts or HTSeq for quantification **(Supplementary Fig. 38)**.

We converted the raw counts from the 28 quantification pipelines using six normalization methods, followed by an assessment of expression data quality using PCA-based SNR **(Supplementary Fig. 39a)**. Trimmed mean of M values (TMM), and DESeq normalization methods appeared to improve the raw counts most effectively, while upper quartile (UQ) normalization exhibited the poorest improvement **(Supplementary Fig. 39b)**. Then we examined the gene expression distribution for all normalization methods, and found that the median gene expression from DESeq was the highest, followed by TMM, total counts (TC), and UQ, while fragments per kilobase million (FPKM) and transcripts per million (TPM) had similarly low levels **(Supplementary Fig. 40)**.

The setting of low-expression gene filtering conditions may affect the interpretation of differential expression calls **(Supplementary Fig. 23)**. To elucidate the impact of filtering conditions on the performance of differential analysis, we evaluated six filtering methods and various threshold values (0–70%) across five differential analysis tools, utilizing four RNA-seq datasets representing different sequencing depth levels **(Supplementary Fig. 27) (Materials and Methods)**. Across all six filtering methods, elevating the threshold values resulted in an increase in both the DEGs number and the true positive rate (TPR) until they reached their respective peak values, accompanied by a slight yet acceptable decrease in precision **(Supplementary Fig. 41)**. Such threshold effects were observed for five differential analysis tools, including edgeR, DESeq2, limma, and DEGseq, except for EBSeq, which employed stringent internal filtering criteria **(Supplementary Fig. 42)**. Overall, the six filtering methods led to general consistency in terms of the maximum number of DEGs and the highest TPR across data from all laboratories **(Supplementary Fig. 43–44)**. Thus, the key consideration shifts to the determination of optimal threshold value. In the context of small changes in precision, opting for a threshold value corresponding to the highest TPR appears to be an effective approach, but the lack of benchmark datasets for assessing sensitivity or precision presents a challenge in practice. In contrast, calculating the maximum number of DEGs is practical. Although there were slight differences between the thresholds based on the maximum number of DEGs and the highest TPR, especially in the Quartet samples **(Supplementary Fig. 45)**, the resulting TPR values corresponding to these two thresholds were highly consistent **(Supplementary Fig. 46)**.

After applying a series of threshold values to filter low-expression genes, we compared the optimal performance of five differential analysis tools with different choices of quantification pipelines, which contributed to 140 differential analysis pipelines **(Supplementary Fig. 27)**. First, the number of DEGs identified in both the Quartet and MAQC samples was assessed. DEGseq identified the highest number of DEGs, followed by edgeR, limma, and DESeq2, while EBSeq detected the lowest **(Supplementary Fig. 47)**. Compared to other tools, DEGSeq appeared to be more influenced by different choices of quantification pipelines, reflected in a broader range of DEGs numbers. Second, when focusing on the accuracy of DEGs calls, edgeR and DESeq2 consistently outperformed other tools, with DEGSeq and limma slightly lower, and EBSeq being the lowest (**Fig. 7a–b and Supplementary Fig. 48)**. Compared to the Quartet reference datasets, alignment-free quantification tools, especially Sailfish and kallisto, were associated with lower MCC coefficients, regardless of the differential analysis tool used **(Supplementary Fig. 49)**. However, the MAQC samples demonstrated small impact of the different quantification pipelines on each differential analysis tool (**Fig. 7b and Supplementary Fig. 48)**. As another accuracy measure, the area under the receiver operating characteristic curve (AUC) was compared across all differential analysis pipelines, which captured the statistical discrimination capability of the DEGs. The edgeR outperformed the other tools, and DESeq2 also exhibited relatively high AUC values (**Fig. 7c and Supplementary Fig. 46)**.

**Fig. 7.**
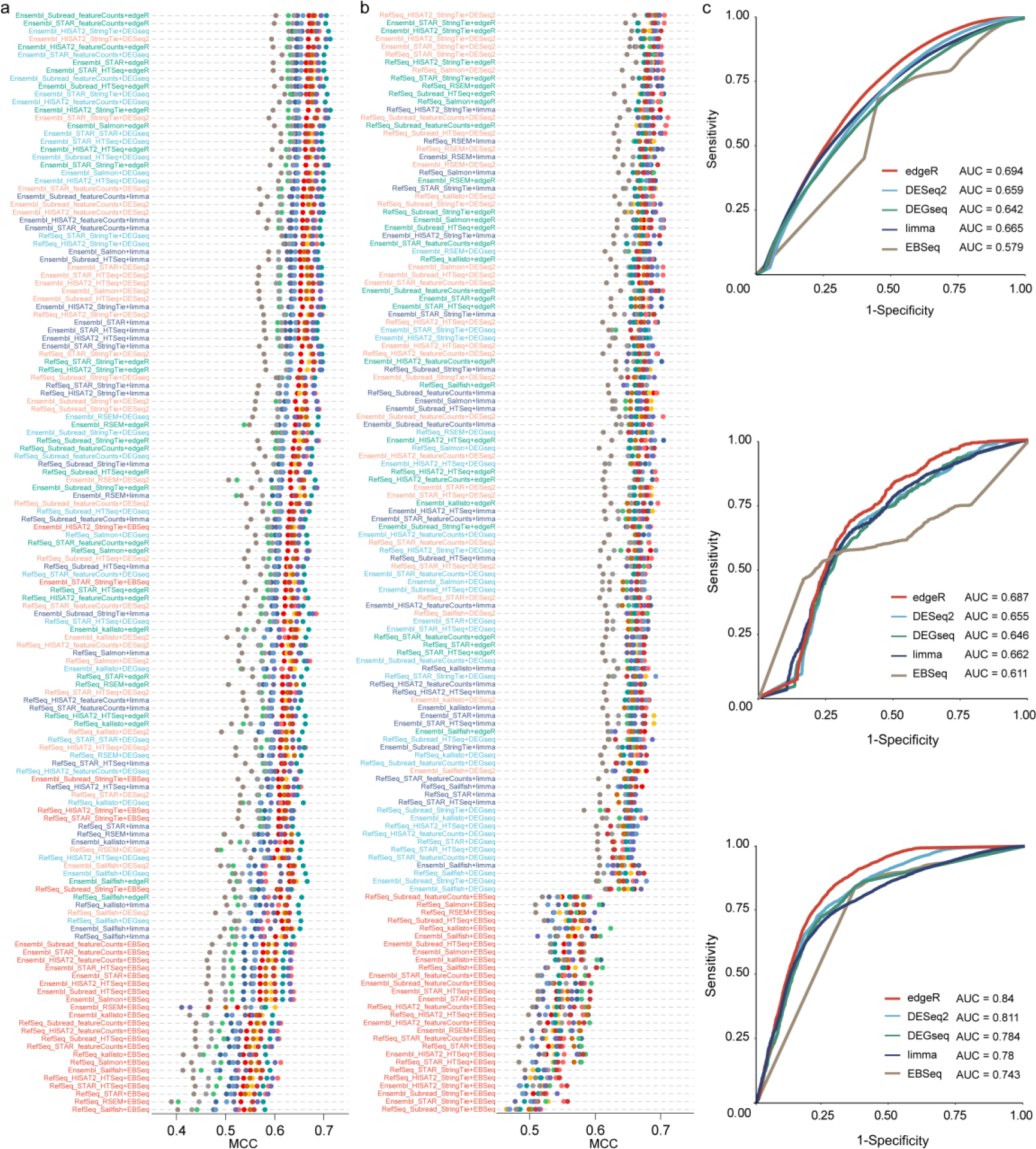
Performance of differential gene expressions analysis tools. **(a)** The Matthews Correlation Coefficients (MCC) was measured based on Quartet reference datasets, and **(b)** MAQC TaqMan dataset. **(c)** ROC analysis of genes in Quartet reference datasets (up), Quartet TaqMan dataset (middle), and MAQC TaqMan dataset (down). For each differential analysis tool, the plot reflects average performance when different annotations, alignment tools, and quantification tools are used for gene expression estimation. The RNA-seq data from lab01 was utilized to calculated the AUC values, and the AUC values for other high-quality benchmark datasets were displayed in **Supplementary Figure 50**. AUC, Area Under the receiver operating characteristic Curve.

## Discussion

As part of the Quartet project, this study represents the most extensive cross-laboratory examination of real-world RNA-seq data and analysis outcomes to date, employing Quartet and MAQC RNA reference materials. Through the systematic assessment of transcriptome data from 45 laboratories and the comparison of over 40 experimental processes and 140 bioinformatics pipelines based on several ‘ground truths’, we attempted to address several questions: (i) the performance of real-world RNA-seq in detecting subtle differential expression; (ii) the sources of inconsistency among laboratories; and (iii) the recommended practices to enhance the accuracy of RNA-seq in practical applications (**Table 1**).

**Table 1.**
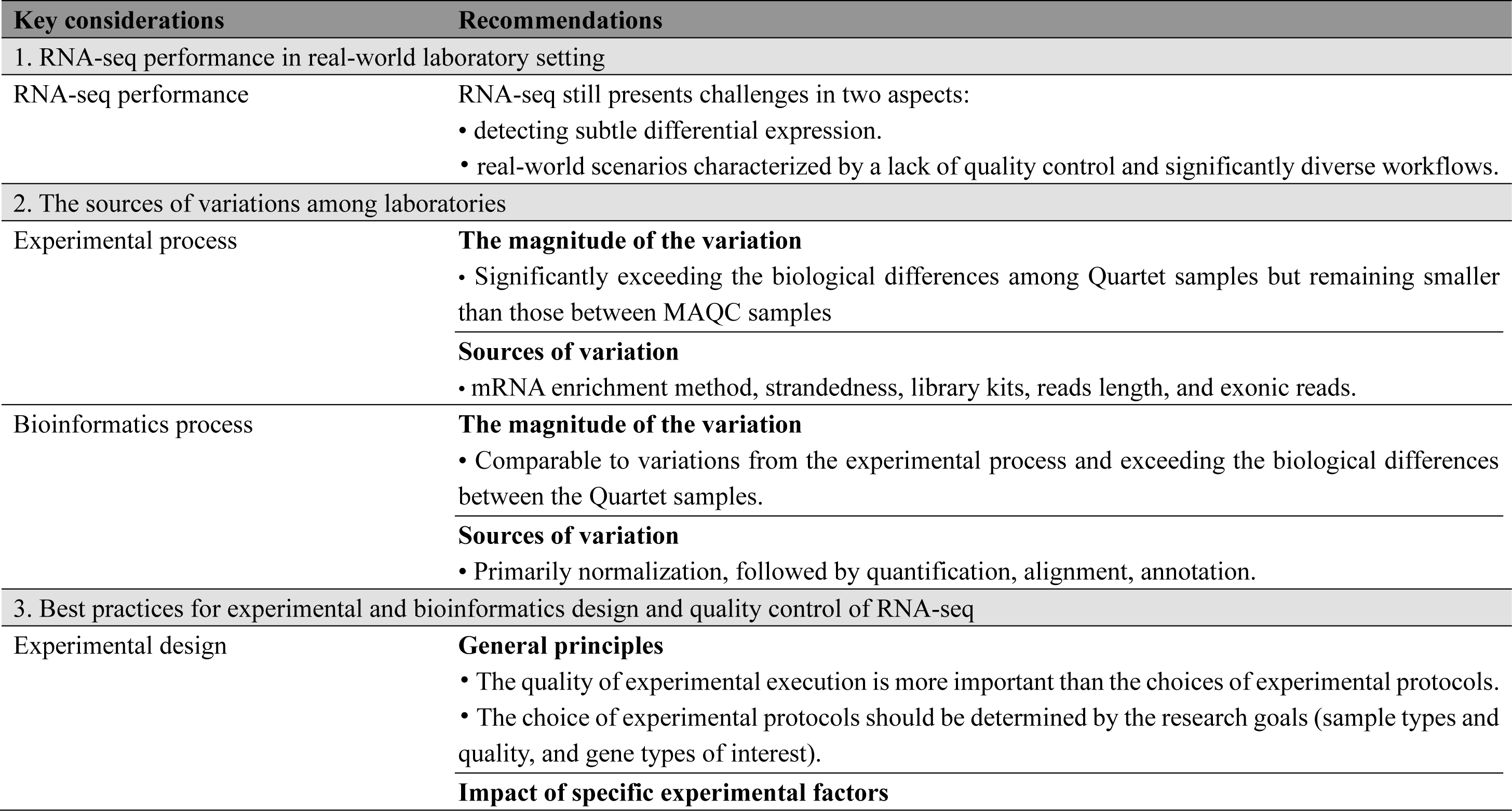
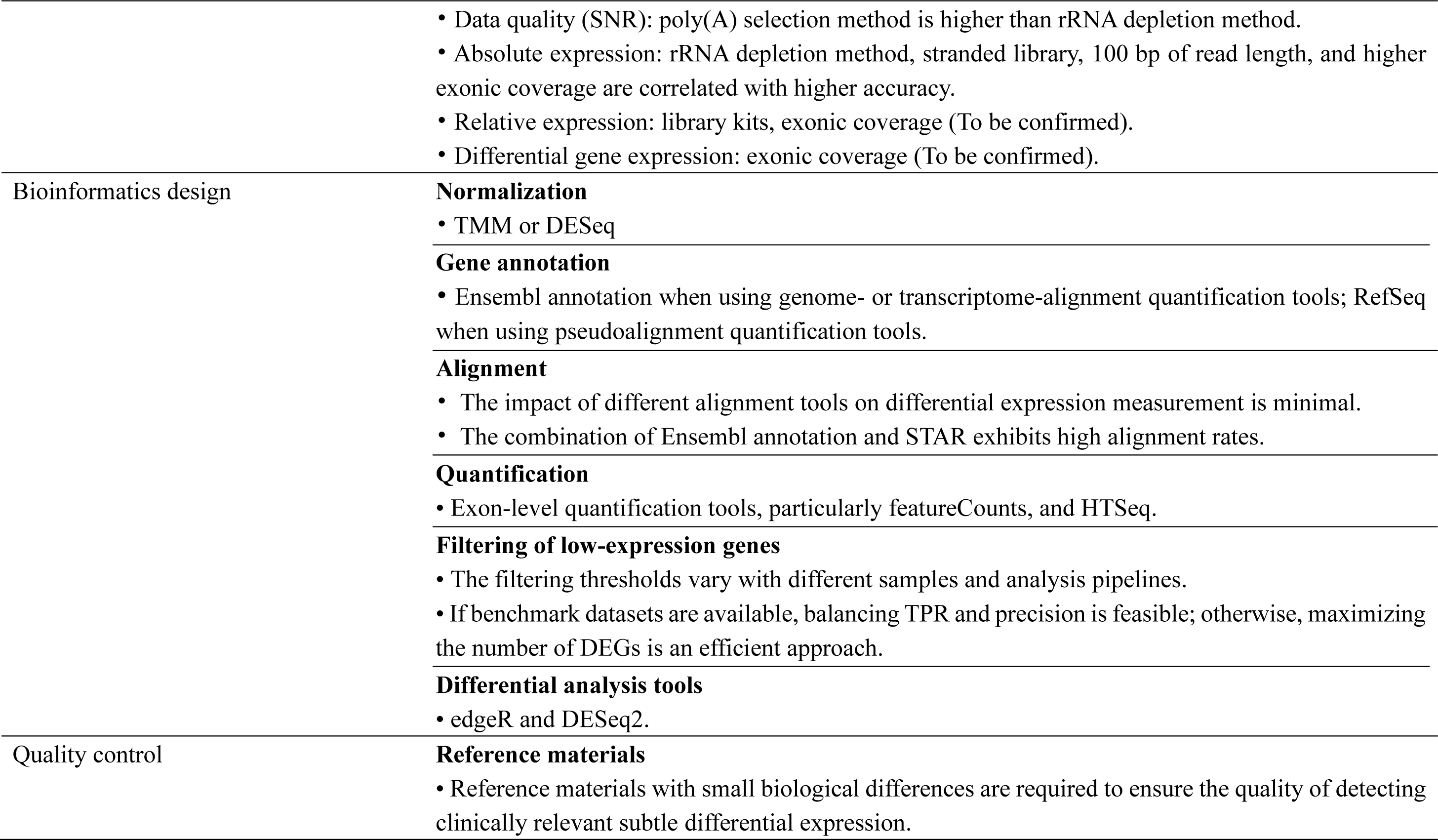
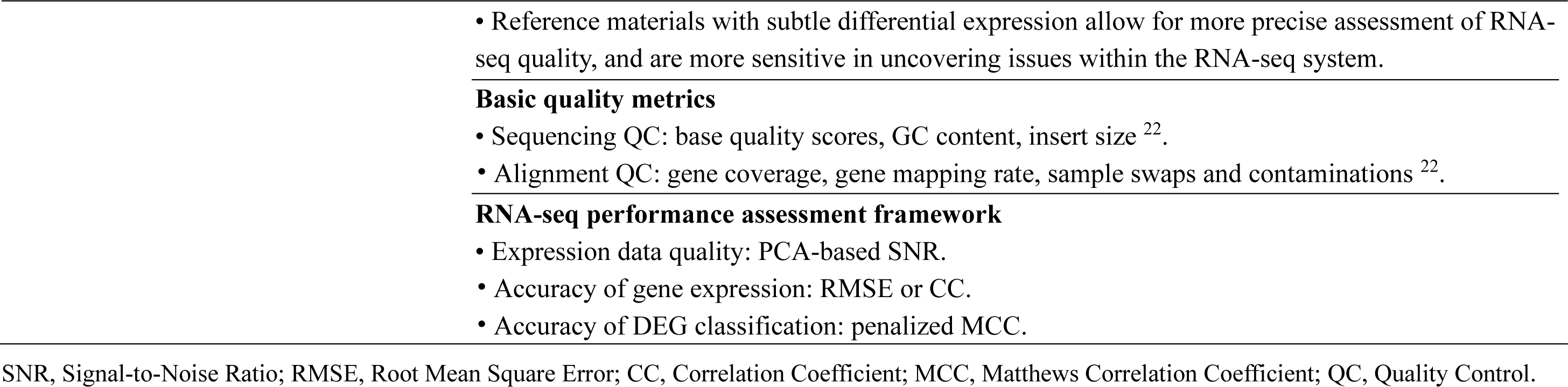
Best practice recommendations.

This study, for the first time, revealed noteworthy real-world inter-laboratory variations in transcriptome profiling performance, especially when detecting subtle differential expression among the Quartet samples. This prompts a reconsideration of the actual performance, which may not be as robust as in previous studies conducted under rigorously controlled RNA-seq workflows ^14, 15, 22^. First, the PCA-based SNR varied significantly across laboratories, with 35.6% (16/45) of expression data considered low quality based on the previously defined cutoff value (SNR = 12) ^32^. Our results also revealed that low data quality correlated with the accuracy of differential analysis calls, highlighting the necessity of quality evaluation prior to downstream analysis within laboratories. Second, absolute expression measurements exhibited substantial inter-laboratory variations as reported in previous studies ^14^. Relative expression also exhibited greater variations when detecting subtle differential expression. Certain laboratories exhibited low consistency with reference datasets and poor recovery of known mixing ratios between mixed samples, which were primarily due to inadequate restoration of inter-sample biological differences in low-quality data or erroneous detection of low-expression genes. Third, the number of DEGs varied widely, and the accuracy metrics for DEG calls demonstrated a broad range across laboratories, even when focusing on protein-coding genes. Differences in data quality, filtering conditions for low-expression genes, differential analysis tools, and the cutoff setting for DEGs classification collectively contribute to such variations, which appear to be more significant than the differences in DEGs calling performance across platforms, sites, and analysis tools previously reported ^14, 15, 21^. Therefore, our results underscore the fact that real-world RNA-seq performance may not fully meet the clinical diagnostic demands, requiring ongoing quality improvement specifically toward subtle differential expression.

The greater inter-laboratory variations in detecting subtle differential expression among the Quartet samples prompted to investigate the sources of variations from diverse RNA-seq workflows, which compensated for previous studies that exclusively focused on the sources of variation under identical protocols and analysis pipelines ^16, 22^. We observed that the technical factors in experimental and bioinformatics processes contributed to a higher proportion of variations in the Quartet samples (89.4% and 75.1%) compared to MAQC samples (45.3% and 34%). While relative expression could correct for the influence of these factors to some extent, they still contributed to a higher proportion of variations under small biological difference conditions (48.2% and 10.9% vs. 12.6% and 1.7%). To be specific, in the experimental process, we identified factors affecting absolute expression quantification, including mRNA enrichment methods, strandedness, library kits, read length, and exonic coverage. In the bioinformatics process, normalization step is the primary source of variations, followed by quantification, alignment, and annotation. These factors have been individually studied ^15, 33–37^, and in contrast, our study revealed the magnitude of their impact in real-world laboratory settings, providing clarity on the priority of technical factors to consider when designing RNA-seq systems.

The experimental design is generally considered to be centered around addressing the biological questions of interest ^38^ (**Table 1**). Experimental factors contribute to deviations in absolute expression measurement, limiting its application ^14^. Given the prominent application of differential expression analysis for potential clinical usage, we particularly focused on the influence of these technical factors in terms of relative expression and differential gene expression measurements. Our results revealed the quality of the experimental execution is the primary determinants of accuracy, not these experimental factors. The impact of low-quality experiments far outweighed that of different experimental protocols on accuracy, and the varied choices within each experimental method have not demonstrated significant differences in differential analysis performance. Nevertheless, it’s important to note that different experimental methods capture distinct transcriptomic features. For example, rRNA depletion method detects more non-coding RNAs and pseudogenes compared to poly(A) selection method ^15, 33^. Stranded and non-stranded libraries mainly contributed to the differential expression of pseudogenes and antisense genes, and stranded RNA-seq enables the accurate quantification of approximately 20% of overlapping genes transcribed from the opposite strands ^34^. Therefore, the choice of experimental protocols would be primarily driven by (i) sample type and quality, such as the extent of RNA degradation ^17^, and (ii) research objectives, which may involve non-coding RNAs, pseudogenes, antisense genes, as well as novel transcripts and alternative splicing events ^33, 34, 39^.

The bioinformatics design, centered on the choice of optimal analysis tools, requires equal attention, as the variations from the bioinformatics processes are comparably significant as those from the experimental processes (**Table 1**). This study assessed different normalization methods from the data quality aspects and found that TMM and DESeq significantly improved the quality of expression data, agreeing with conclusions drawn from previous studies ^40^. For each step of the differential expression analysis, we found that the performance of any analysis tool is not constant but depends on the other tools used in combination with it. Nevertheless, this study provided the optimal bioinformatics design through an evaluation of arbitrary combinations of analysis tools. First, choose Ensembl annotation when using genome- or transcriptome-alignment quantification tools, and choose RefSeq when using pseudoalignment quantification tools. Second, the impact of different alignment tools is relatively small, but previous studies have indicated that varying genome complexity should be considered when making choices ^41^. Third, for quantification, choose tools operating at the exon level, particularly featureCounts and HTSeq. Fourth, the threshold for filtering low-expression genes is not fixed but varies with different samples and analysis tools ^42^. Choosing the threshold based on the maximum number of DEGs is practical. Finally, edgeR or DESeq2 is preferred for conducting differential gene expression analysis.

This study significantly advances the understanding of the role of reference materials in quality control applications by utilizing Quartet and MAQC reference materials in parallel (**Table 1**). Overall, the assessment based on these two reference materials demonstrated common patterns in multiple aspects of the transcriptome across laboratories. Notably, each of the two reference materials has significantly enhanced the reliability and distinctiveness of the assessment and exploration of RNA-seq data. On the one hand, the Quartet samples enabled the assessment in subtle differential expression levels and demonstrated advantages in the performance assessment for different laboratories and various analysis pipelines, underscoring the need for a shift in RNA-seq benchmarking toward subtle differential expression levels. First, Quartet samples with large-scale reference datasets enabled a more precise and comprehensive assessment of the RNA-seq performance. The performance metrics exhibited a broader range than those from the MAQC samples in terms of SNR values for assessing data quality, correlation coefficients for assessing gene expression accuracy, and MCC coefficients for evaluating the accuracy of DEG calls. This implies a higher discriminative ability for discovering performance differences among different batches, protocols, sites, and analysis tools. Second, Quartet samples allowed for a more sensitive uncovering of technical noise. In the context of subtle biological differences among the Quartet samples, the variations introduced by experimental and bioinformatics factors become more pronounced. Third, the Quartet reference datasets revealed no systemic differences with the RNA-seq data at both absolute and relative expression levels. Methodological differences between RNA-seq and TaqMan RT-qPCR have previously limited gene expression assessments concerning correlation analyzes ^14^, which are considered to have limitations in representing consistency ^43^. The Quartet reference datasets showed a lower RMSE with RNA-seq data compared to TaqMan datasets, allowing for a direct comparison of the quantitative values of gene expression. On the other hand, the MAQC samples established connections with previous milestone studies, contributing to a deeper understanding of real-world RNA-seq performance based on these traditional RNA reference materials in the community. Moreover, a large-scale TaqMan RT-qPCR dataset for the MAQC samples ensures an unbiased performance assessment, effectively complementing the Quartet reference datasets originated from the Ensembl-HISAT2-StringTie pipeline that may introduce biases especially when assessing diverse RNA-seq analysis pipelines ^32^.

In summary, this study unveils significant inter-laboratory variations in real-world transcriptome profiling when detecting subtle differential expression, especially with respect to data quality, absolute expression, and differential gene expression. The investigation of the sources of inter-laboratory variations at both experimental and bioinformatics aspects has highlighted key points for the development and optimization of RNA-seq methods. This study provided best practice recommendations regarding the experimental and bioinformatics design and quality control of RNA-seq (**Table 1**). These will aid researchers in accurately identifying subtle changes in disease conditions, accelerating the transition of RNA-seq into a diagnostic tool. Furthermore, these data can also be used to address other aspects of transcriptome profiling, including alternative splicing, gene fusion, RNA editing, and RNA variations.

## Materials and Methods

### RNA Reference samples preparation

Four Quartet RNA reference materials (M8, F7, D6, D5) were used ^32^, and External RNA Control Consortium (ERCC) spike-in transcripts were added to M8 and D6 samples at manufacturer recommended amounts (4456740, Thermo Fisher Scientific) ^13^. Samples T1 and T2 represent mixtures of samples M8 and D6 at the defined ratios of 3:1 and 1:3, respectively, and thus hold ‘built-in truths’ of sample mixing ratios. Universal Human Reference RNA (740000, Agilent Technologies) and Human Brain Reference RNA (QS0611, Thermo Fisher Scientific) were used, which were labeled as MAQC samples A and B by MAQC Consortium ^12^. MAQC B sample was paired with MAQC A sample as a control sample for differential analysis, while Quartet D6 sample served as a control sample for differential analysis of sample M8, F7, D5, T1, and T2. Based on these reference materials, three technical replicates were prepared for 8 RNA samples, resulting in a total of 24 RNA samples (**Fig. 1a**). All the samples dispensed as 8 μL aliquots into 200 μL thin-wall polypropylene PCR tubes with a concentration of 200 ng/μL and stored at -80 ℃.

### RNA-seq workflow

The samples were transported to each laboratory on dry ice, and the ERCC reference sequences and gene annotation files were provided with the names of the 92 ERCC genes modified to ‘SPIKEIN’ followed by the corresponding identifier. Laboratories conducted the experiments and data analysis following their routine procedures. To accurately capture batch effects within the laboratories, the sample grouping information was provided to the laboratories after they submitted the sample quality results, raw FASTQ files, and quantification results at the gene and transcript levels. Subsequently, laboratories were required to submit differential analysis results at gene and transcript level, and alternative splicing results.

### TaqMan RT-qPCR

Primers and TaqMan probes were designed for 91 genes based on the RNA sequences. Among them, *C1ORF43* was selected as the reference gene for the PCR method. Primers and probes were synthesized by Sangon Biotech, and the sequences are shown in **Supplementary Table 5**. Before proceeding with the bulk qPCR experiments, we designed two sets of primers and probes for the reference gene and the target gene (*CD180*) to verify the acceptable impact of primer and probe selection on the results. Then the amplification efficiency of the primers and probes was confirmed to meet the requirements by performing gradient dilution experiments with the samples. The results of the *CD180* gene were used for inter-batch quality control for qPCR experiments.

Five µg of each Quartet RNA sample was reverse transcribed using the PrimeScript™ RT reagent Kit (RR037A, TaKaRa) in a 50 µl reaction. This reaction mixture was incubated at 37 °C for 15 minutes, then for 5 seconds at 85 °C and finally for termination at 4 °C. cDNA obtained in the previous step was used as template for qPCR. qPCR reactions were run in 96-well plates, the qPCR reactions were carried out using Premix Ex Taq™ (RR390A, TaKaRa) containing 2 μL of cDNA, 0.4 μL of each forward and reverse primers, 0.8 μL of TaqMan probes in a 20 μL final volume reaction. The qPCR was performed on an Applied Biosystems 7500 Real-Time PCR System using the following cycling conditions: 30 seconds at 95 °C followed by 45 cycles of 5 seconds at 95 °C and 34 seconds at 56 °C. Three replicates per sample per gene were conducted for eliminating random variations.

Comparative Ct method (delta delta Ct method) was used to calculate the fold differences for the three sample pairs (M8/D6, F7/D6 and D5/D6) with housekeeping gene *C1ORF43* as endogenous control. For the RT-qPCR data, a gene is classified as differentially expressed gene (DEG) when the student’s t-test p-value < 0.05 and fold change ≥ 2 or ≤ 0.5.

TaqMan data for MAQC samples A and B were obtained through the Gene Expression Omnibus database (accession number GSE5350), which was processed as above. Undetectable CT values (CT>35 or CT=0) were removed prior to normalization. The differential gene analysis was performed as previous study, with gene *POLR2A* serving as endogenous control ^15^.

### Relative expression calculation

Relative expression data were obtained within each laboratory on a gene-by-gene basis. Specifically, relative expressions were calculated based on log2FPKM values. For each gene, the mean of expression profiles of replicates of reference sample(s) (for example, D6) was first calculated and then were subtracted from the log2FPKM values of that gene in other samples.

### RNA-seq performance metrics

The PCA-based SNR was used to assess the data quality at the gene expression level, which reflected the ability of data to distinguish the intrinsic biological differences among different sample groups from technical noises present in replicates. The calculation method of PCA-based SNR as shown in the previous study ^23^. Genes with at least one reads in all selected samples were included for PCA analysis. The Pearson correlation coefficient was used to evaluate the consistency between the observed absolute or relative expression and the ground truth. The RMSE was used to measure the difference between RNA-seq data and Quartet reference datasets and TaqMan datasets. The MCC were used to measure consistency of DEGs detected from a dataset for a given pair of samples with those from the reference datasets. The true positives, true negatives, false positives, and false negatives were judged as shown in Fig. 1b. Then MCC was calculated as follow:

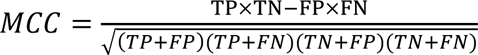

Mixing M8 and D6 into T1 and T2 samples allows for complementary assessment of the accuracy and reproducibility of RNA-seq. The fold changes between M8/D6, T1/D6, and T2/D6 comparisons should adhere to the following equation. A nonlinear robust fit (nlrob) was performed for RNA-seq data from laboratories, and the fitted curves were compared to expected curves (**Fig. 1b**). Then, the RMSE between the observed fold changes and the expected fold changes from the following equation for T1 or T2 versus D6 were calculated.

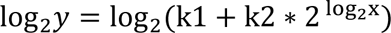

where y represents the expected fold change for T1 or T2 versus D6 and x represents the fold change for M8 versus D6. The correction z of the known mixing coefficients k1 = z/(z+3) and k2 = 3z/(3z+1) arising out of different ratios of mRNA versus total RNA in the samples M8 and D6 has been determined by RT-qPCR assay. In brief, 10 genes with a broad range of fold change were tested using RT-qPCR, and average z values from 10 RT-qPCR results were calculated for samples T1 and T2. The obtained z values were 0.974 ± 0.06 for T1 and 0.949 ± 0.09 for T2. Then, the z values obtained from the top ten laboratories’ RNA-seq data, capable of recovering of mixed ratios, are 0.965±0.024 and 0.941 ± 0.026 for sample T1 and T2, which further validate the correction values. Finally, the z values from RT-qPCR assays were used.

### Alignment and gene quantification

To analyze the sources of variation from the experimental process, we employed the same analysis pipeline for raw FASTQ data from all laboratories. Preliminary processing of raw reads was performed using fastp (v.0.23.2) to remove adapter sequences ^44^. Sequences were aligned to the GRCh38 genome assembly (https://ftp.ensembl.org/pub/release-109/fasta/homo_sapiens/dna/Homo_sapiens.GRCh38.dna.primary_assembly.fa.gz) using STAR (v.2.7.10b) ^45^ with Ensembl annotation release-109 (https://ftp.ensembl.org/pub/release-109/gtf/homo_sapiens/Homo_sapiens.GRCh38.109.gtf.gz). Gene quantification was conducted using StringTie v2.2.1 ^46^. The log2 transformation was then performed based on Fragments Per Kilobase of transcript per Million mapped reads (FPKM) values. To avoid infinite values, a value of 0.01 was added to the FPKM value of each gene before log2 transformation.

Quality control analysis of sequencing data at pre-alignment and post-alignment level was conducted using FastQC (v.0.11.558), Qualimap (v.2.0.060) ^47^, and MultiQC (v.1.8)^48^.

### Filtering of low-quality data

To avoid the impact of low-quality experiments on the examination of experimental methods in terms of various performance metrics, including data quality and accuracy of gene expression and differential gene expression, we selected RNA-seq data from 31 laboratories using two criteria: (i) SNR value greater than 20 after applying the uniform analysis pipeline and (ii) the difference less than 6 between SNR17 and SNR18 values.

### Bioinformatics Pipelines Benchmark Protocols

#### Benchmark datasets

High quality data from laboratories was selected for benchmark study. The benchmark datasets were selected based on three criteria. Firstly, data displaying high duplication rate, abnormal GC distribution, abnormal sequence length distribution, uneven nucleotide composition, and low base quality was excluded based on basic sequencing quality. Subsequently, data with a SNR value below 20 was filtered out. Third, the absence of contamination between samples was required based on ERCC spike-in evaluation. Furthermore, data derived from diverse RNA-seq protocols was required to reduce bias in the benchmark study.

#### Gene annotation

Two human gene annotations were included as the guiding reference for alignment and quantification tasks in this study, including the Ensembl release-109 annotation (https://ftp.ensembl.org/pub/release-109/gtf/homo_sapiens/Homo_sapiens.GRCh38.109.gtf.gz) and the recent RefSeq annotation (2023-03-21) (https://ftp.ncbi.nlm.nih.gov/refseq/H_sapiens/annotation/GRCh38_latest/refseq_identifiers/GRCh38_latest_genomic.gtf.gz). All these annotations were generated based on the human reference genome GRCh38. The gene annotation files were used in conjunction with reference genome or transcriptome files from the corresponding database.

#### RNA-seq analysis tools

The list of RNA-seq tools, versions, and the command line used in the analysis are listed in **Supplementary Table 6**. We integrated alignment tools including STAR (v.2.7.10b) ^45^, HISAT2 (v.2.2.1) ^49^, and Subread (v.2.0.3) ^50^, genome-alignment quantification tools like featureCounts (v.2.0.3) ^51^, HTSeq (v.2.0.2) ^52^, and StringTie (v.2.2.1 ^46^, transcriptome-alignment quantification tools, RSEM (v.1.3.1) ^53^, as well as alignment-free quantification tools, including Kallisto (v.0.48.0) ^54^, Salmon (v.1.10.1) ^55^, and Sailfish (v.0.9.0) ^56^. For differential analysis, edgeR (v.3.42.4) ^57^, limma (v.3.56.2) ^58^, DESeq2 (v.1.40.2) ^59^, DEGseq (v.1.54.0) ^60^, and EBSeq (v.1.40.0) ^61^ were included and compared. The mapping information of each mapping tool was evaluated using Samtools flagstat and stats function ^62^. The number of mismatches was detected using the NM tag. The junctions were extracted from Bam files using ‘junction_annotation.py’ in RSeQC package (v.5.0.1) ^63^. Transcript-level reads counts from Sailfish and kallisto were transformed to gene-level counts using tximport package (v.1.28.0) ^64^.

#### Normalization methods

We consider six normalization methods: total counts (TC), fragments per kilobase million (FPKM), transcripts per million (TPM), trimmed mean of M values (TMM), upper quartile (UQ) normalizations, and normalization method used by DESeq2 (v.1.40.2). TC also known as CPM (Counts Per Million), corrects for library size (expressed in million counts) so that each count is expressed as a proportion of the total number reads in the sample. FPKM and TPM are similar methods that correct for both library size and gene length, but TPM divides counts by gene length first and then by total number of transcripts in the sample, resulting in each normalized sample having the same number of total counts. The TMM approach is to choose a sample as a reference sample and the others as test samples. Under the hypothesis that the majority of genes are not DEGs, a scaling factor is calculated to adjust for each test sample after excluding highly expressed genes and genes with high log ratios between the test and the reference sample ^65^. The TMM normalization method is implemented in the edgeR package (v.3.42.4) by means of the calcNormFactors function ^57^. UQ normalization first removes all zero-count genes and calculates a scaling factor for each sample to match the 75% quantile of the counts in all the samples ^66^. UQ normalization was performed using the uqua function in package NOISeq (v.2.44.0) ^67^. DESeq normalization method is also based on the hypothesis that most genes are not DEGs. The scaling factor for a given sample is computed as the median of the ratio of the read count and the geometric mean across all samples for each gene ^68^. Raw counts were normalized using the estimateSizeFactors() and sizeFactors() functions in the DESeq package (v.1.40.2).

#### Filtering Conditions for Low-Expression Genes

Data from four different laboratories, with varying sequencing depth levels ranging from low to high, were utilized to validate the optimal filtering methods and thresholds. We calculate the maximum (max), median, and sum of raw read counts and CPM for each gene from the replicated samples, resulting in six different combined filtering methods. Using each filtering method, we applied a series of thresholds, ranging from low to high, to filter out up to 70% of lowly expressed genes. To facilitate comparison of different filtering methods, the real threshold values were transformed into percentile-based thresholds. We next examined the performance of different differential analysis tools after applying different filtering conditions. The true positive rate (TPR) measures the proportion of DEGs that are accurately detected as positive by the differential analysis tools. Precision measures the proportion of the detected DEGs made are correct (true positives).

### Statistical analysis

All statistical analyses were performed using R statistical packages (v.4.3.0) and python (v.3.10.10). PCA was conducted with the univariance scaling, using the prcomp (v.3.6.2) function. Principal variance component analysis (PVCA) was performed by pvca package (v.1.40.0) to quantifies the proportion of variance explained by each influencing factor ^69^.

## Data availability

The raw sequence data reported in this paper have been deposited in the Genome Sequence Archive (Genomics, Proteomics & Bioinformatics 2021) in National Genomics Data Center (Nucleic Acids Res 2022), China National Center for Bioinformation / Beijing Institute of Genomics, Chinese Academy of Sciences (GSA-Human: HRA005937) that are publicly accessible at https://ngdc.cncb.ac.cn/gsa-human.

## Acknowledgements

This study was supported by National Key R&D Project of China (2023YFC3402500). We thank the 45 laboratories for performing RNA-seq and for returning the raw data and analysis results on time.

