## Supplementary Materials for "Toward Best Practice in Identifying Subtle Differential Expression with RNA-seq: A Real-World Multi-Center Benchmarking Study Using Quartet and MAQC Reference Materials"

### 1. Supplementary Figures

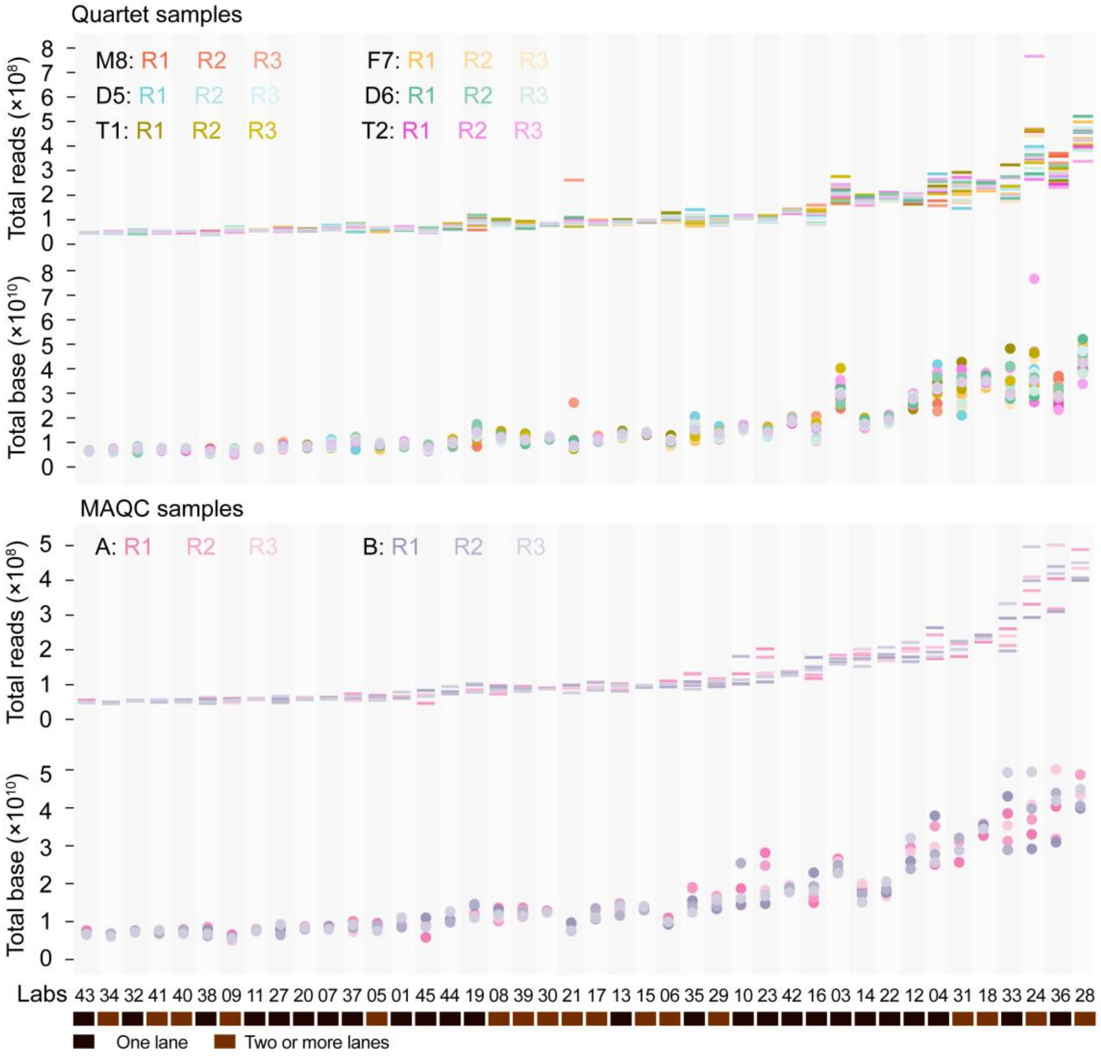

**Supplementary Figure 1. Distribution of sequencing depth of RNA-seq data.** Total reads and total bases of Quartet (up) and MAQC samples (down) were compared, where higher sequencing depth is associated with greater inter-sample variations.

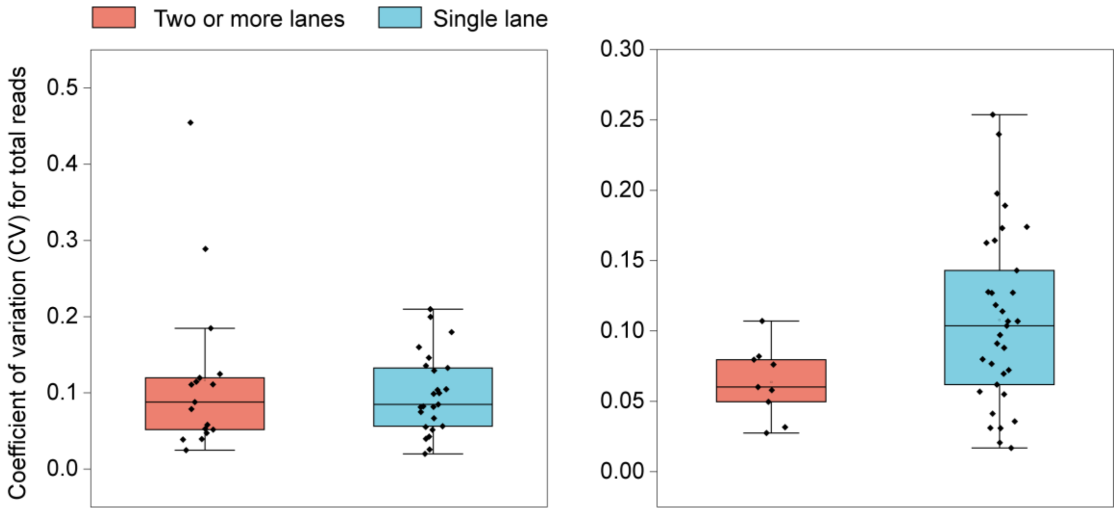

**Supplementary Figure 2. The influence of different lanes on the sequencing depth.** Assigning the 24 libraries into different lanes within the same laboratory did not lead to a higher coefficient of variation (CV) for total reads across samples when compared to assigning the 24 libraries into a single lane.

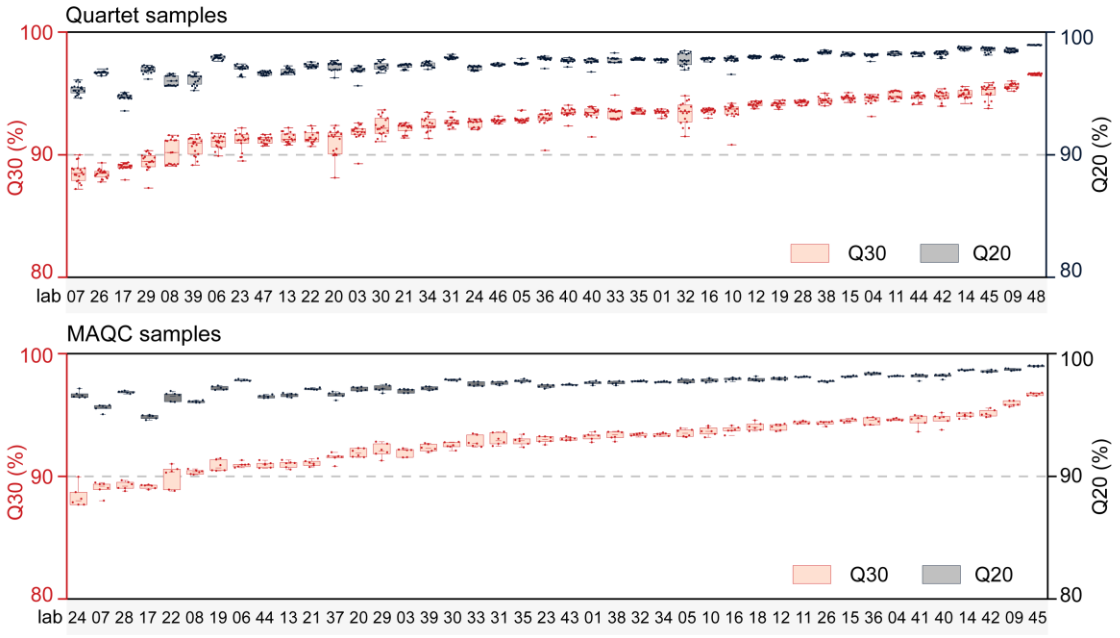

**Supplementary Figure 3. Quality values of sequencing reads of RNA-seq data.** Q30 for Quartet (up) and MAQC samples (down) was calculated as the percentage of bases with a quality score of 30 or higher, indicating the base call accuracy is 99.9%. Q20 was calculated as the percentage of bases with a quality score of 20 or higher, indicating the base call accuracy is 99 %.

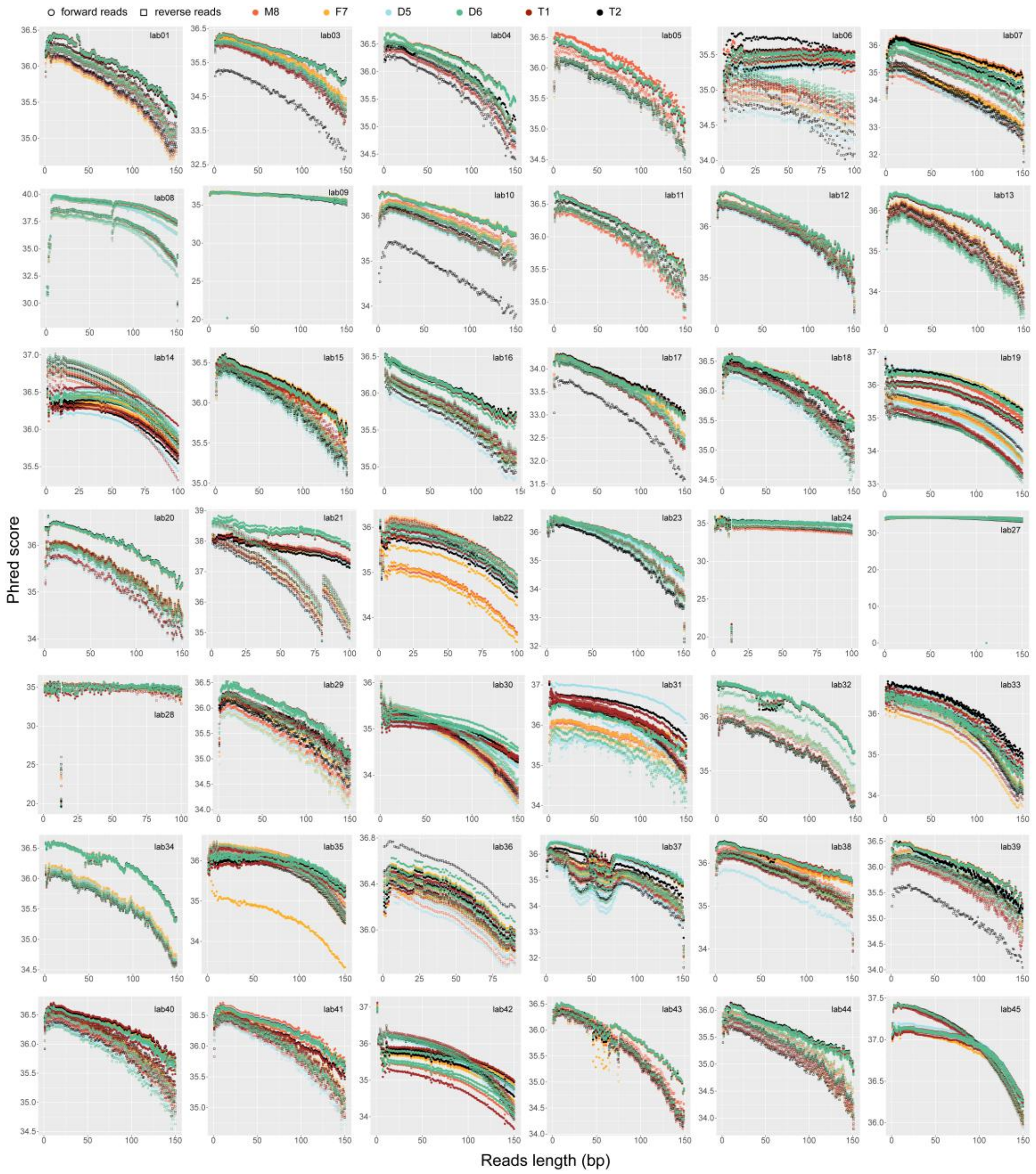

**Supplementary Figure 4. Base quality values of representative sequencing reads for Quartet samples.** The base quality distribution in the first about 10 bases was biased in most laboratories. The quality of reverse reads was generally lower than that of forward reads in most laboratories. Different colors represent different samples, with squares and circles representing forward and reverse reads, respectively.

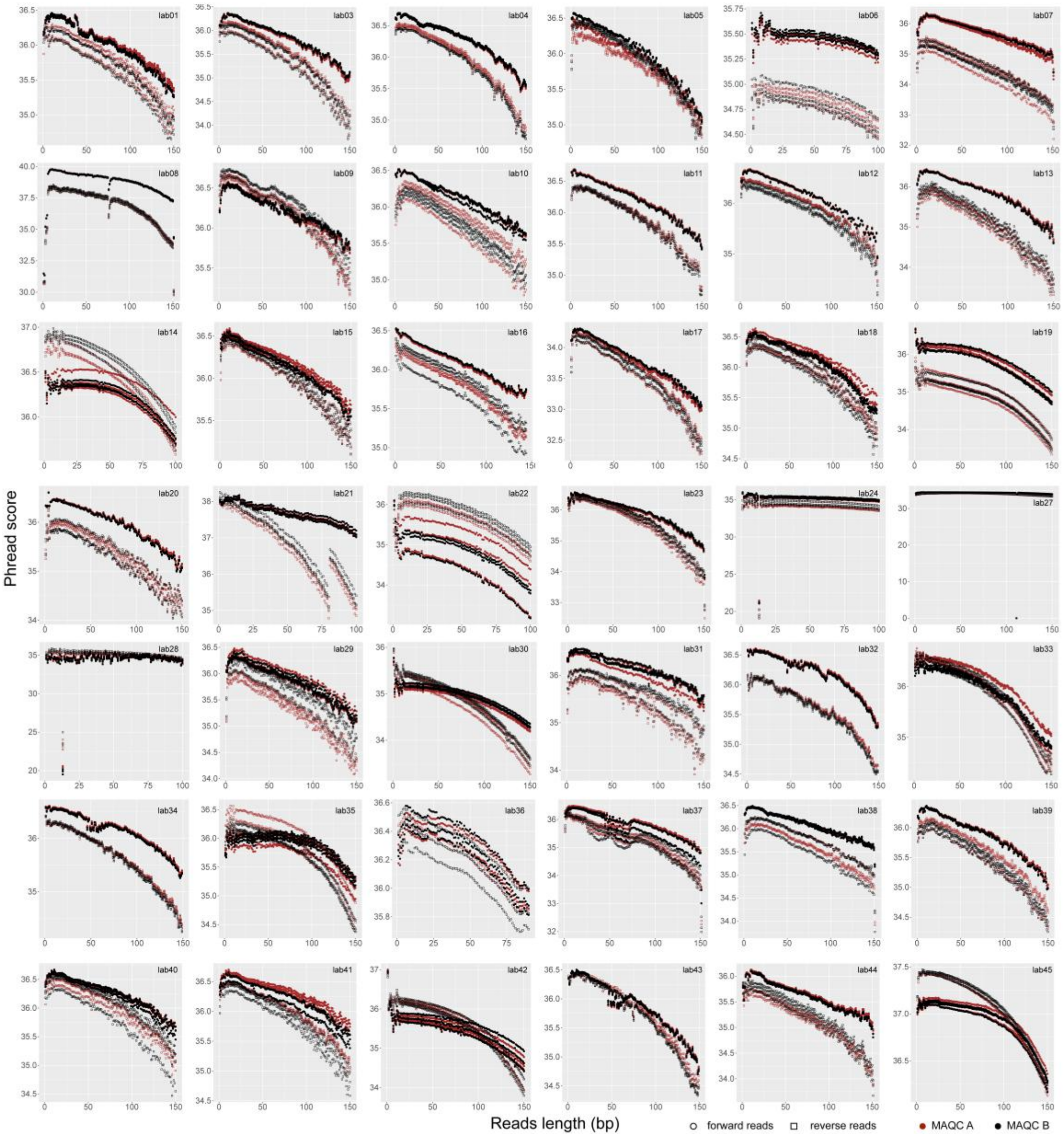

**Supplementary Figure 5. Base quality values of representative sequence reads for MAQC samples.** Different colors represent different samples, with squares and circles representing forward and reverse reads, respectively.

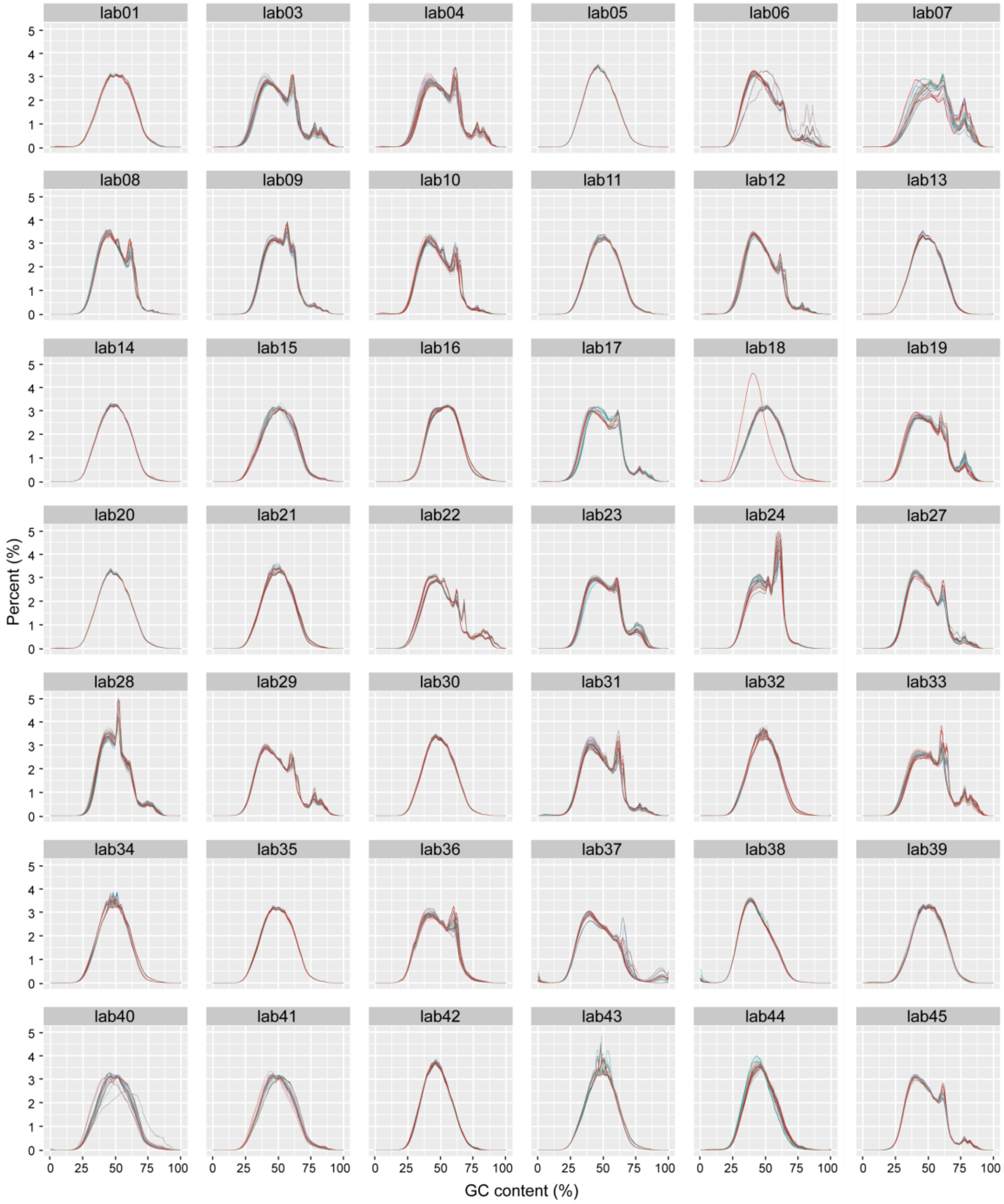

**Supplementary Figure 6. GC distribution of RNA-seq data for Quartet samples.** Three replicates of M8, F7, D5, D6, T1, and T2 were included.

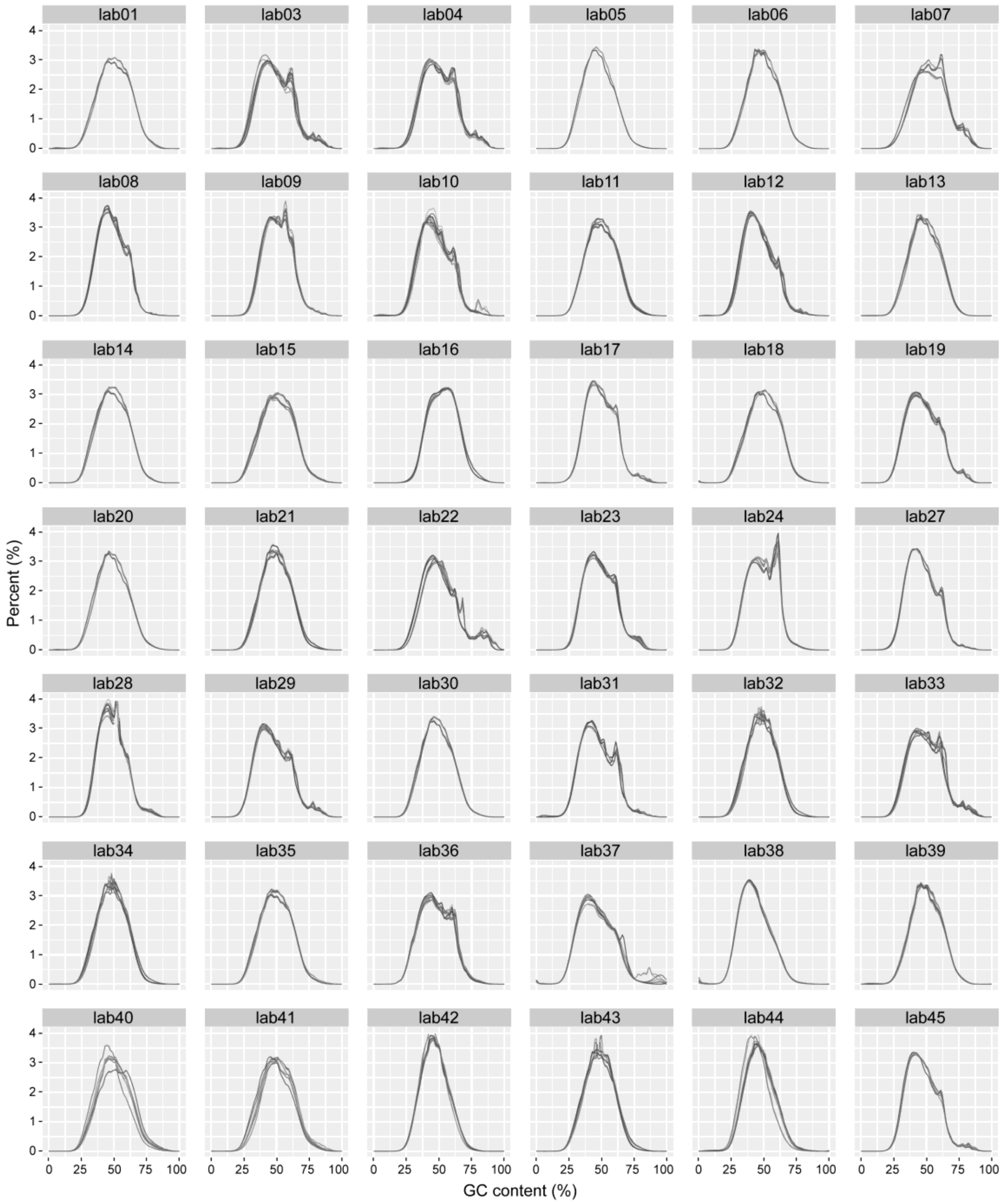

**Supplementary Figure 7. GC distribution of RNA-seq data for MAQC samples.** Three replicates of MAQC A and B were included.

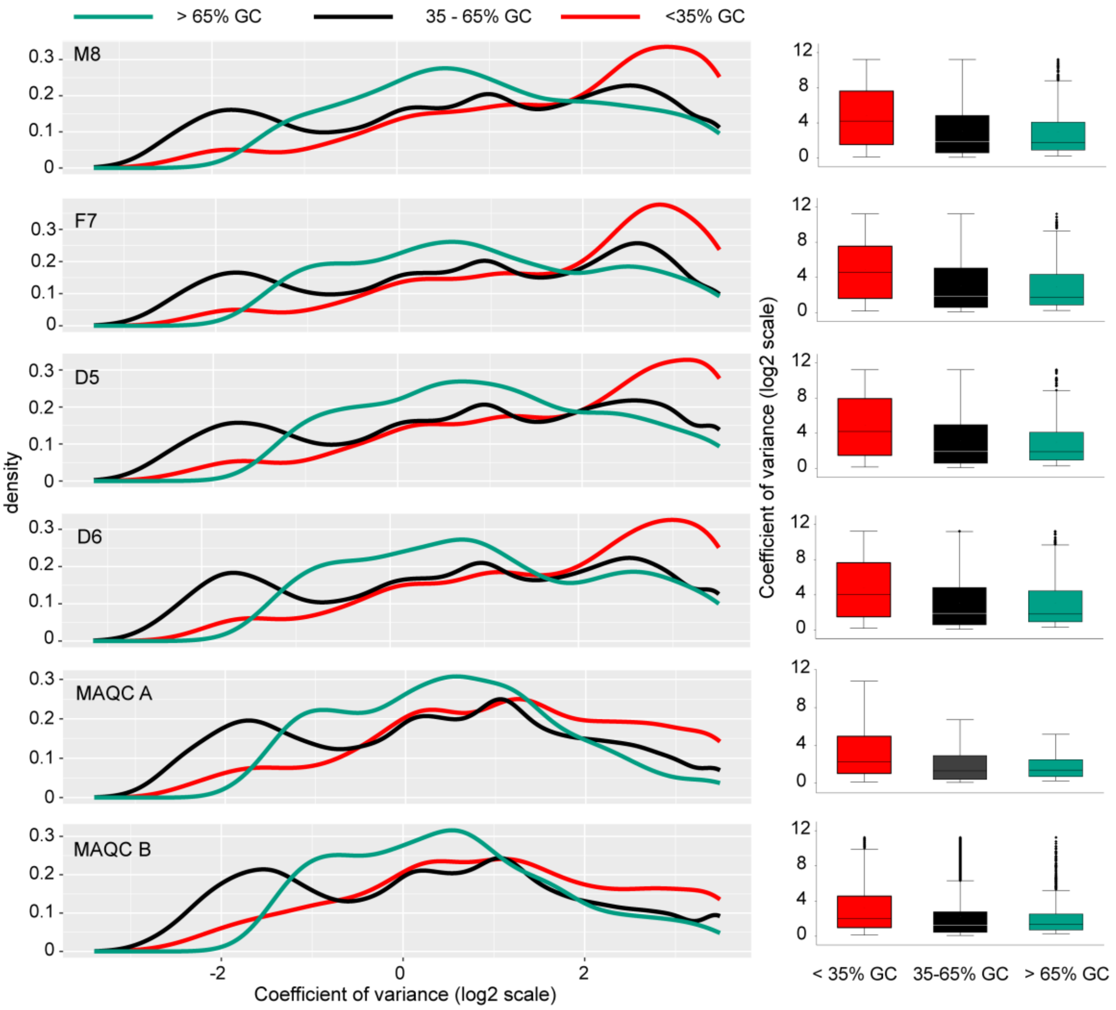

**Supplementary Figure 8. Impact of GC content on inter-laboratory agreement of gene expression.** Low GC content is associated with high inter-laboratory variations of gene expression measurement.

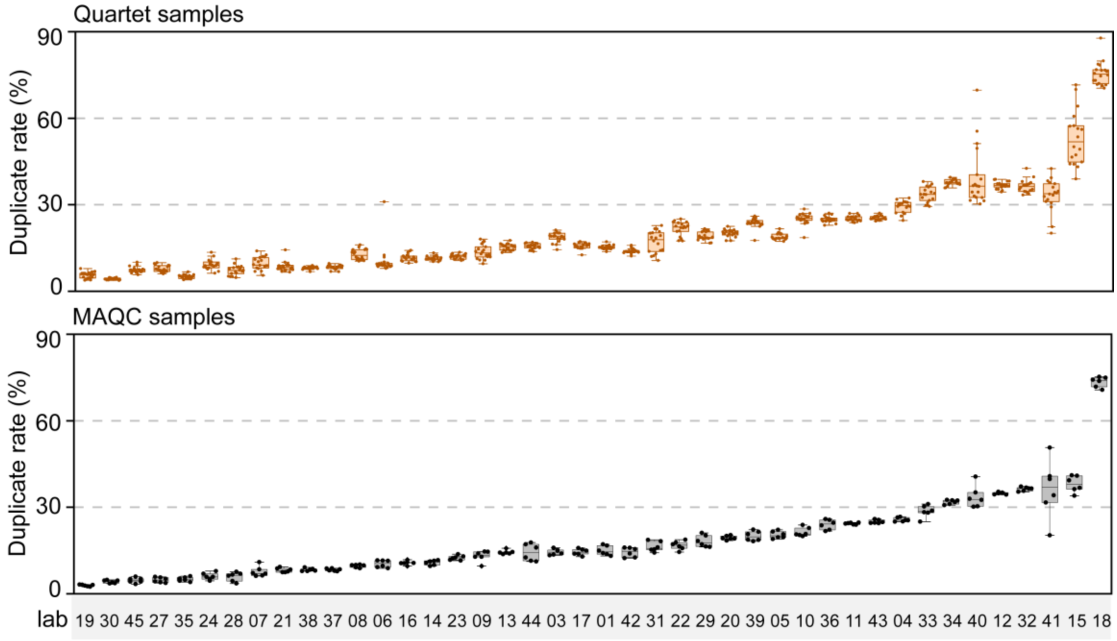

**Supplementary Figure 9. The duplicate rate of RNA-seq data.** The duplicate rate of sequencing reads in Quartet (up) and MAQC (down) samples was calculated using fastp[1]. Previous studies have suggested that the duplicate rate should be below 30% [2-4].

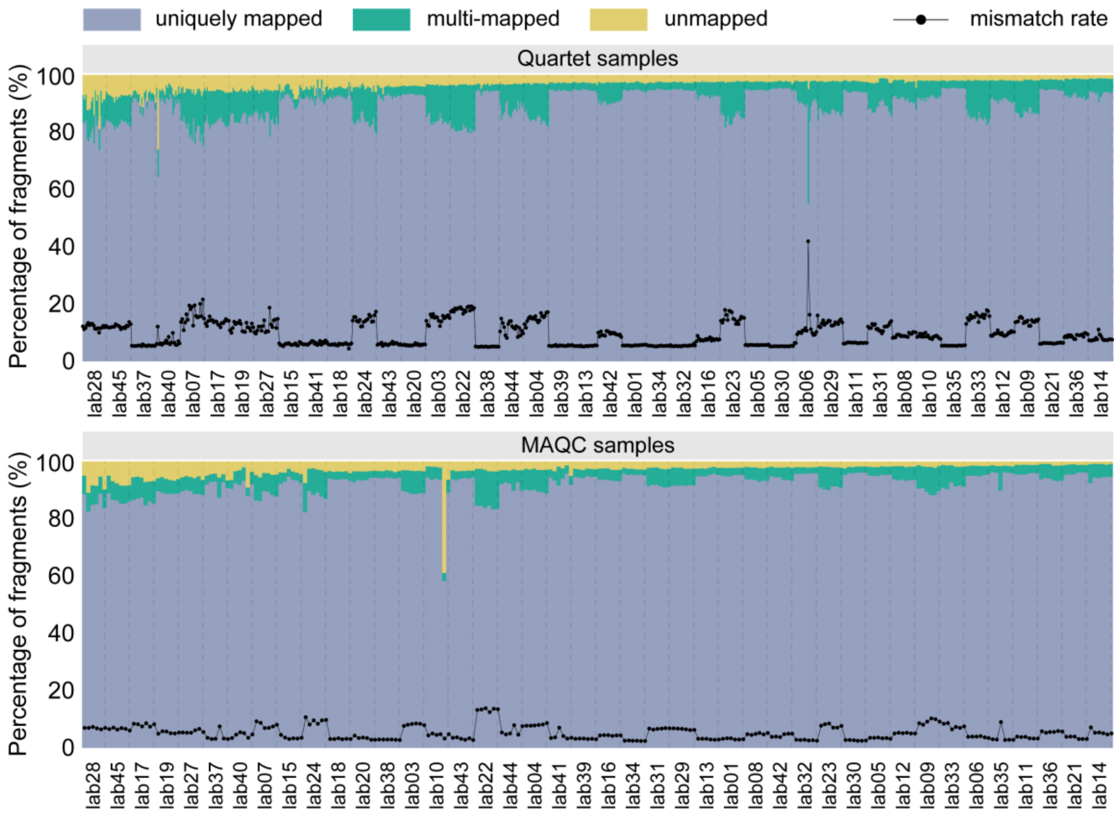

**Supplementary Figure 10. The mapping statistic for all RNA-seq data.** Raw data of Quartet (up) and MAQC (down) samples was mapped to reference genome using STAR [5]. The black dots indicate the mismatch rate.

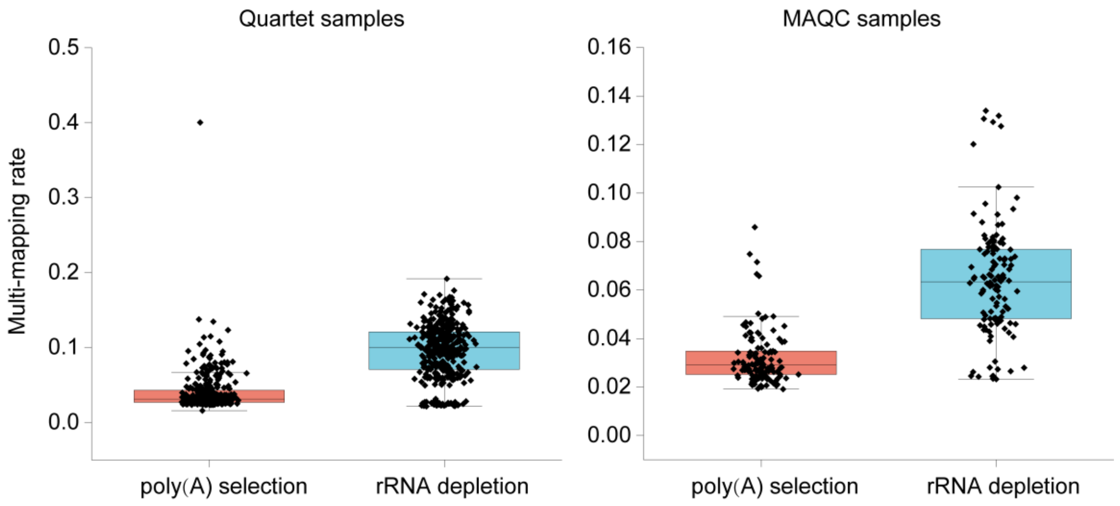

**Supplementary Figure 11. The influence of mRNA enrichment methods on multi-mapping rate.** Raw data of Quartet (left) and MAQC (right) samples was mapped to reference genome using STAR [5]. For both Quartet and MAQC samples, rRNA depletion methods corresponding to a high multi-mapping rate.

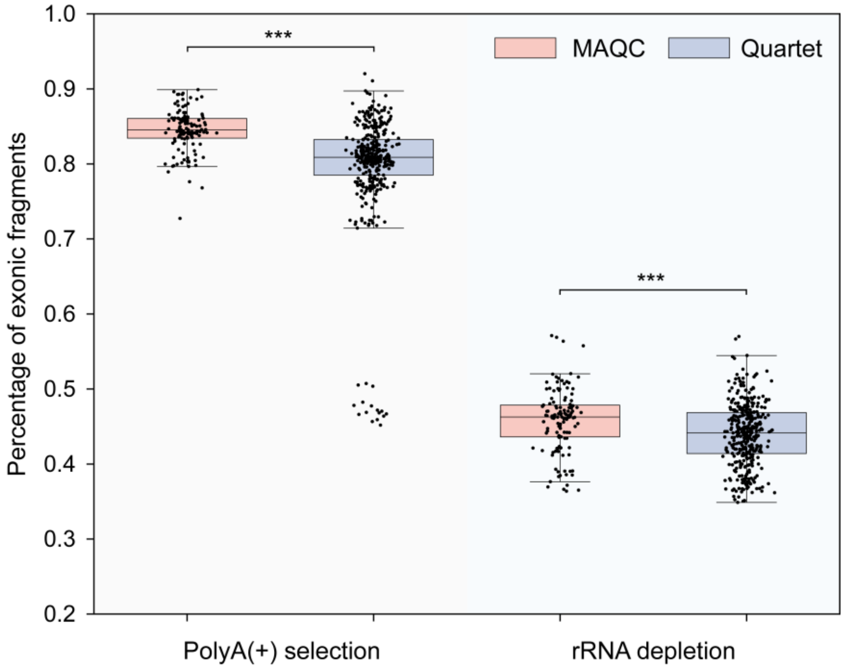

**Supplementary Figure 12. Percentage of exonic reads.** A significant higher exonic reads in MAQC samples than Quartet samples for both poly(A) selection method and rRNA depletion method. This may be attributed to more highly expressed genes in MAQC samples. *** indicates a p-value < 0.001. The significance was tested using unpaired t-test.

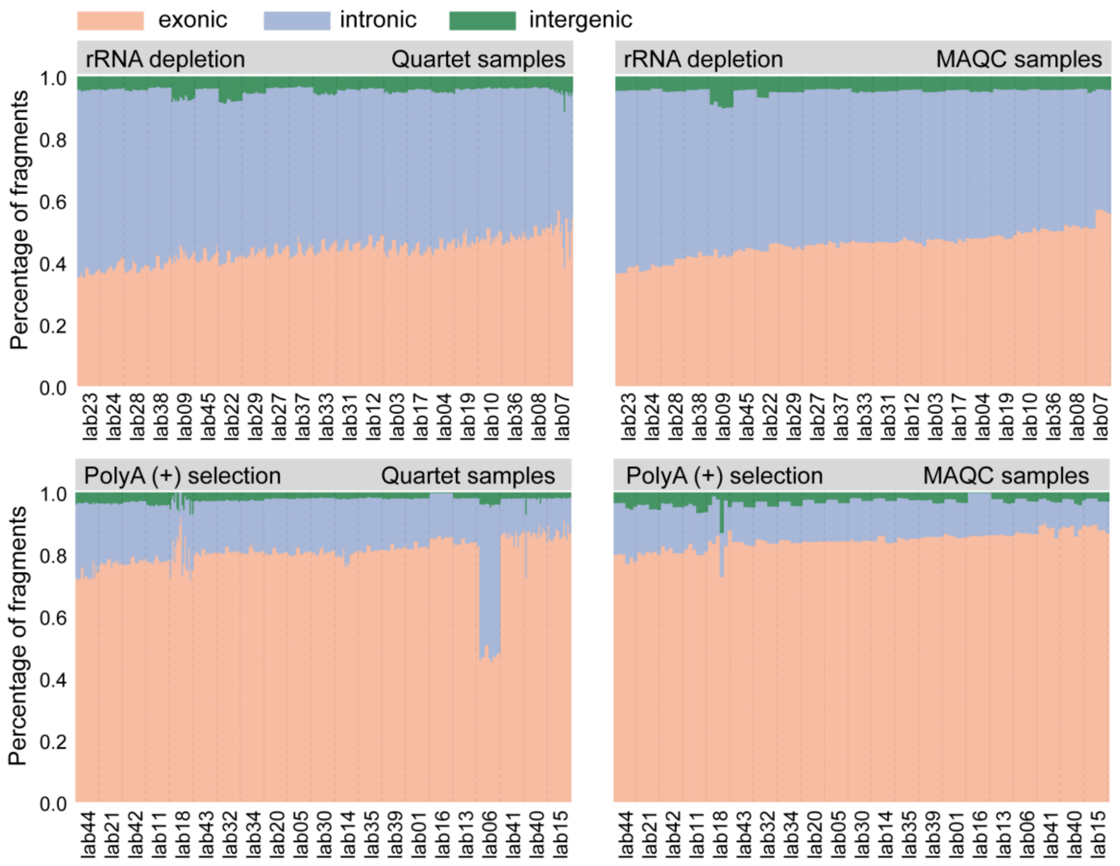

**Supplementary Figure 13. Percentage of reads mapped to exonic, intronic, and intergenic region.** The poly(A) selection protocol consistently associated with higher percentage of exonic reads. For lab06 using the poly(A) selection protocol, quartet samples showed a low percentage of exonic reads, despite the overall high mapping rate, suggesting the presence of potential DNA contamination.

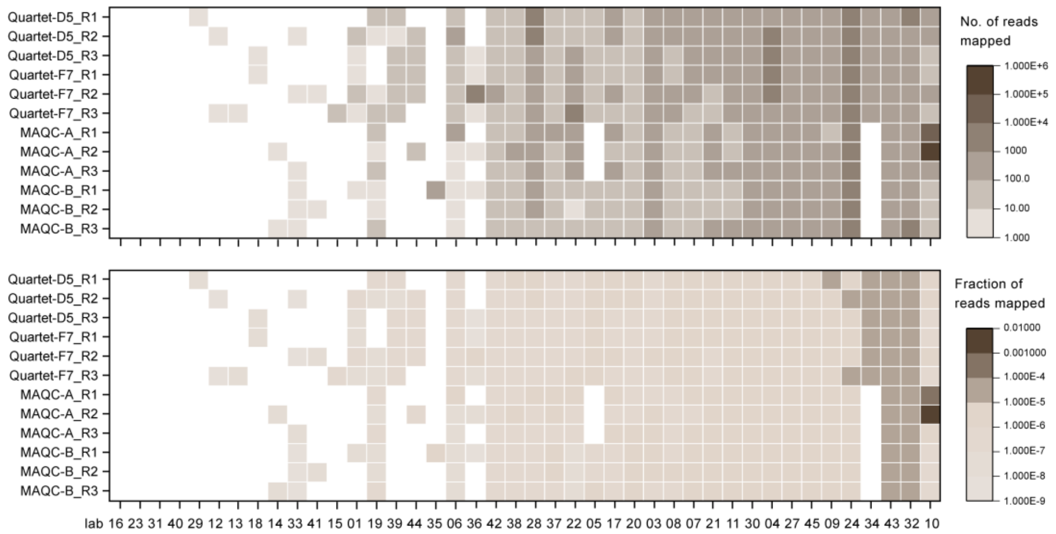

**Supplementary Figure 14. Cross-contamination assessment based on external ERCC controls.** In replicates of MAQC A, B, D5, and F7 samples, reads aligned to the ERCC genes should not be present, offering a valuable opportunity to assess potential cross-contamination.

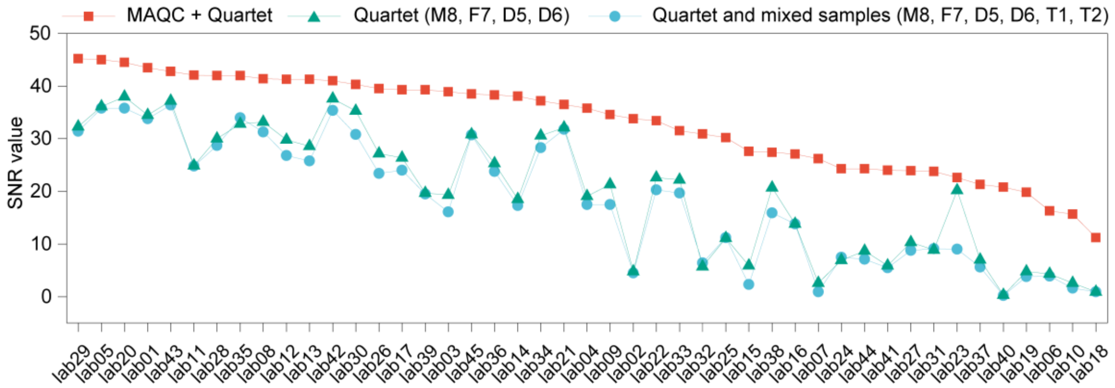

**Supplementary Figure 15. Comparison of SNR values for different combinations of reference material.** The red squares indicate the SNR values from all 24 MAQC and Quartet samples. The cyan triangles indicate the SNR values from M8, F7, D5, D6, T1, and T2, while blue circles indicate the SNR values from M8, F7, D5, and D6. SNR, signal-to-noise ratio.

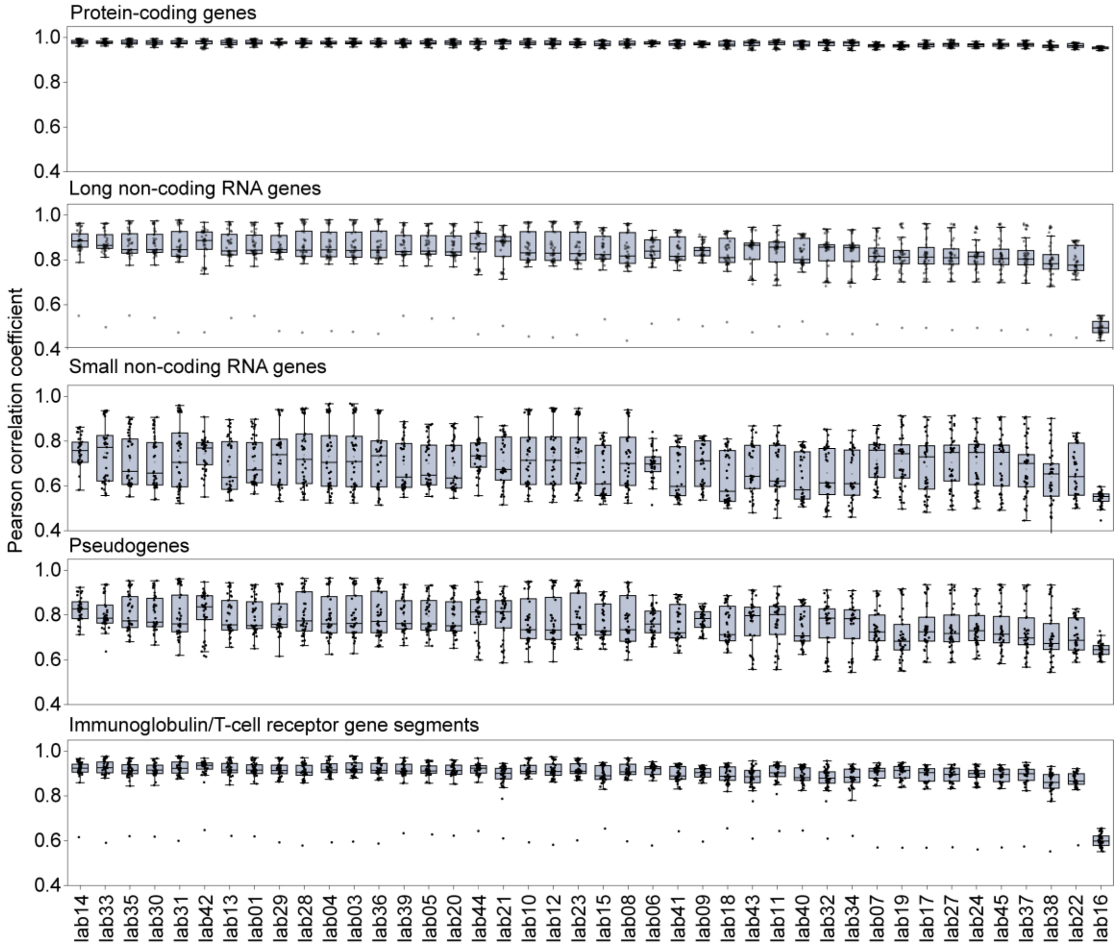

**Supplementary Figure 16. The agreement of gene expression among laboratories for different gene types.** After applying fixed analysis pipeline, the agreement among laboratories was notably high for protein-coding genes but comparatively lower for small non-coding genes and pseudogenes.

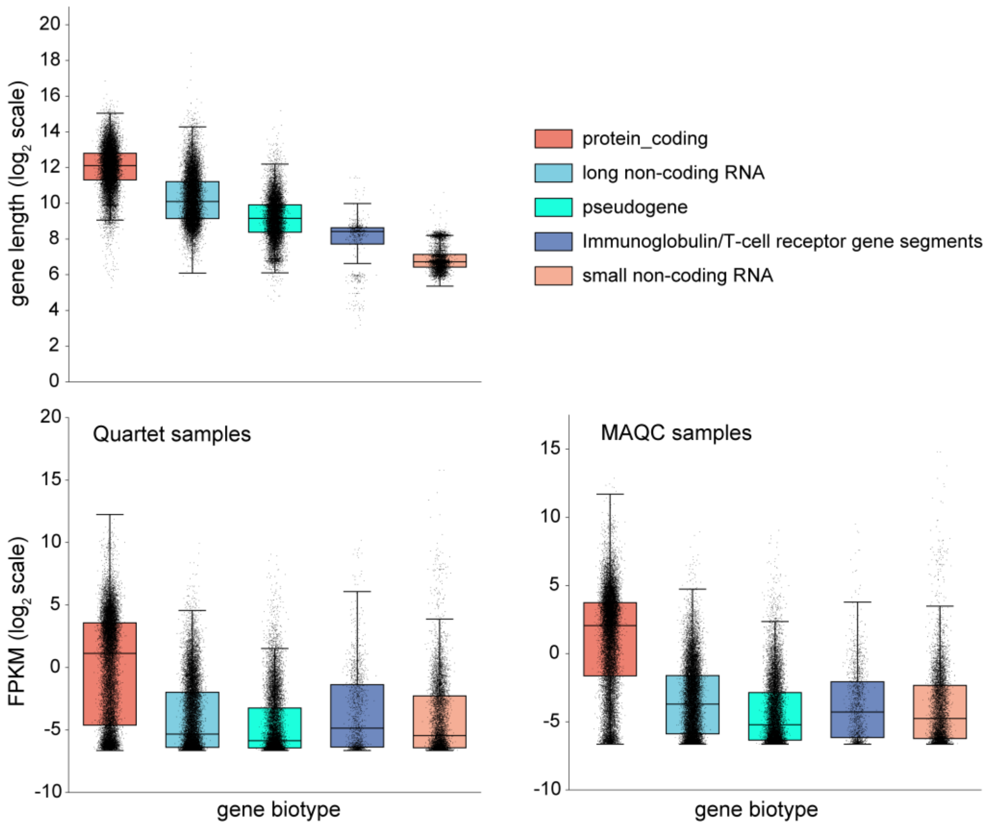

**Supplementary Figure 17. The gene lengths and expression levels for five gene types.** Small non-coding RNA, immunoglobulin/T-cell receptor gene segments, and pseudogene are generally shorter in length, corresponding to a lower inter-laboratory reproducibility. Additionally, in both Quartet and MAQC samples, long non-coding RNA, small non-coding RNA, immunoglobulin/T-cell receptor gene segments, and pseudogene exhibit low expression levels, which also could explain the lower inter-laboratory reproducibility for these gene types.

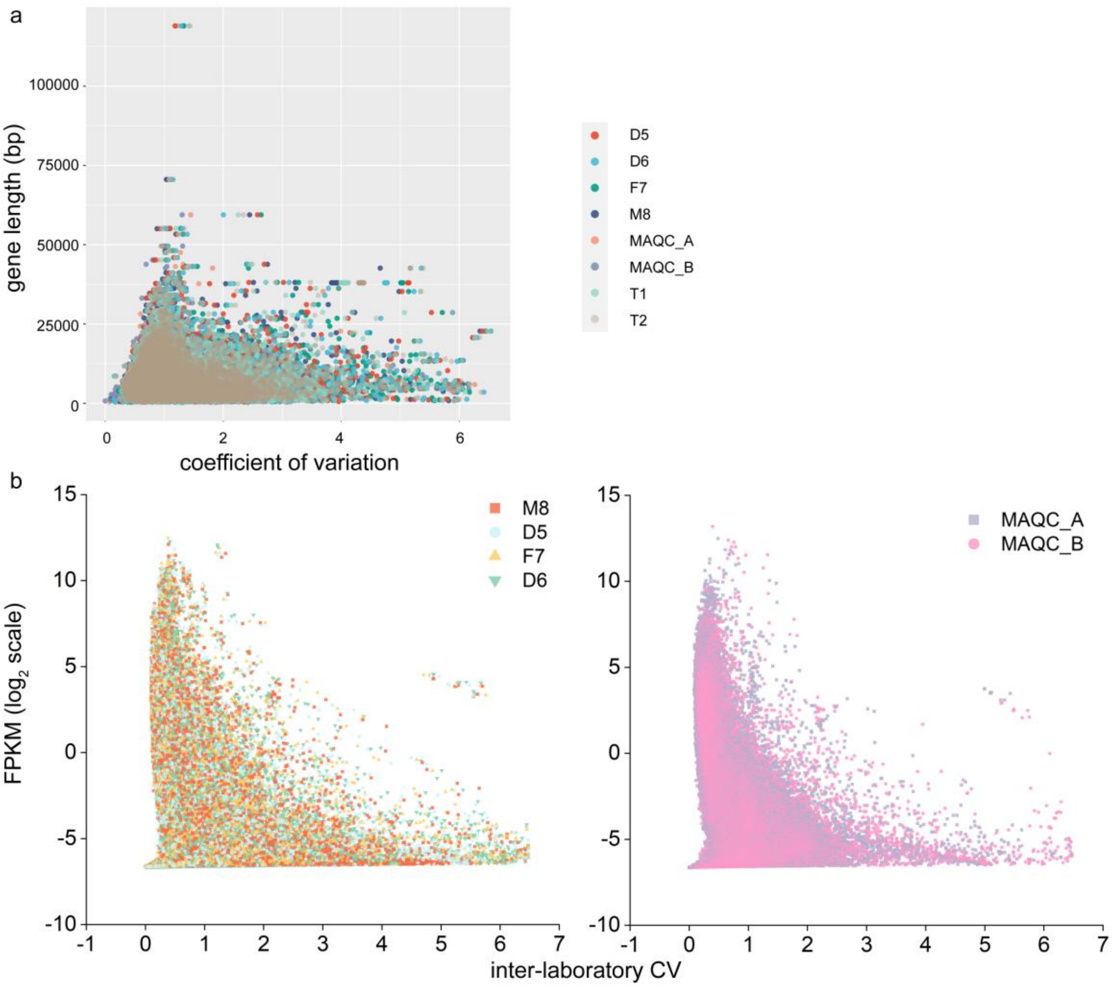

**Supplementary Figure 18. The influence of gene lengths and gene expression on inter-laboratory reproducibility.** (a) Genes with low gene length tended to have a high coefficient of variances (CV) across laboratories. (b) Low-expression genes showed a lower coefficient of variances (CV) across laboratories.

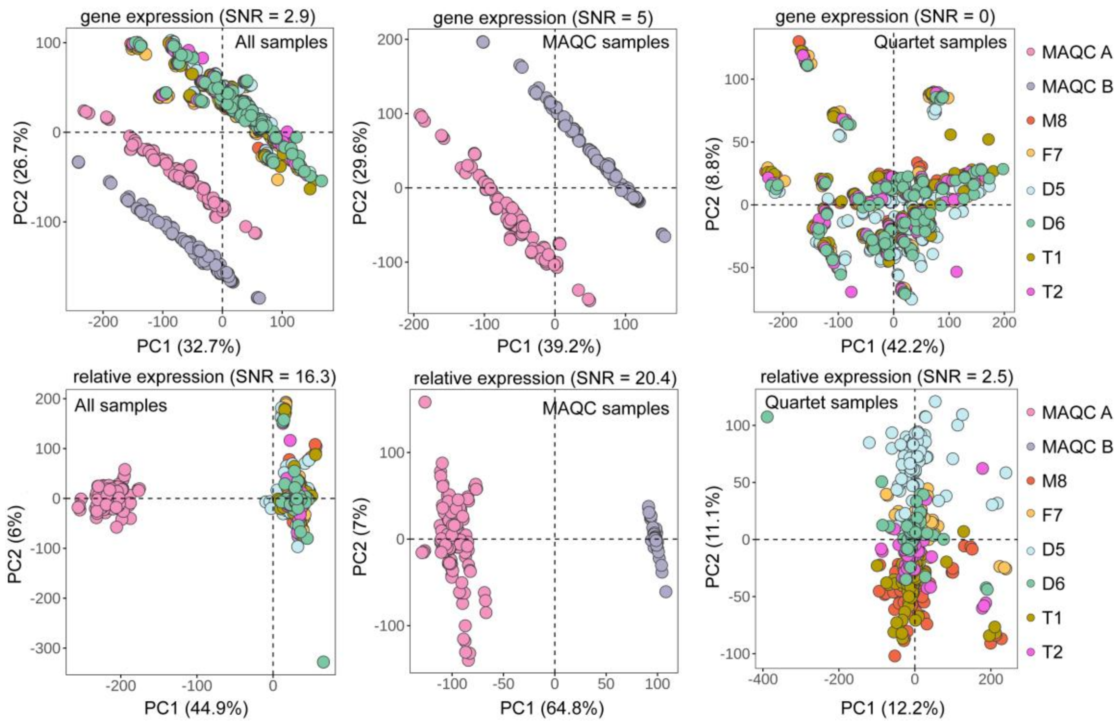

**Supplementary Figure 19. Principal component analysis (PCA) by all pooling RNA-seq data for different sample combinations.** The MAQC samples exhibit huge biological differences, and performing PCA with the inclusion of MAQC samples can lead to an overestimation of the data quality. Relative expression, as opposed to absolute expression, help distinguish samples from inter-laboratory variations. SNR, signal-to-noise ratio.

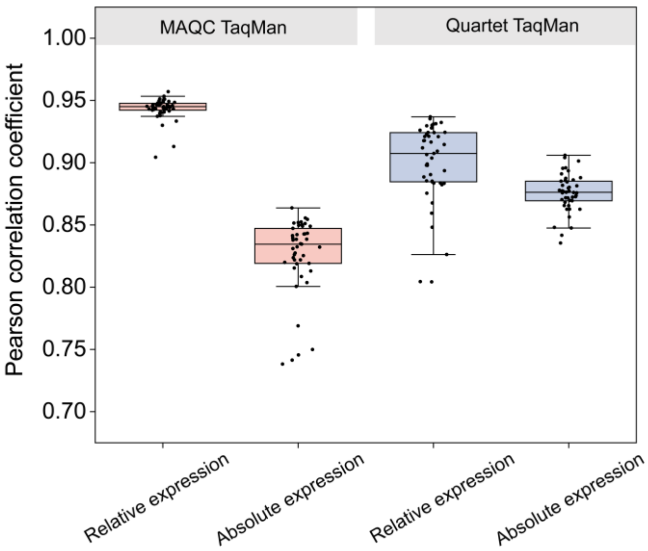

**Supplementary Figure 20. Relative expression measurements were more accuracy than absolute expression measurements.** Based on TaqMan datasets for both Quartet and MAQC samples, relative expression consistently exhibited Pearson higher correlation coefficients when compared to absolute expression.

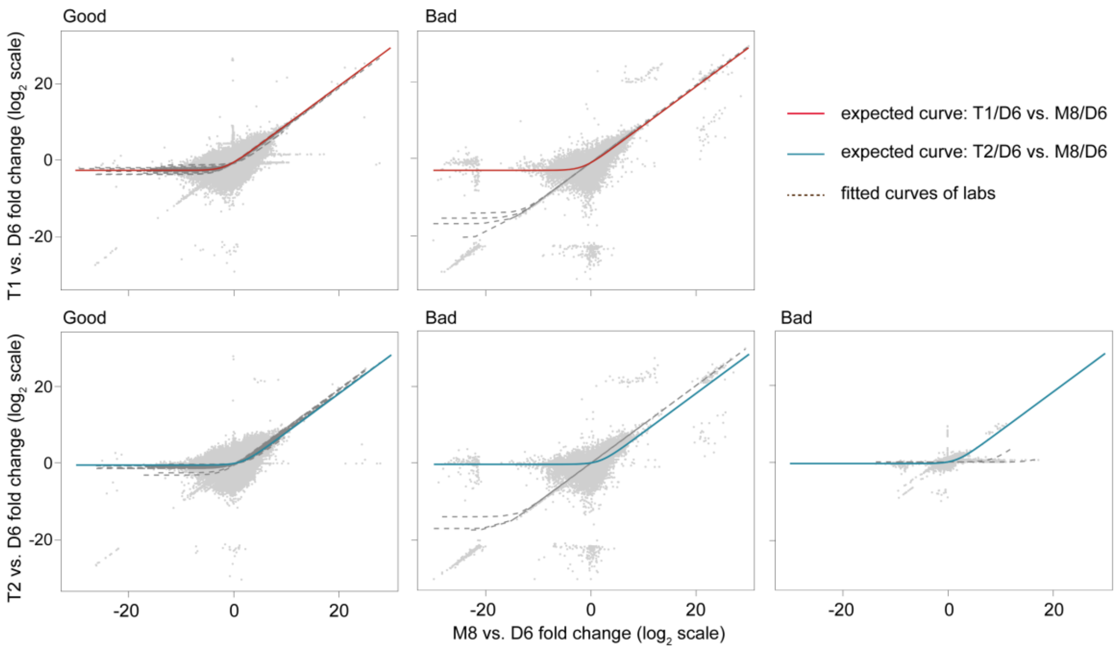

**Supplementary Figure 21. Comparison between laboratories with good and bad recovery of mixture ratios of sample T1 and T2.** The red and cyan solid line traces the expected curve after mRNA/total-RNA shift correction. The grey dashed lines indicate the fitted curves from data of laboratories. The genes reported by all laboratories are shown in grey. Primarily, the presence of genes with extremely large fold differences resulted in the deviation of the fitting curve. These records were attributed to erroneous calculation of low-expressed genes in individual samples.

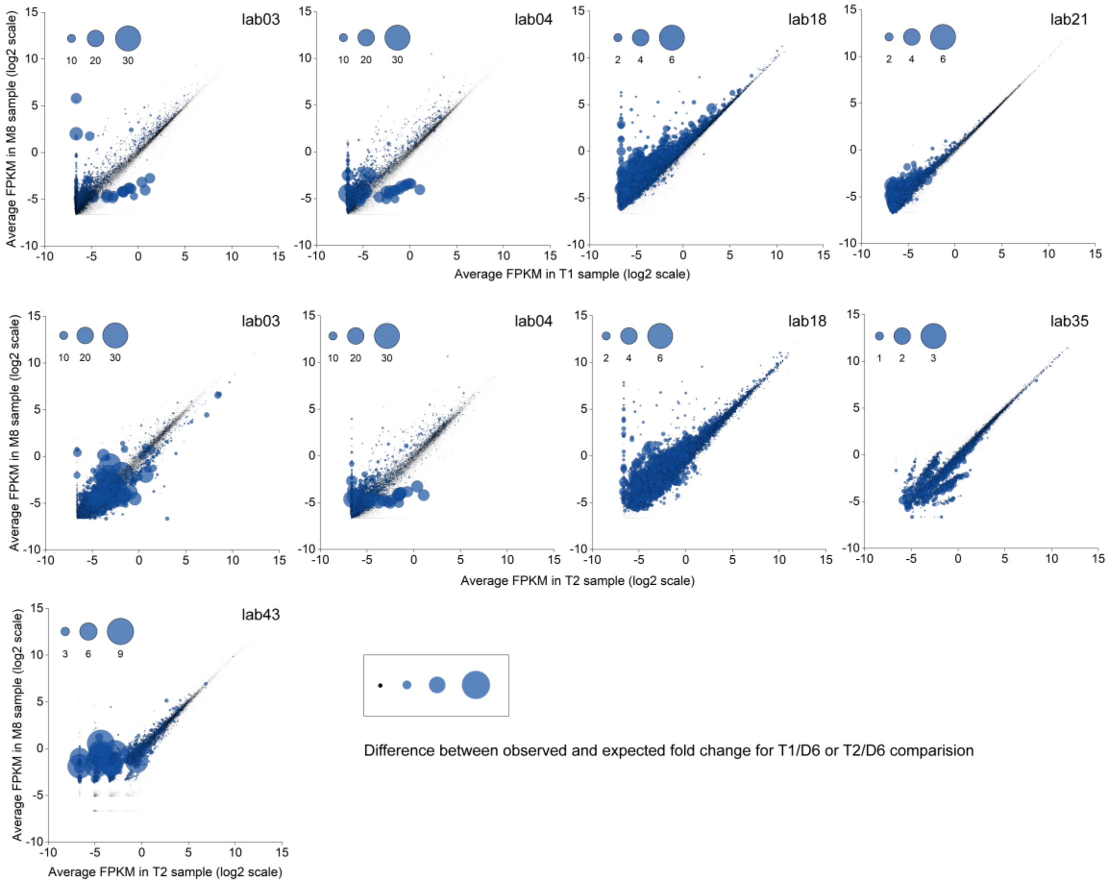

**Supplementary Figure 22. Presentation of genes in RNA-seq data with poor recovery of mixed ratios.** We focused on four laboratories exhibiting poor recovery of mixing proportions for T1 samples and five laboratories with poor recovery for T2 samples (Supplementary Figure 21). The absolute differences between the expected and observed fold changes for T1/D6 or T2/D6 comparisons were calculated for all genes, and the impact of gene expression levels on such absolute differences was analyzed. In the plot, the absolute differences are represented by the size of circles, where larger circles indicate greater differences between the expected and observed fold change for the respective genes. These results indicate that genes with poor recovery of expected mixing ratios (larger circles) tend to exhibit lower expression levels.

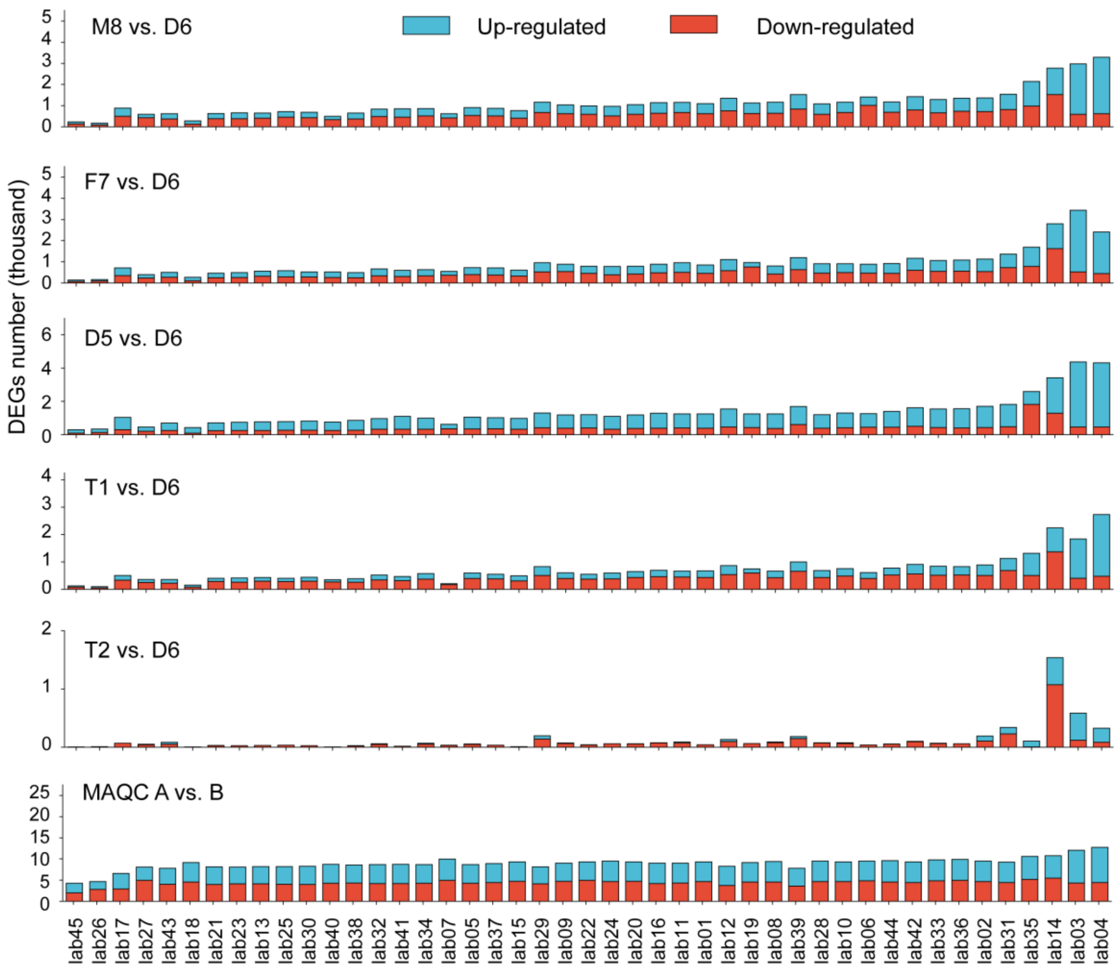

**Supplementary Figure 23. Comparison of the number of DEGs among laboratories.** For the purpose of facilitating inter-laboratory comparisons, only the number of differentially expressed genes (DEGs) among protein-coding genes were compared. The laboratories demonstrated significant variations in the number of DEGs.

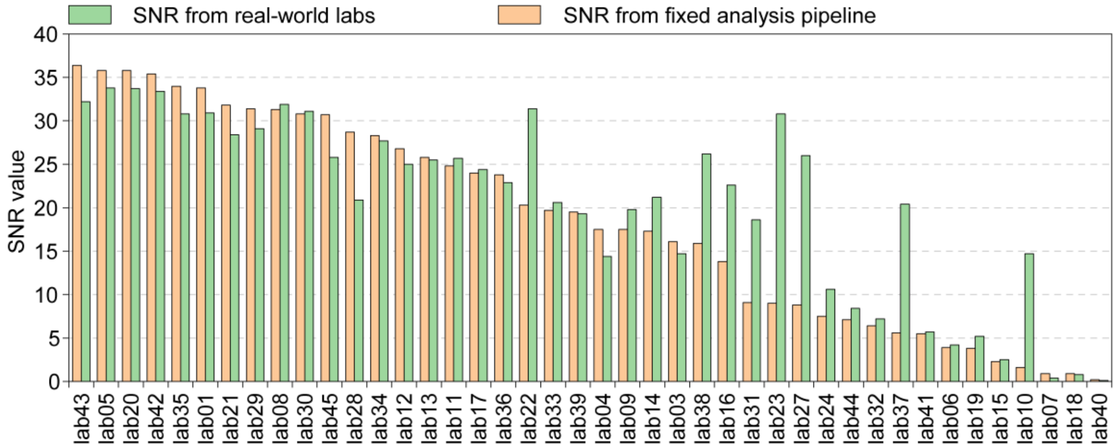

**Supplementary Figure 24. Data quality (SNR values) after applying fixed analysis pipeline.** More than half of the laboratories demonstrated an increased SNR after applying the uniformed analysis pipeline, especially for those with initially lower SNR values, indicating the failure of bioinformatics pipelines in these laboratories. However, some laboratories experienced a decrease in SNR values after applying the uniformed analysis pipeline. This observation may be related to the different number of genes considered, as the uniformed analysis pipeline encompassed all genes with non-zero read counts, whereas real laboratories included a smaller gene set due to their gene type preferences and the selections of gene annotation and analysis tools. SNR, signal-to-noise ratio.

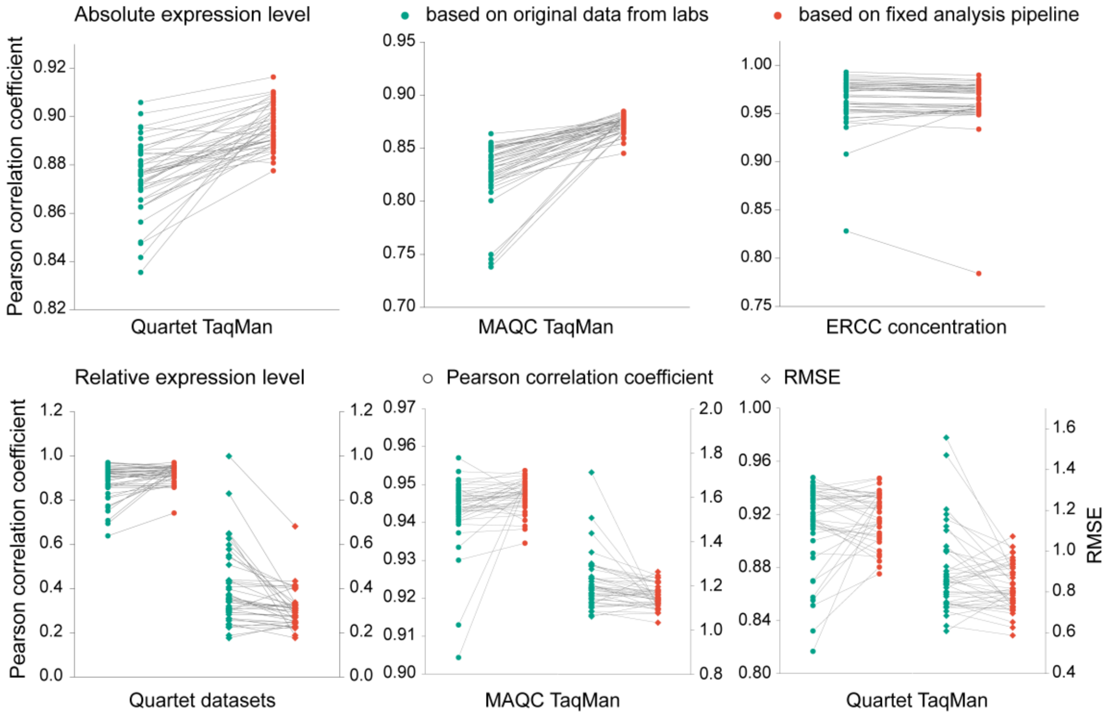

**Supplementary Figure 25. The accuracy of absolute and relative expression after applying a fixed analysis pipeline.** After applying the analysis pipeline, Ensembl-STAR-StringTie, Pearson correlation coefficients between relative expression and the reference datasets increased. The red dots represent the metric calculated from the data submitted by laboratories, whereas the cyan dots indicate the metric after applying the fixed analysis pipeline. The circles indicate Pearson correlation coefficient between laboratories and reference datasets, and the diamonds indicate the Root Mean Square Error (RMSE).

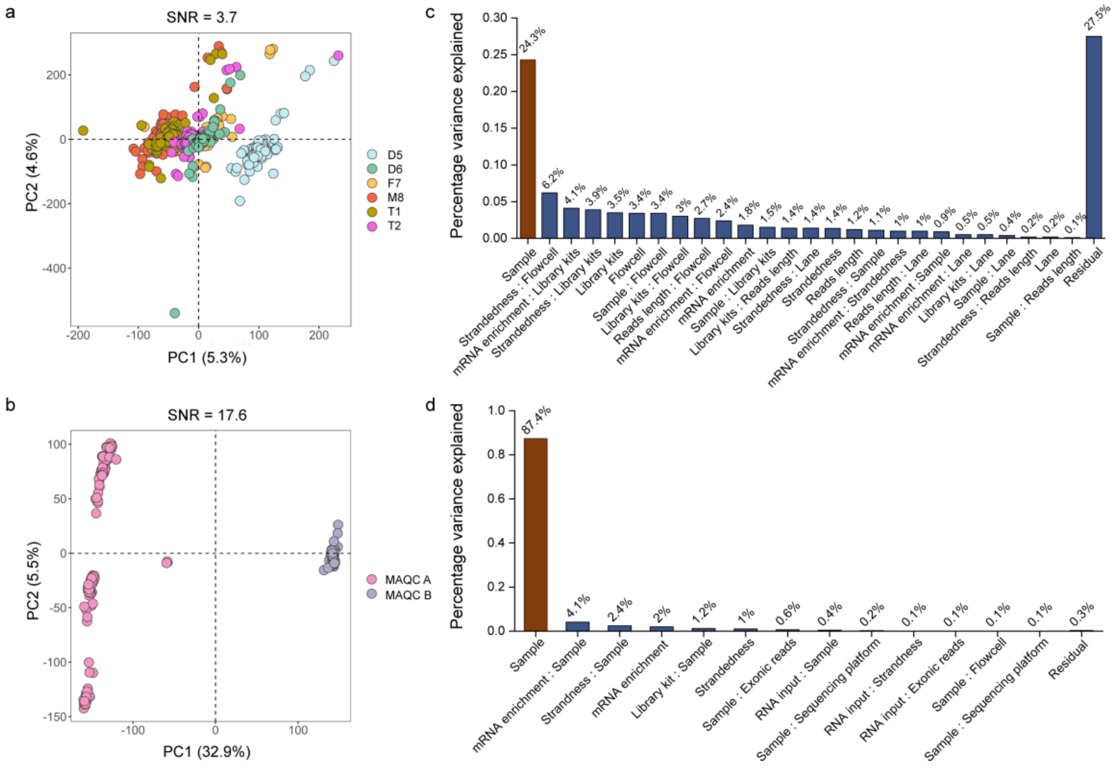

**Supplementary Figure 26. Calculation of relative expression could eliminate the variations from experimental process.** (a) Scatterplots of PCA on RNA-seq data of all laboratories for Quartet samples and (b) MAQC samples at relative expression levels. (c) Principal variance component analysis quantifies the proportion of variance explained by each experimental factor for Quartet samples, (d) and MAQC samples. The red bars represent biological differences between samples. When calculating relative expression, the relative contribution of each experimental factor to total variations decreased compared to the biological difference across Quartet (up) or MAQC (down) samples. For Quartet samples, there still 27.5% of variations from experimental process that cannot be eliminated, implying the impact of unknown confounding factors. SNR, signal-to-noise ratio.

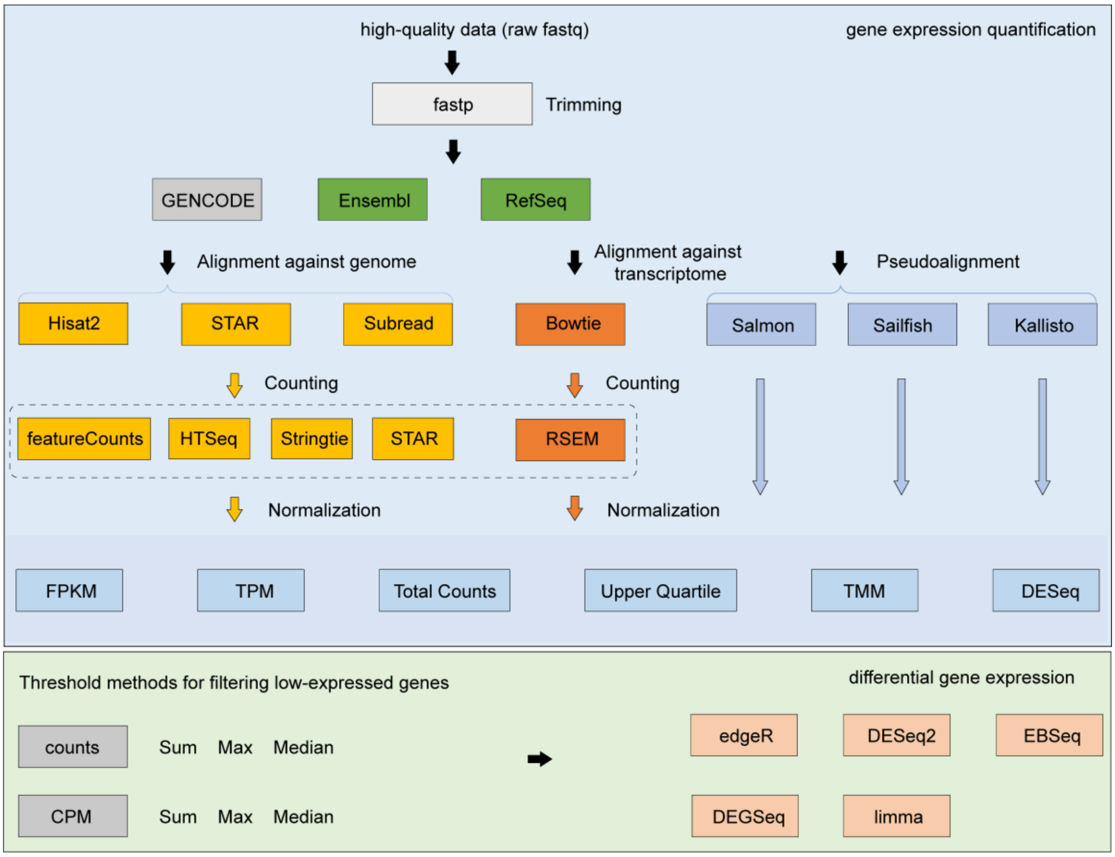

**Supplementary Figure 27. The benchmark analysis workflow.** Upper panel represents the raw gene expression quantification workflow. Every box contains the algorithms or methods used for the RNA-seq analysis at trimming, alignment, counting, and normalization. The lower panel represents the threshold methods evaluated for filtering low-expressed genes and the algorithms used for the differential gene analysis.

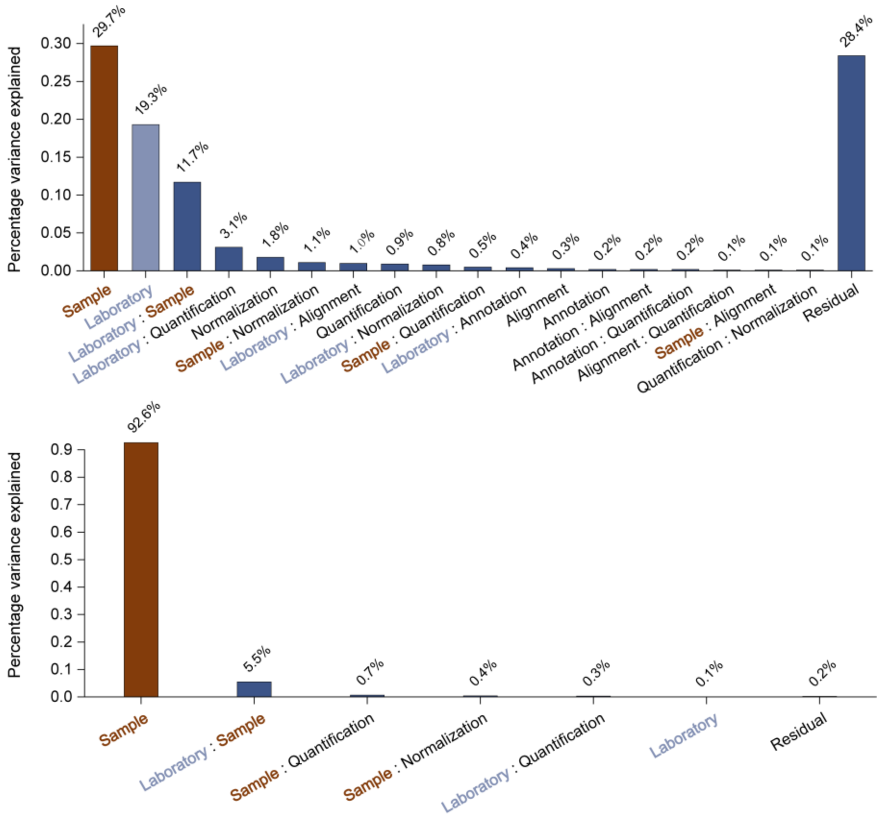

**Supplementary Figure 28. Calculating relative expression could correct the influences of different bioinformatics tools.** When calculating relative expression, the relative contribution of each bioinformatics step to total variations decreased compared to the biological difference across Quartet (up) or MAQC (down) samples. For Quartet samples, there still 28.4% of variations from bioinformatics process that cannot be eliminated, implying the inherent performance difference in various analysis tools. The red bars represent biological differences between samples, while the light blue bars represent differences among benchmark datasets from 13 different laboratories.

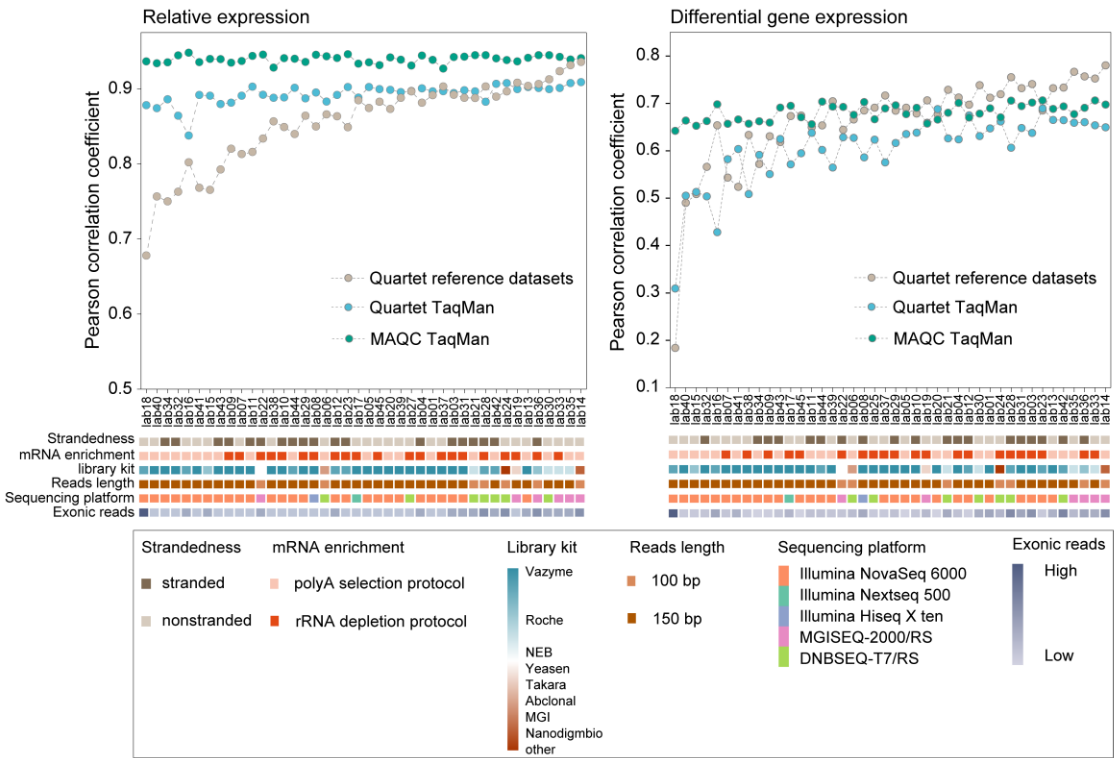

**Supplementary Figure 29. The accuracy of 42 experimental processes after applying the uniformed analysis pipeline.** We examined the laboratories that exhibited higher consistency with the Quartet reference datasets and TaqMan datasets at relative expression levels, and found that they were not exclusively associated with specific wet bench protocols **(Fig. 3f)**. This indicates that while specific factors contribute to inter-laboratory variations, each experimental protocol has the potential to generate accurate expression profiles.

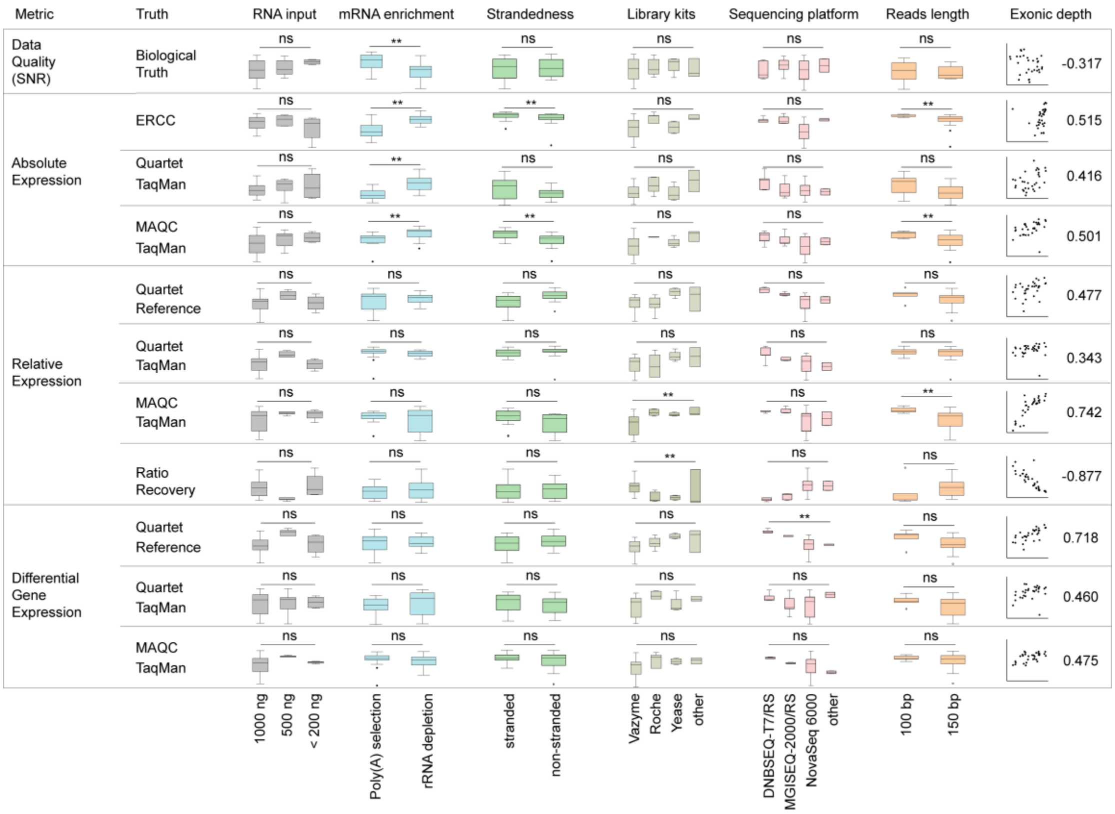

**Supplementary Figure 30. The influence of experimental factors under different performance metrics.** Performance metrics are divided into data quality, accuracy of absolute and relative expression and differential gene expression. The impact of exonic coverage is evaluated by Spearman correlation analyzes. Significance testing was conducted based on normal distribution assumptions using one-way analysis of variance (ANOVA) and paired t-tests, or, in cases where normal distribution was not observed, independent samples were subjected to Kruskal-Wallis test and Mann-Whitney U test. ** indicates a p-value $< 0.05$. ns, not significant.

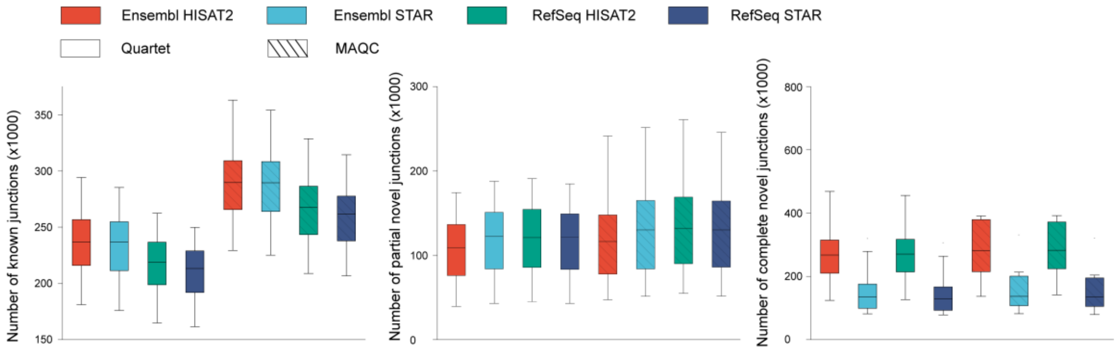

**Supplementary Figure 31. Comparison of junctions detected by four alignment schemes.** All the junctions supported by at least one reads in one of three replicates were included.

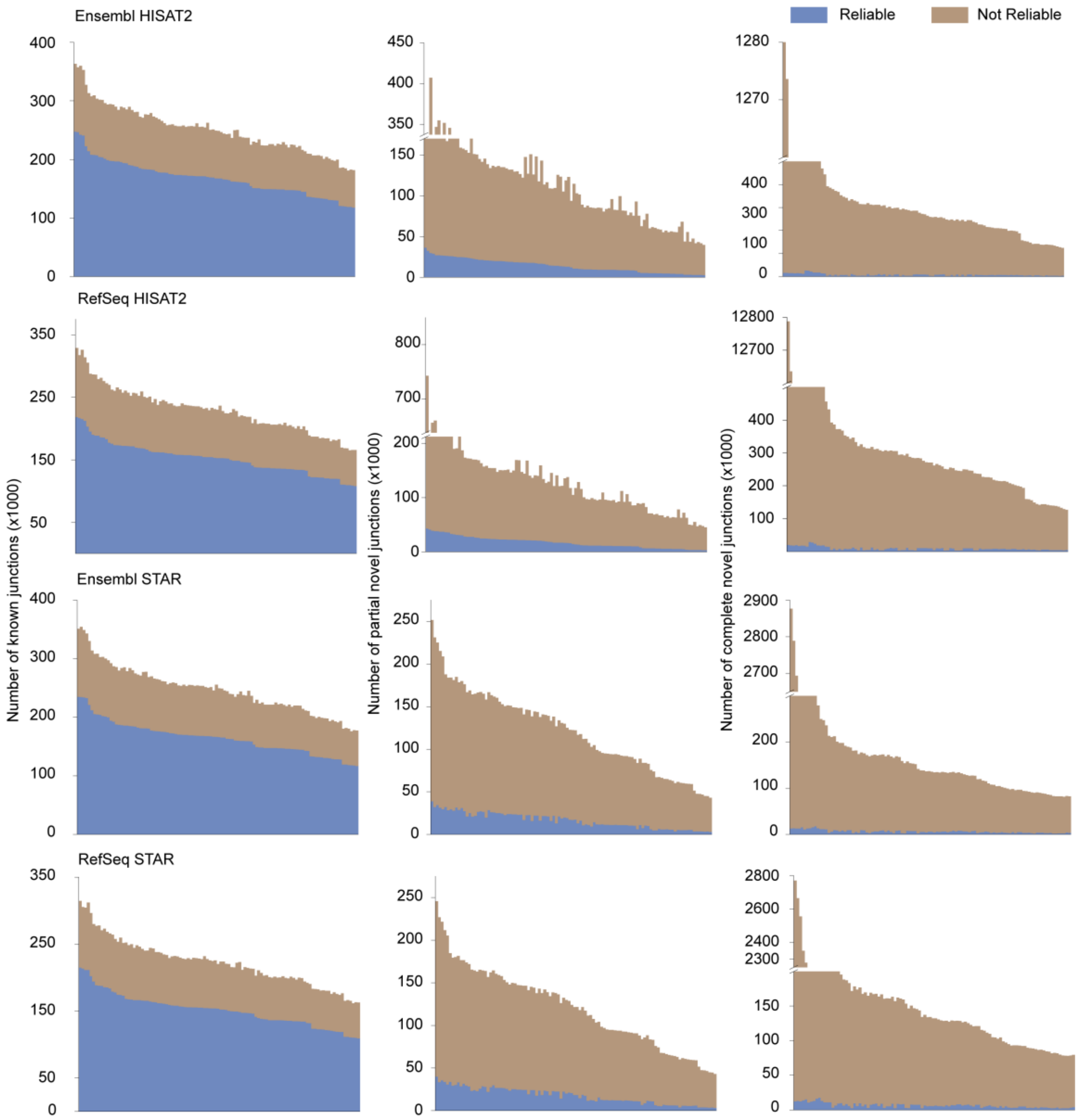

**Supplementary Figure 32. The proportion of reliable and unreliable junctions detected by the four alignment schemes.** Reliable junctions were defined as those supporting by at least one reads for all three replicates, while unreliable junctions were defined as those lacking read support in at least one of the three replicates. Most of the known junctions are reliable, while most of the novel junctions are unreliable.

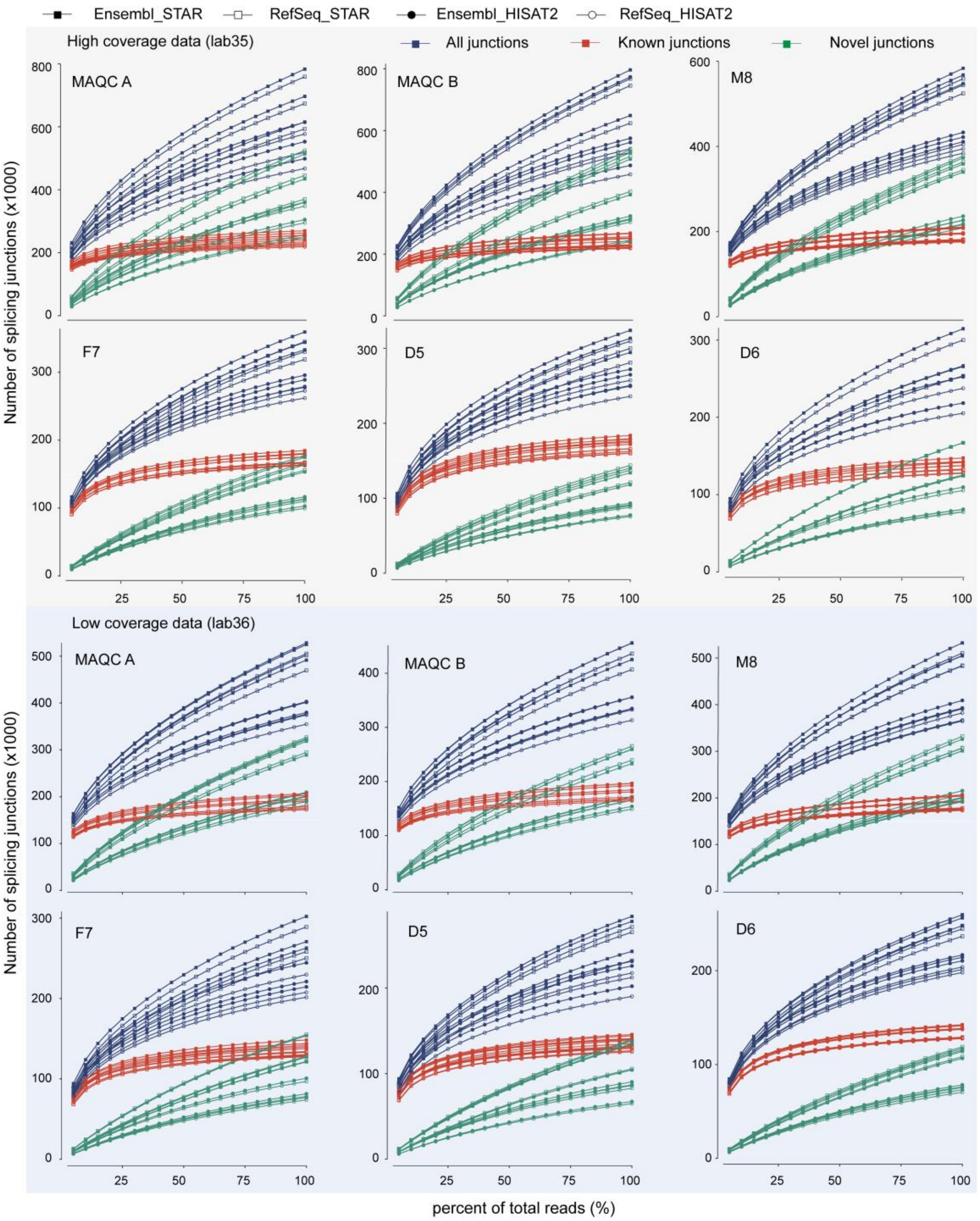

**Supplementary Figure 33. Examining whether the sequencing depth is sufficient for detecting junctions.** Representative data with high and low sequencing depth from two laboratories were resampled, and junctions were detected at each depth levels. Increasing sequencing depth appears to primarily benefit the detection of novel junctions.

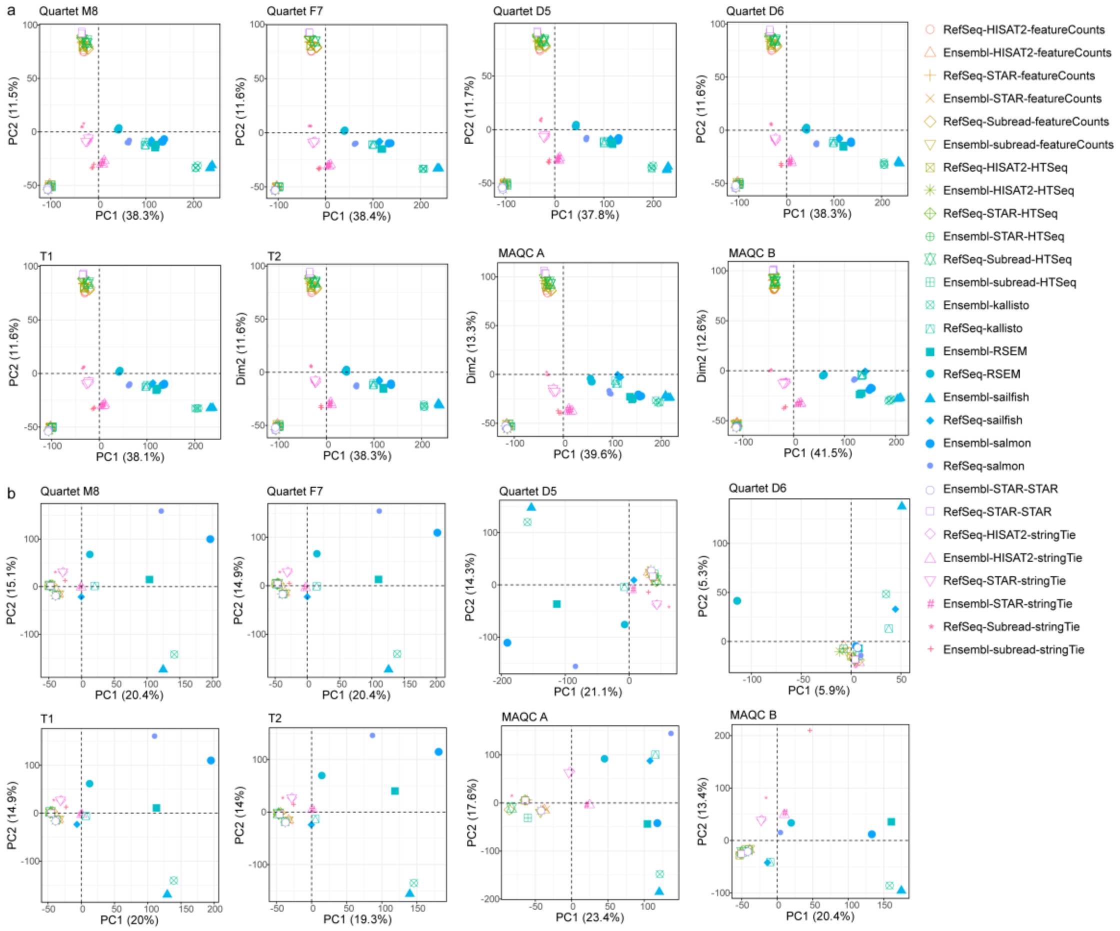

**Supplementary Figure 34. Scatterplots of PCA on RNA-seq data from the 28 quantification pipelines (marked in shapes).** Clustering patterns of all quantification pipelines at absolute (a) and relative expression levels (b). All quantification pipelines are clustered primarily based on exon-level quantification tools such as featureCounts, HTSeq, StringTie, and STAR, or transcript-level quantification tools like RSEM, Salmon, kallisto, and Sailfish. In addition, these pipelines are dispersed by different gene annotations and alignment tools. Especially, gene annotations have a significant impact on the absolute expression measurements of quantitative tools such as featureCounts, HTSeq, and STAR. Different gene annotation exhibited more pronounced impact on the relative expression measurements of transcript-level quantification tools. The RNA-seq data from lab01 were included for PCA analyzes, and identical shapes and colors represent replicates in the PCA plots.

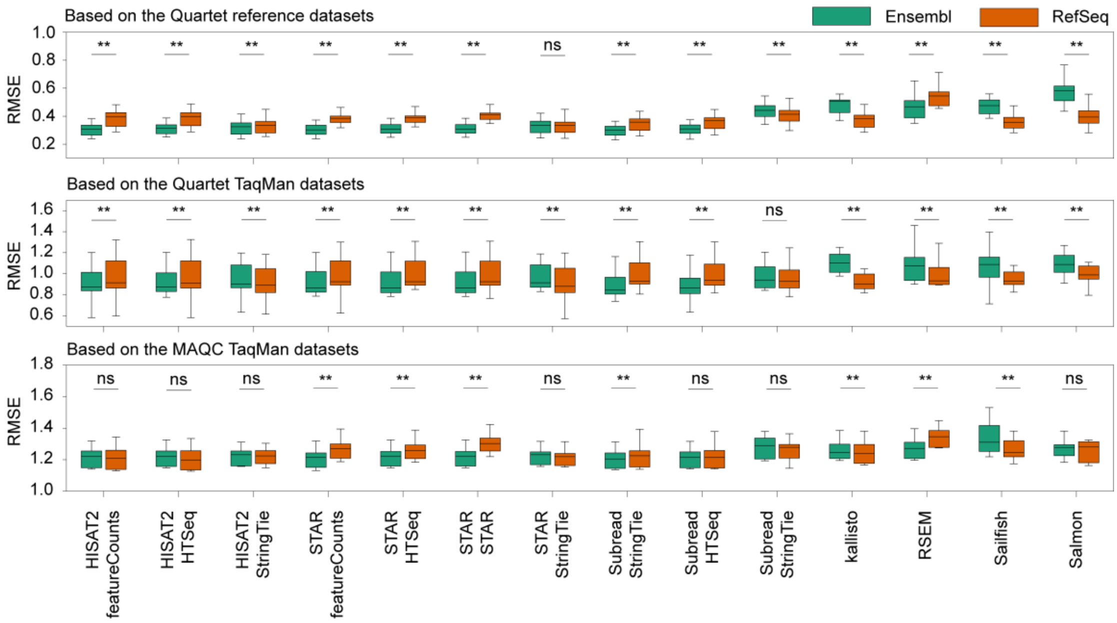

**Supplementary Figure 35. The impact of gene annotation on the accuracy of relative expression.** All 28 quantification pipelines are evaluated based on the Quartet reference datasets, and TaqMan datasets for Quartet and MAQC samples. The impact of gene annotations was analyzed under the same conditions of alignment and quantification tools. Box plots represent 13 benchmarked datasets. Significance testing was conducted using paired t-tests. ** represents p-value < 0.05. ns, not significant.

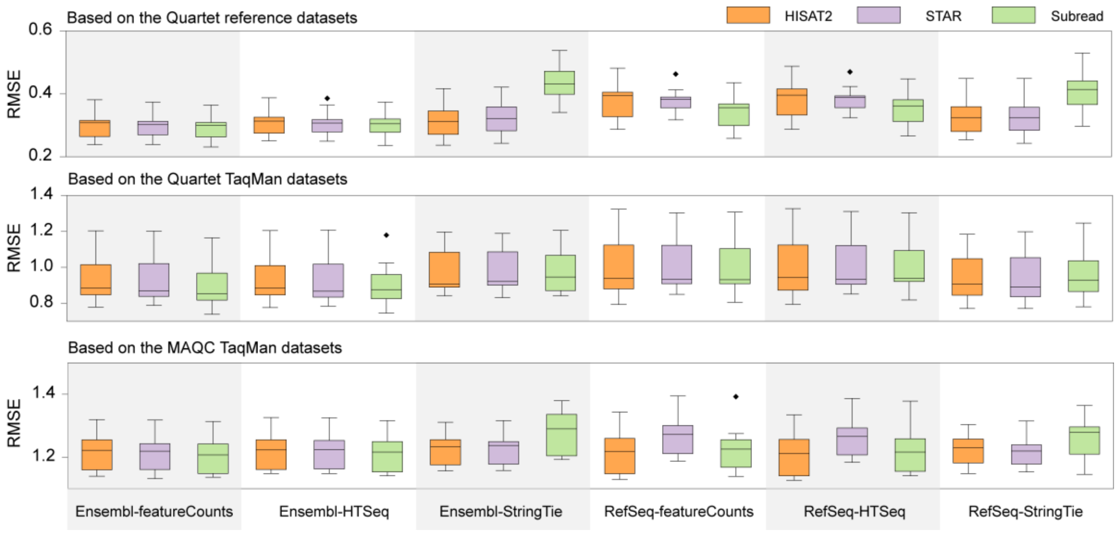

**Supplementary Figure 36. The impact of alignment tools on the accuracy of relative expression.** All 18 quantification pipelines involving sequence alignment are evaluated based on the Quartet reference datasets, and TaqMan datasets for Quartet and MAQC samples. The impact of alignment tools was analyzed under the same conditions of gene annotation and quantification tools. Box plots represent 13 benchmarked datasets.

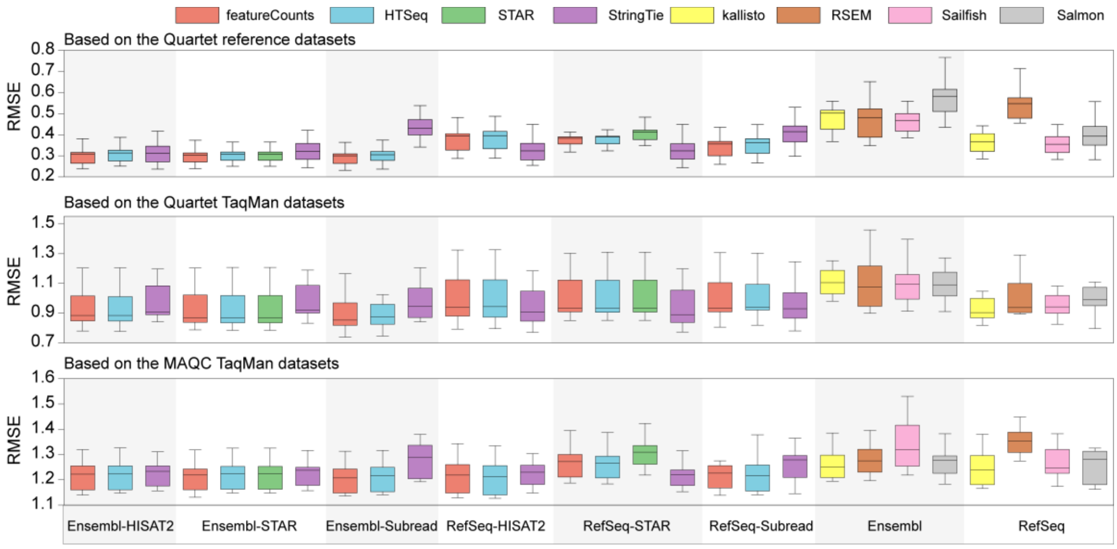

**Supplementary Figure 37. The impact of quantification tools on the accuracy of relative expression.** All 28 quantification pipelines are evaluated based on the Quartet reference datasets, and TaqMan datasets for Quartet and MAQC samples. The impact of quantification tools is analyzed under the same conditions of gene annotation and alignment tools. Box plots represent 13 benchmarked datasets.

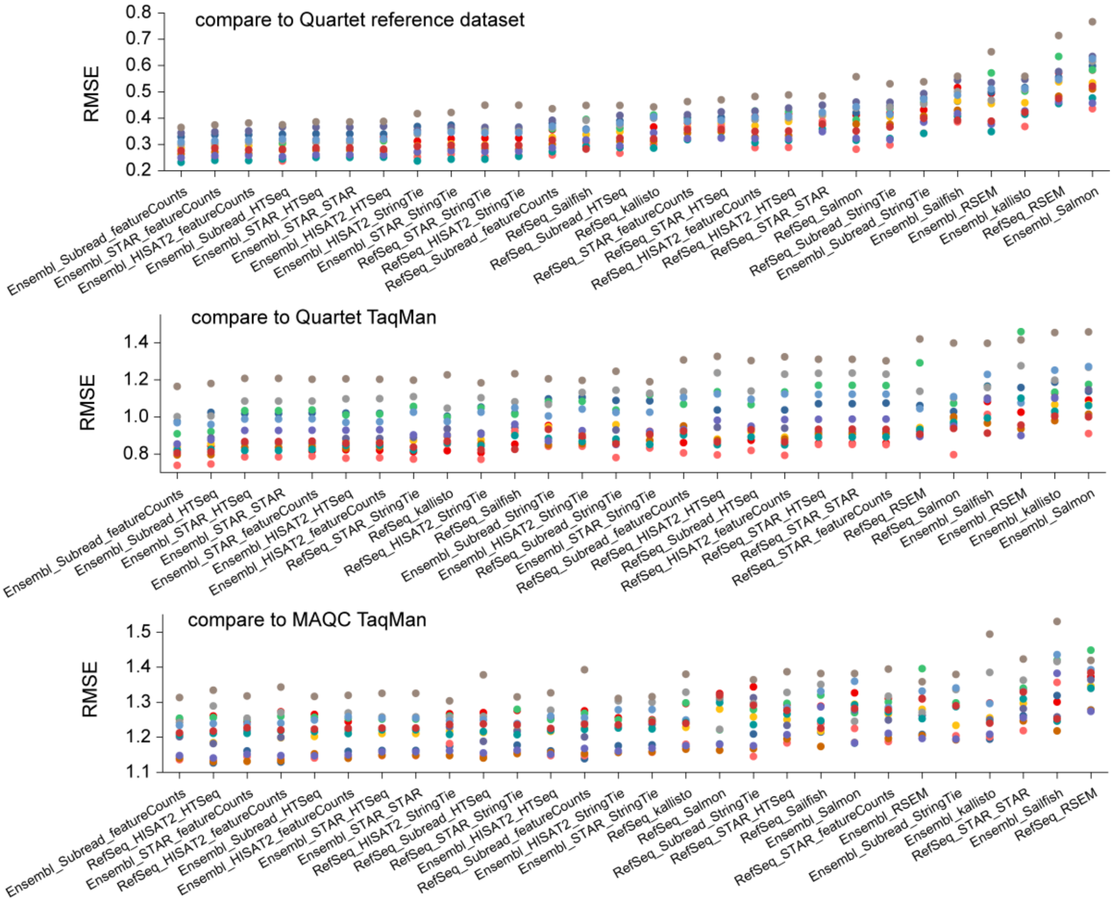

**Supplementary Figure 38. The performance of 28 quantification pipelines at relative expression levels.** The 28 quantification pipelines were ranked based on the median RMSE for 13 benchmark datasets. While all quantification pipelines exhibit similar ranking patterns based on Quartet reference datasets (top) and TaqMan datasets for Quartet (middle) and MAQC (bottom) samples, the Quartet reference datasets significantly enhances the distinguishment of quantification pipelines performance.

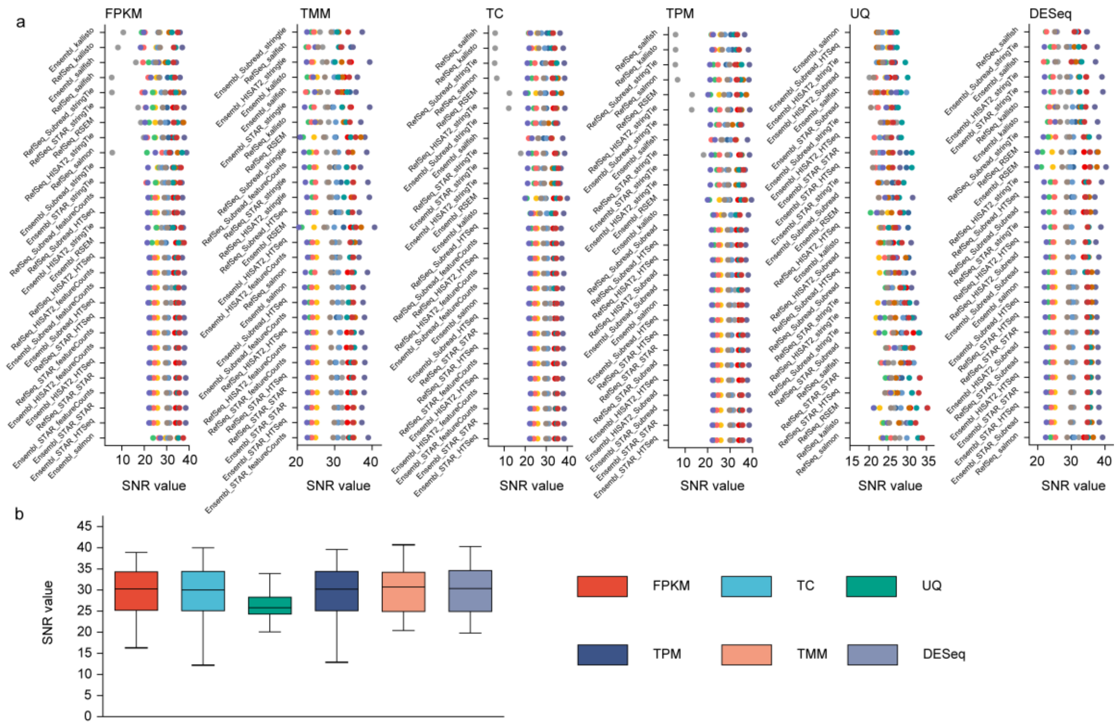

**Supplementary Figure 39. Comparison of the SNR of RNA-seq data from different quantification pipelines and normalization methods.** (a) SNR values for different combined pipelines. (b) Comparison of SNR values for different normalization methods with different choices of quantification pipelines.

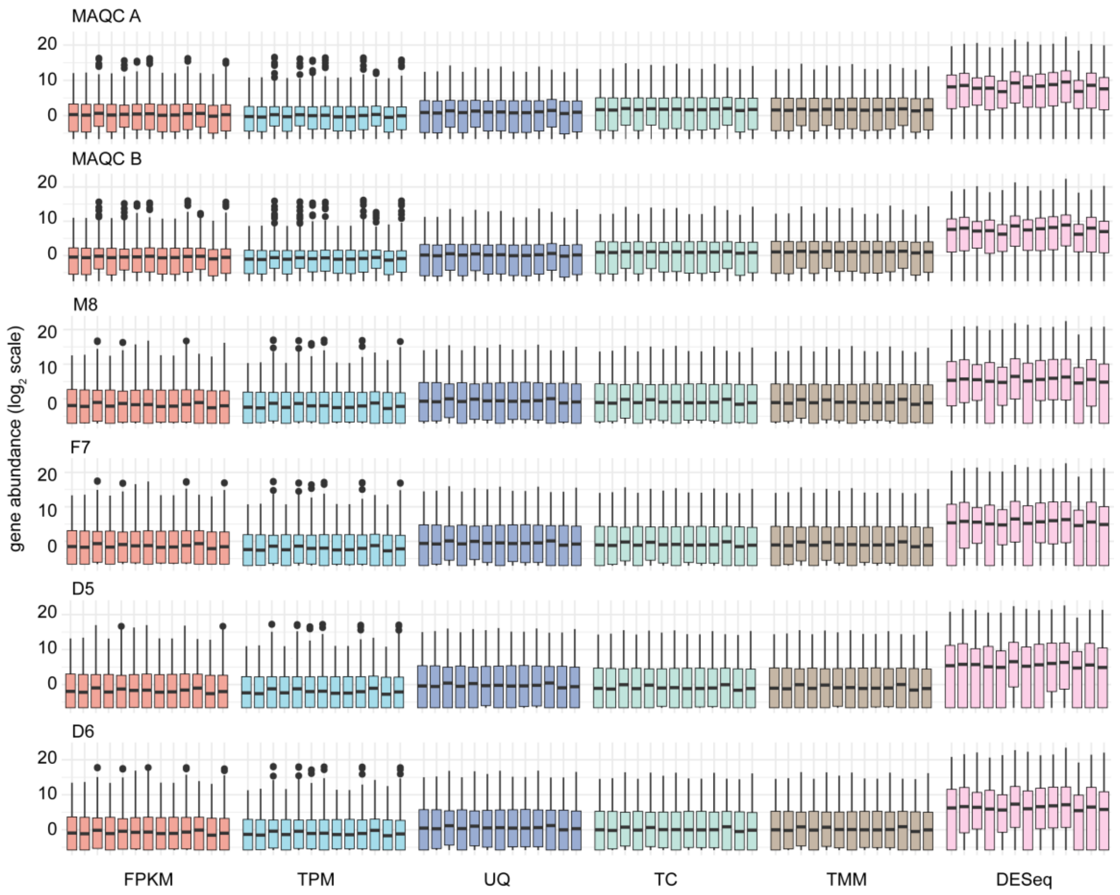

**Supplementary Figure 40. Distribution of gene expression using different normalization methods.** The gene expression from each normalization method were compared for the Quartet and MAQC samples.

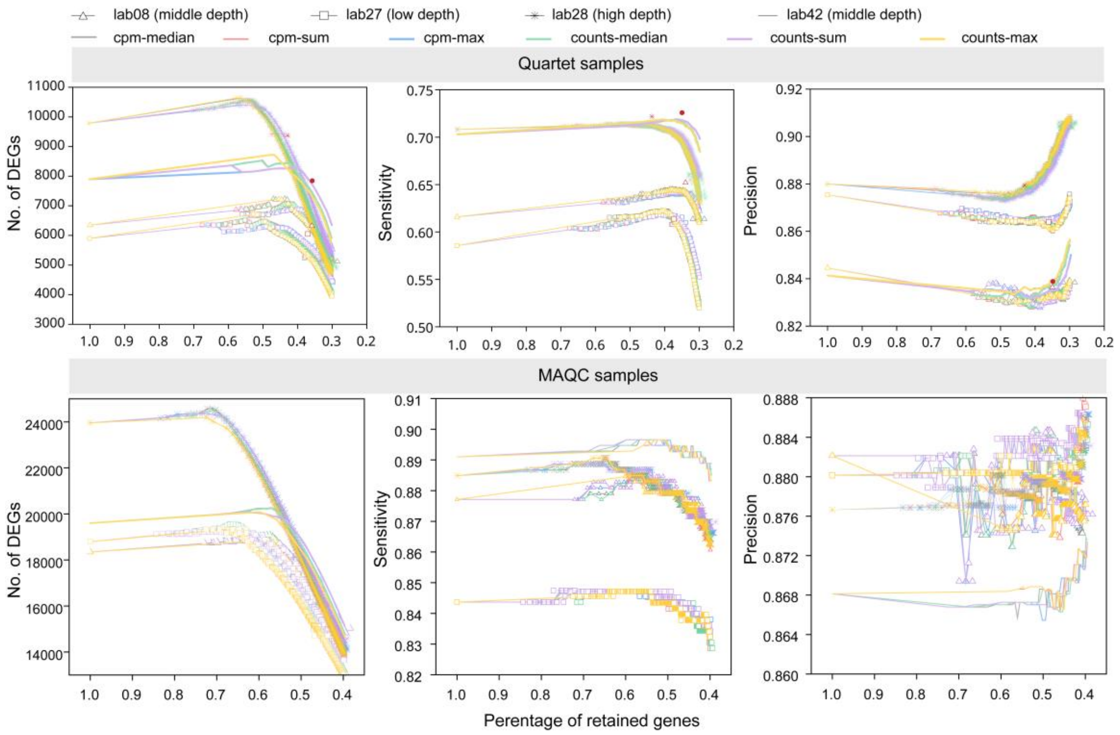

**Supplementary Figure 41. Quantitative assessment of low-expression gene filtering methods.** RNA-seq data representing high, medium, and low sequencing depth from four laboratories were employed for validation of different filtering conditions. Using the Ensembl-STAR-StringTie-edgeR pipeline, we calculated the number of DEGs, the true positive rate (sensitivity), and the precision as a function of filtering threshold for different filtering methods in Quartet samples (upper panel) and MAQC samples (lower panel). The red dots with different shapes represent the automated thresholds by edgeR.

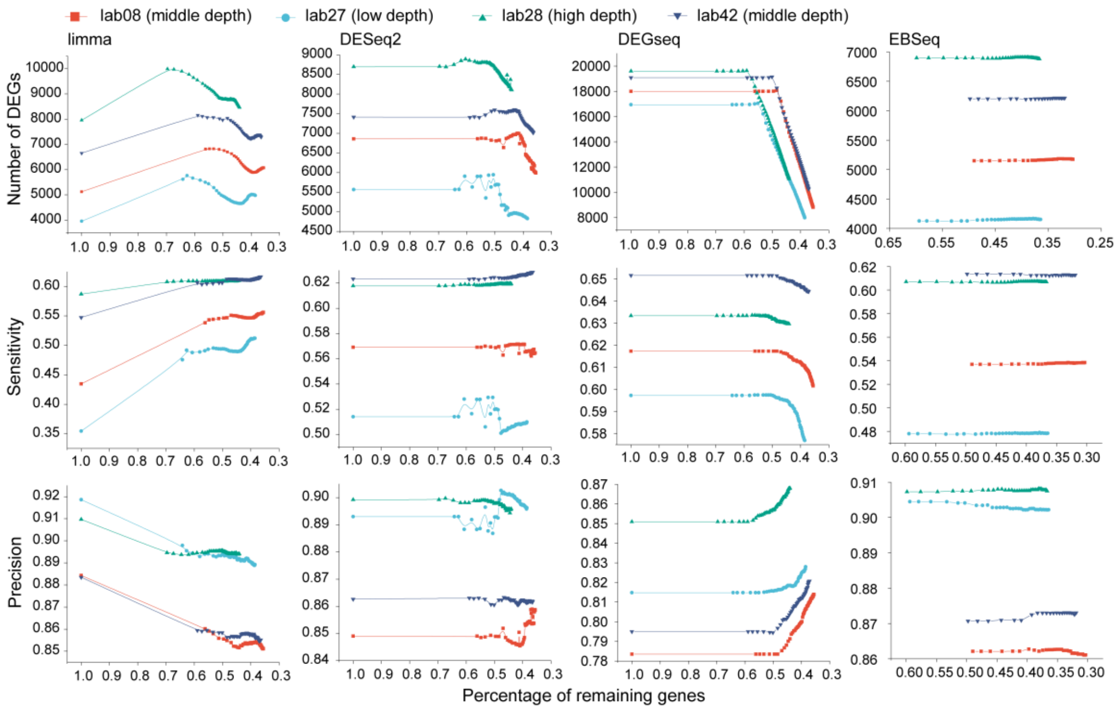

**Supplementary Figure 42. Quantitative assessment of low-expression gene filtering for different differential analysis tools.** For other tools including limma [6], DESeq2 [7], DEGseq [8], and EBSeq [9], we filtered low-expression genes based on the sum of reads counts, and calculated the number of DEGs, the true positive rate (sensitivity), and the precision as a function of filtering threshold.

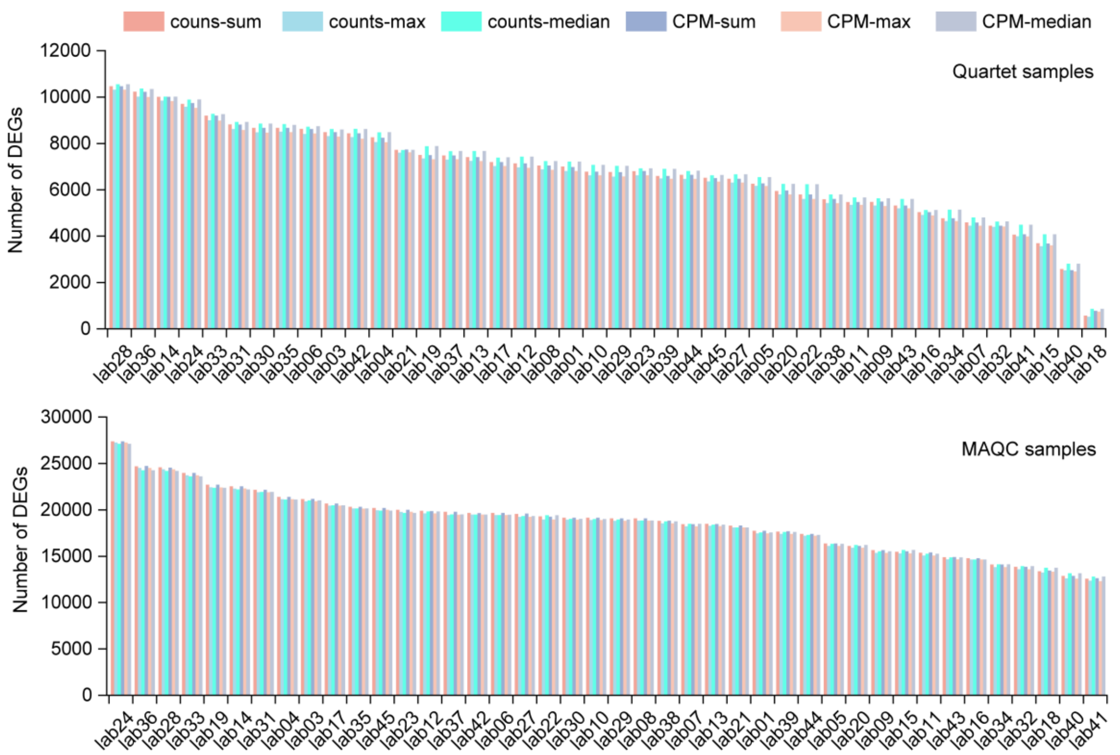

**Supplementary Figure 43. Comparison of the maximal number of DEGs using different filtering methods.** The maximal number of DEGs for Quartet (up) and MAQC (down) samples were calculated using edgeR after applying a series of filtering thresholds. DEGs, differentially expressed genes.

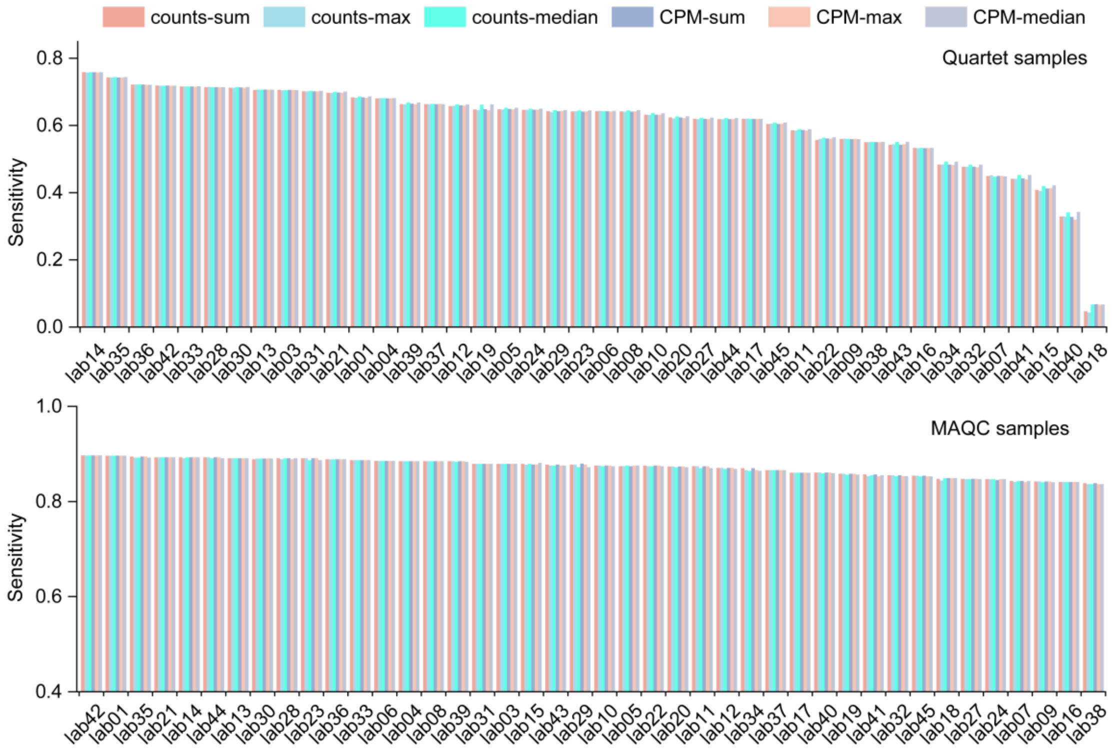

**Supplementary Figure 44. Comparison of the maximal sensitivity using different filtering methods.** The differential expression profiles for Quartet (up) and MAQC (down) samples at different threshold values were compared to Quartet reference datasets and MAQC TaqMan datasets, respectively, to calculate the maximal true positive rate. The edgeR was used for differential expression analysis. CPM, counts per million.

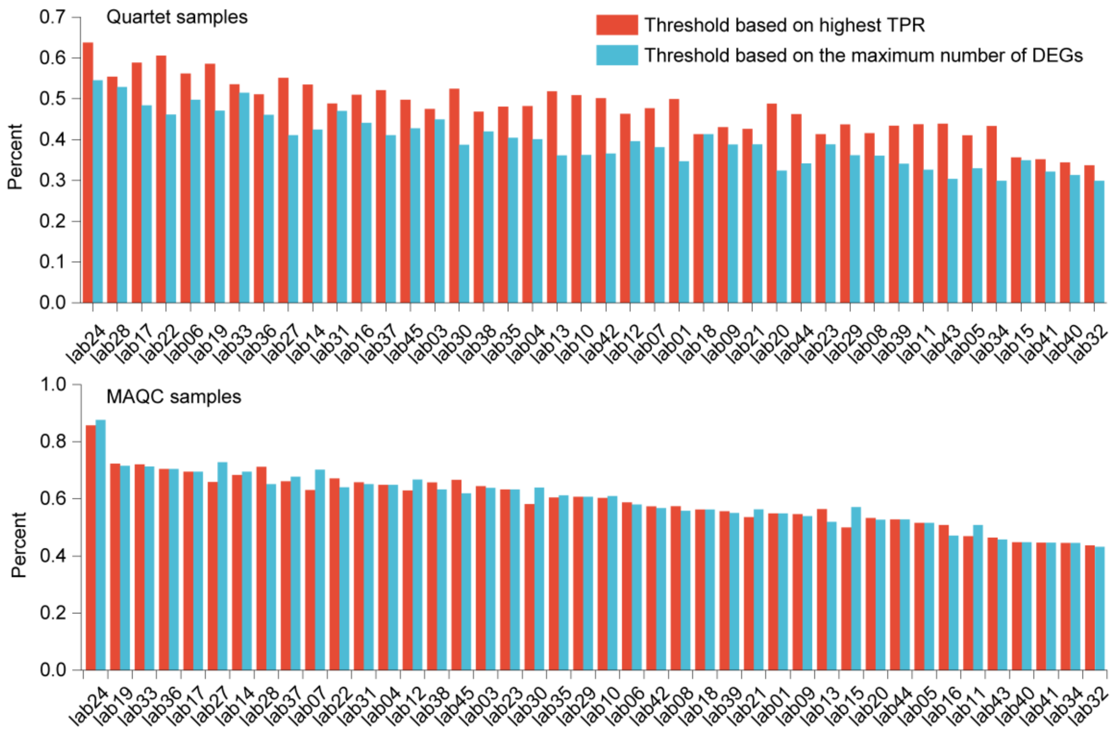

**Supplementary Figure 45. Comparison of the optimal threshold values determined by maximum number of DEGs and highest TPR.** For all RNA-seq from laboratories, the threshold values corresponding to the highest TPE and the maximum number of DEGs were calculated and compared for Quartet (up) and MAQC (down) samples. TPR, true positive rate.

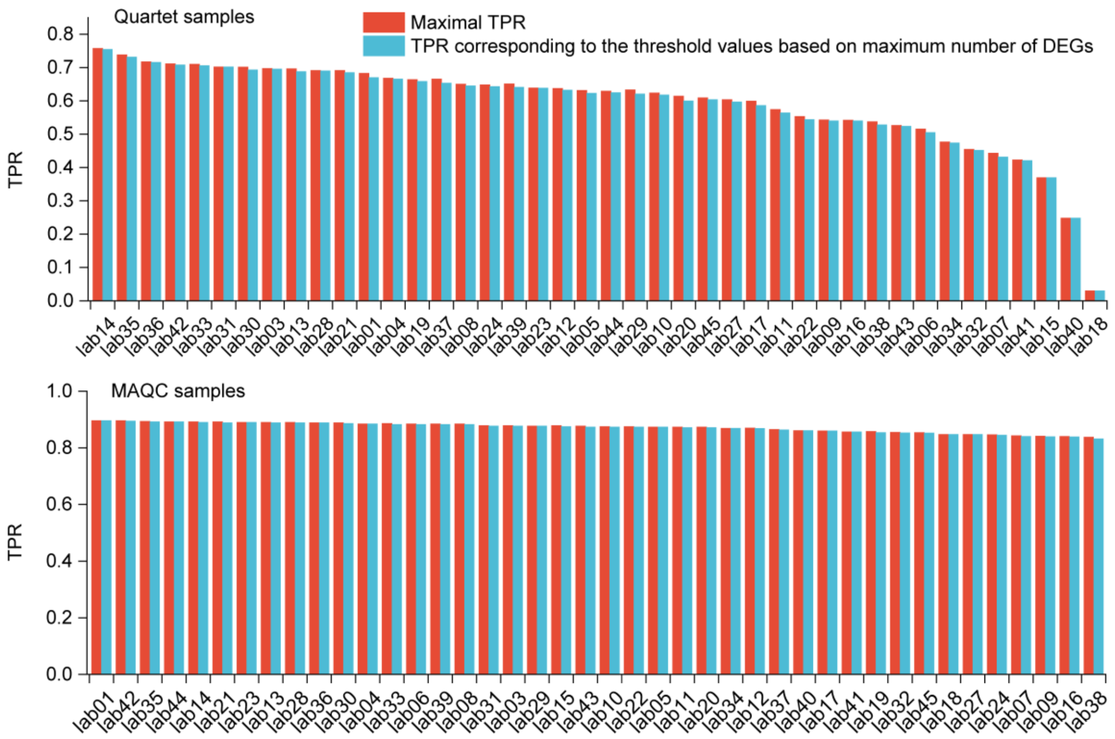

**Supplementary Figure 46. Comparison of the TPR corresponding to thresholds determined by maximum total number of DEGs and highest TPR.** The differential expression profiles for Quartet (up) and MAQC (down) samples at different thresholds were compared to the Quartet reference datasets and MAQC TaqMan datasets, respectively, to calculate the TPR. TPR, true positive rate.

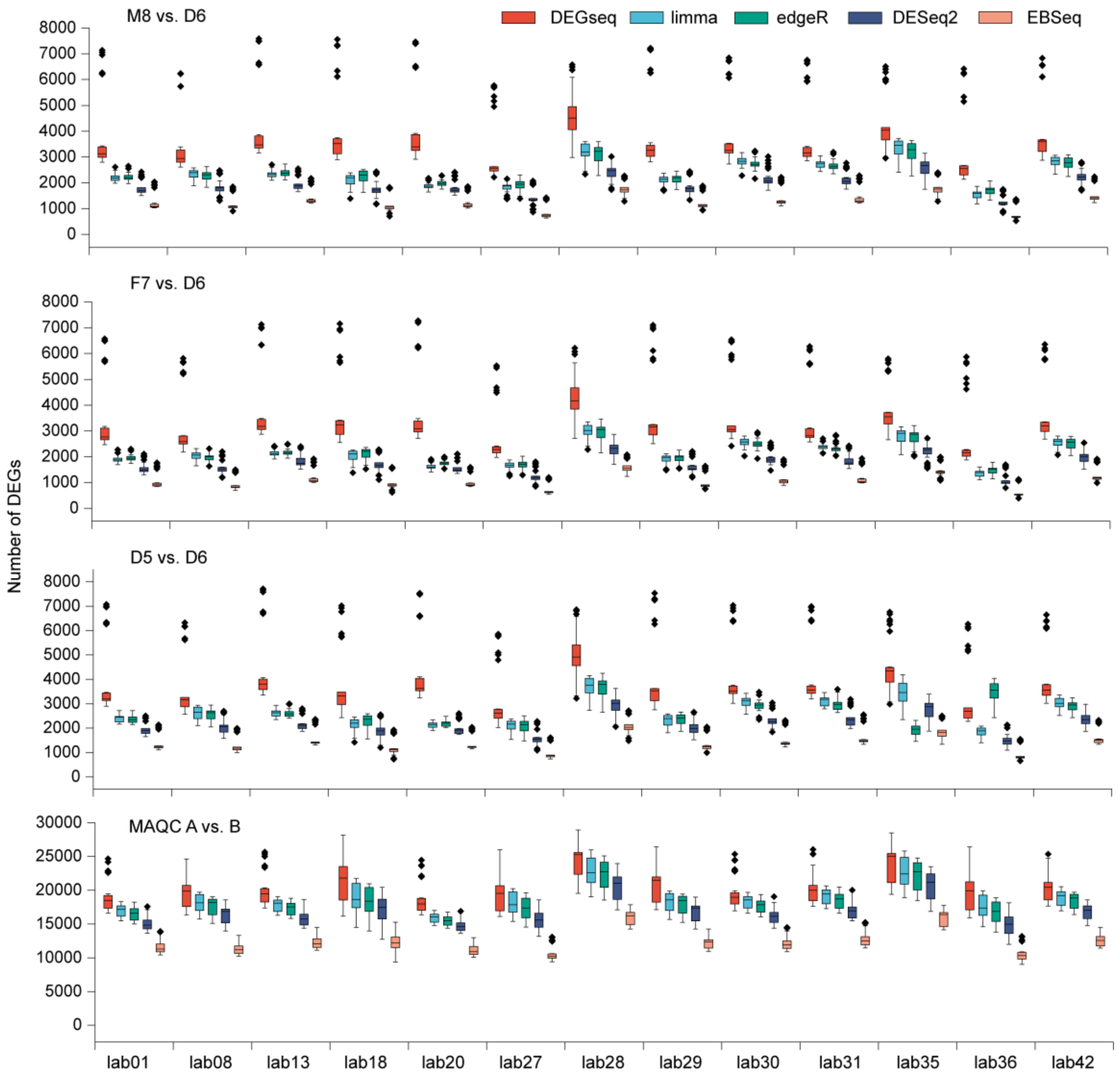

**Supplementary Figure 47. The number of differentially expressed genes (DEGs) detected by five differential analysis tools.** The number of DEGs was calculated for different differential analysis tools with the choice of different quantification pipelines in MAQC and Quartet samples. The pipelines including StringTie consistently resulted in a higher number of DEGs, depicted as outlier diamonds in the graphs.

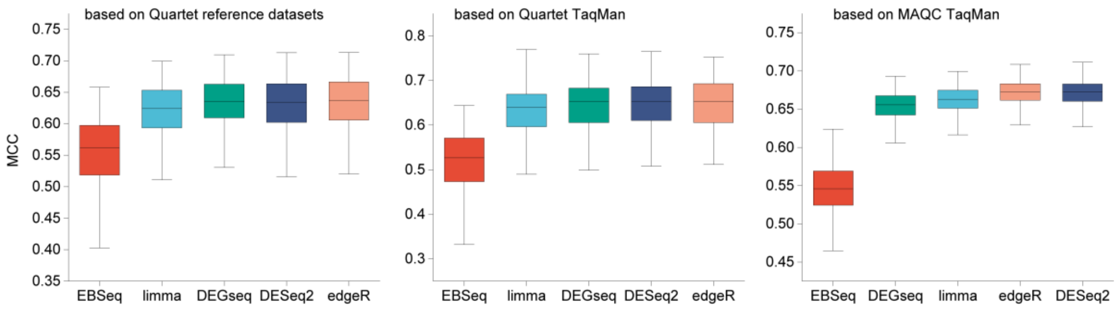

**Supplementary Figure 48. Comparison of performance of five differential analysis tools.** The box plots illustrate the performance of five differential analysis tools considering different gene annotations, alignment tools, quantification tools, and multiple samples. MCC, Matthews Correlation Coefficient.

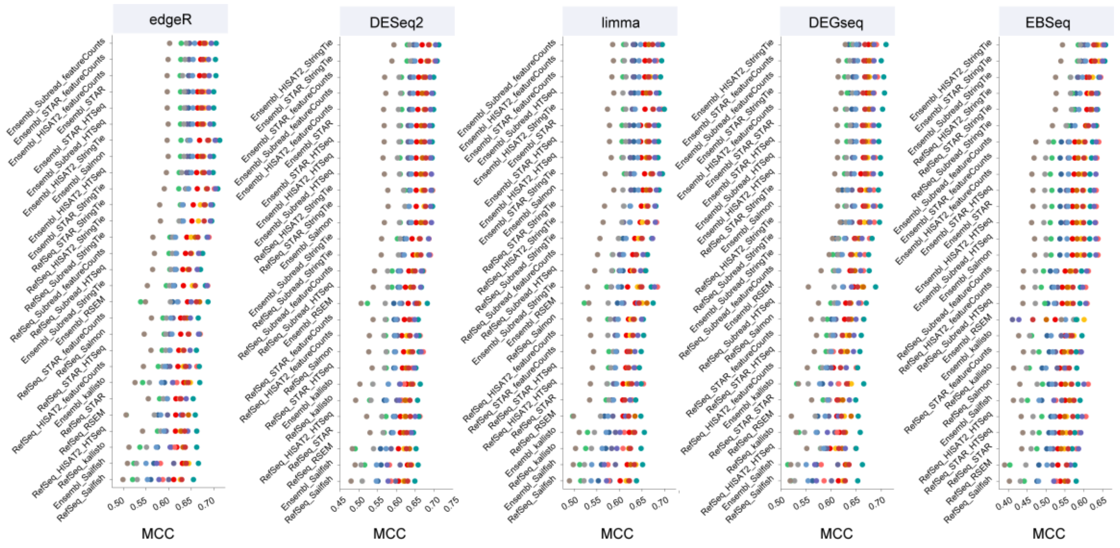

**Supplementary Figure 49. The influence of different quantification pipelines on five differential analysis tools.** The 140 differential analysis pipelines were assessed based on Quartet reference datasets. MCC, Matthews Correlation Coefficient.

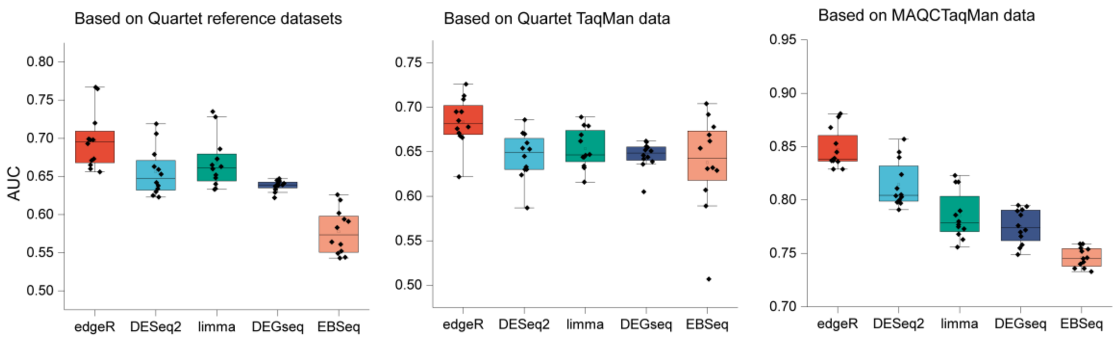

**Supplementary Figure 50. The assessment of five differential analysis tools using AUC values.** The box plots display the performance of each differential analysis tool in 13 benchmarked datasets. AUC, The area under the receiver operating characteristic curve.

### 2. Supplementary Notes

#### 2.1 The truth in sample panel

Our sample panel introduced three reference datasets including Quartet reference datasets, and TaqMan datasets for Quartet and MAQC reference materials, along with two 'built-in' including ERCC spike-in ratios and the mixing ratios in T1 and T2 samples.

First, Quartet project has developed a ratio-based reference datasets, which encompassed a total of 31,155 results from comparisons between M8, F7, and D5 with D6 [10]. A total of 5,036 differential expression genes (DEGs) were defined. After intersecting with the latest version of Ensembl gene annotation, the remaining count is 30,976, comprising 76.8% of protein-coding genes, 13.7% of long non-coding RNAs, 1.1% of small non-coding RNAs, and 6.4% of pseudogenes, and 1.9% of immunoglobulin/T-cell receptor gene segments.

Second, MAQC project has conducted 1,044 TaqMan RT-qPCR assays for MAQC samples. After removing genes with undetectable CT values (CT>35 or CT=0), 830 genes were remained. In comparison, we performed RT-qPCR experiments for 91 genes selected from the Quartet reference datasets, which provided new TaqMan datasets of 270 results for Quartet samples. A high level of concordance between the RT-qPCR and the reference datasets in terms of fold change was observed (Pearson correlation = 0.93), and 88% of differential gene expression results were consistent **(Supplementary Fig. 51)**.

a

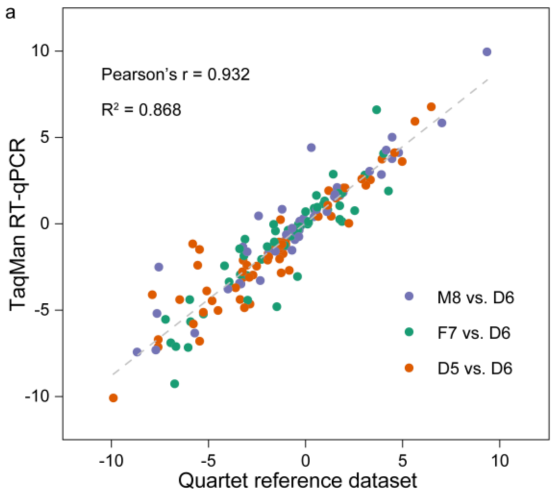

b

| TaqMan RT-qPCR | Quartet reference dataset: NonDEG | Upregulate | Downregulate |
|---|---|---|---|
| NonDEG | 37 | 8 | 10 |
| Upregulate | 2 | 41 | 0 |
| Downregulate | 2 | 8 | 84 |

**Supplementary Figure 51. Validation of ratio-based Quartet reference dataset.** (a) Scatter plots of $log_2$ fold changes (FC) of gene expression between Quartet reference datasets and TaqMan RT-qPCR. (b) Comparison of differentially expressed genes (DEGs) between Quartet reference datasets and TaqMan RT-qPCR. A total of 88% of genes with consistent classification results.

The distribution of relative gene expression for above Quartet reference datasets and TaqMan datasets for Quartet and MAQC samples was summarized in **Supplementary Fig. 52**. Genes in Quartet TaqMan datasets showed a relative lower expression level.

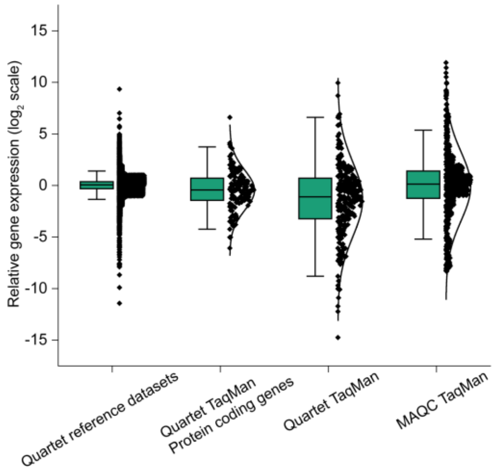

**Supplementary Figure 52. Comparison of the TaqMan and Quartet reference datasets.**

Third, the ERCC spike-in ratios provided a supplementary assessment of the accuracy and reproducibility. (i) All 92 ERCC genes in Mix 1 and Mix 2 have known wide-ranging concentrations, making them suitable for assessing absolute expression measurement. (ii) The ratio between Mix 1 and Mix 2 for four subgroups of ERCC genes were 4:1, 1:2, 2:3 and 1:1, respectively.

Fourth, sample M8 and D6 were mixed into T1 and T2 in a known ratios (3:1 and 1:3). Therefore, the relative gene expression between M8/D6, and T1/D6 or T2/D6 comparisons should adhere to the expected fitting curve (Materials and Methods). Additionally, based on known mixing ratios, the expected fold change for T1/D6 and T2/D6 comparisons can be calculated using fold change for M8/D6 comparison. The RMSE value between expected and observed fold change allows for an assessment of accuracy and reproducibility across all genes.

#### 2.2 The comprehensive performance assessment framework for RNA-seq data

The comprehensive performance assessment framework involved multifaceted properties of transcriptome data, including data quality, gene expression, and differential gene expression (**Fig. 1b**).

First, the quality of expression data was quantified using the PCA-based signal-to-noise ratio (SNR) values, which measures the laboratory's ability to distinguish biological signals between different samples from the technical noise of replicates. PCA-based SNR values were highly effective in discriminating data quality across laboratories, especially when working with Quartet samples characterized by subtle biological differences.

Second, the gene expression was examined by calculating the accuracy and reproducibility of absolute and relative expression. Our assessment extended to cover

92 ERCC genes, hundreds of genes in TaqMan datasets, tens of thousands of genes in Quartet reference datasets, and all genes through examining the mixing ratios recovery. The inclusion of a broader set of gene significantly enhanced the assessment precision. Third, the number of differentially expressed genes (DEGs) were compared across laboratories, and the accuracy of DEG calls were evaluated based on Quartet reference datasets and TaqMan datasets for Quartet and MAQC samples.

This comprehensive performance assessment framework addressed several challenges (**Supplementary Figure 53**). (i) the study addressed challenges related to gene annotations from different sources and versions used in real-world laboratories, which typically hindered comparisons across laboratories and reference datasets [29]. We matched gene IDs of different types and versions to generate an intersection gene set, facilitating comparisons of transcriptome data. (ii) the data quality, differential analysis tools, and filtering parameters all influence the differential gene expression list, which may lead to certain genes present in reference datasets being missed, raising the question of how to categorize these genes in performance metric calculation. We excluded genes unreported by laboratories due to the choice of different gene annotations and defined the remaining unreported genes labeled as DEGs in the reference dataset as false negatives for calculating the MCC metric. Altogether, this performance assessment framework facilitates the integration and comparison of real-world transcriptome data. Moving forward, with the emergence of omics reference datasets like Quartet, there is a growing need for standardized benchmarking tools available for open access, providing a one-stop solution for comprehensive RNA-seq performance assessment [30].

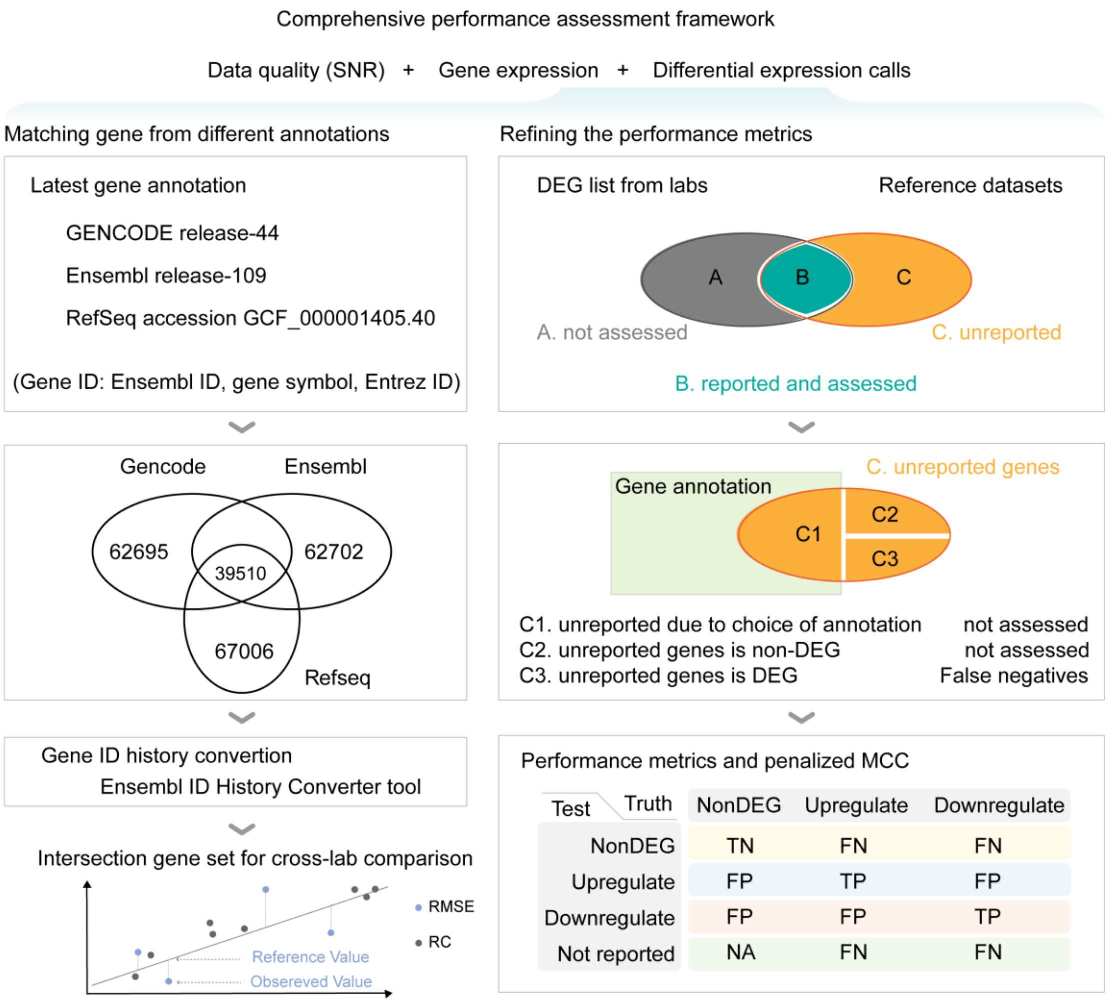

**Supplementary Figure 53. The performance assessment framework for RNA-seq data.**

### 2.3 RNA-seq assessment based on ERCC spike-in controls.

The reads mapped to the ERCC genes should account for approximately 1% of the total exonic reads, providing an assessment of library quality. Laboratories demonstrated significant variations regarding the percentage of reads mapped to ERCC **(Supplementary Fig. 54)**. The rRNA depletion protocol correlated to a higher fraction of reads aligned to ERCC genes, while the poly(A) selection protocol exhibited a lower mapping ratio but better recovered the relative ratio among the four subgroups of ERCC sequences **(Supplementary Fig. 55)**.

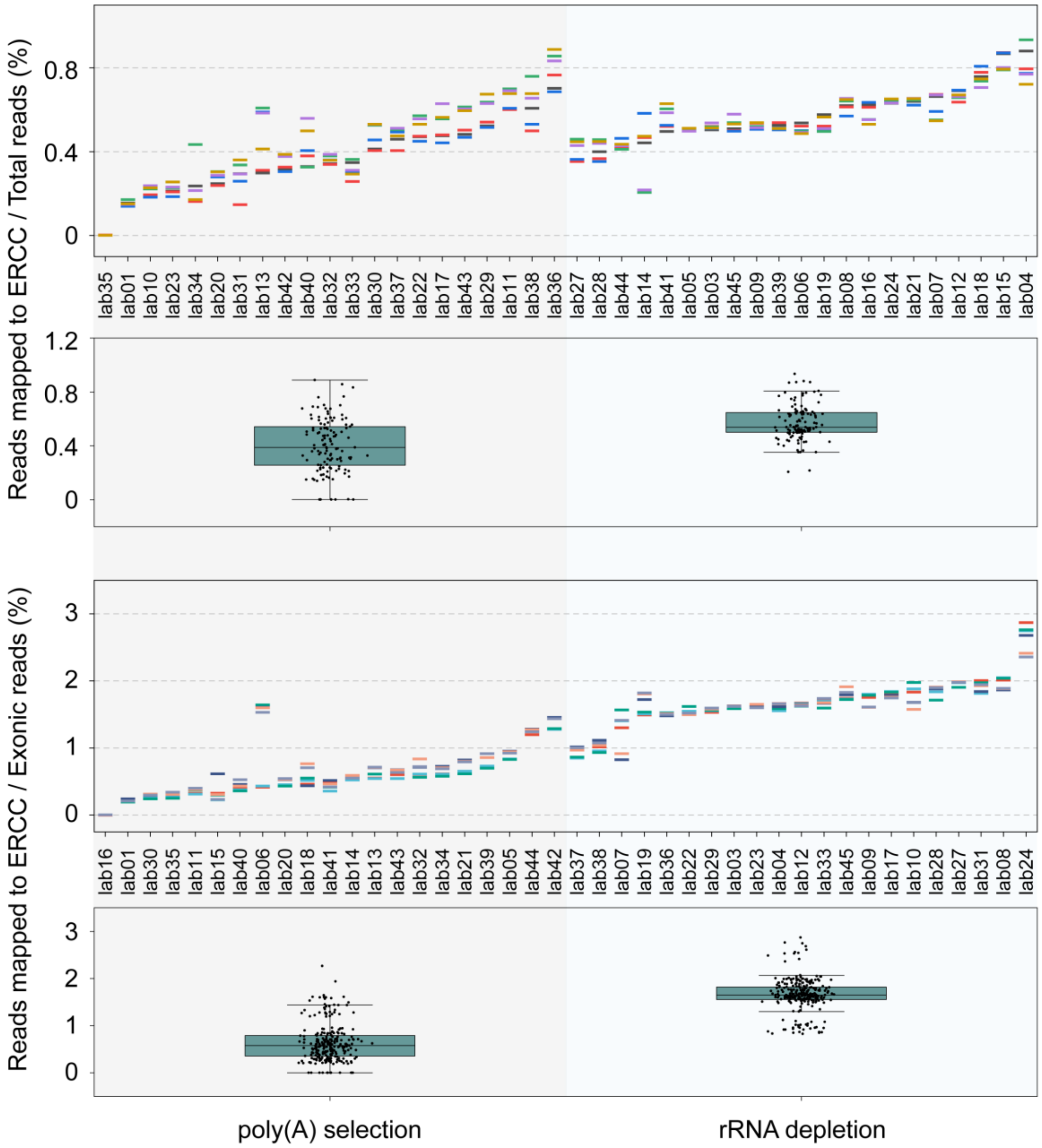

**Supplementary Figure 54. Percentage of reads mapped to ERCC genes.** The rRNA depletion protocol correlated to a higher fraction of reads mapped to ERCC genes.

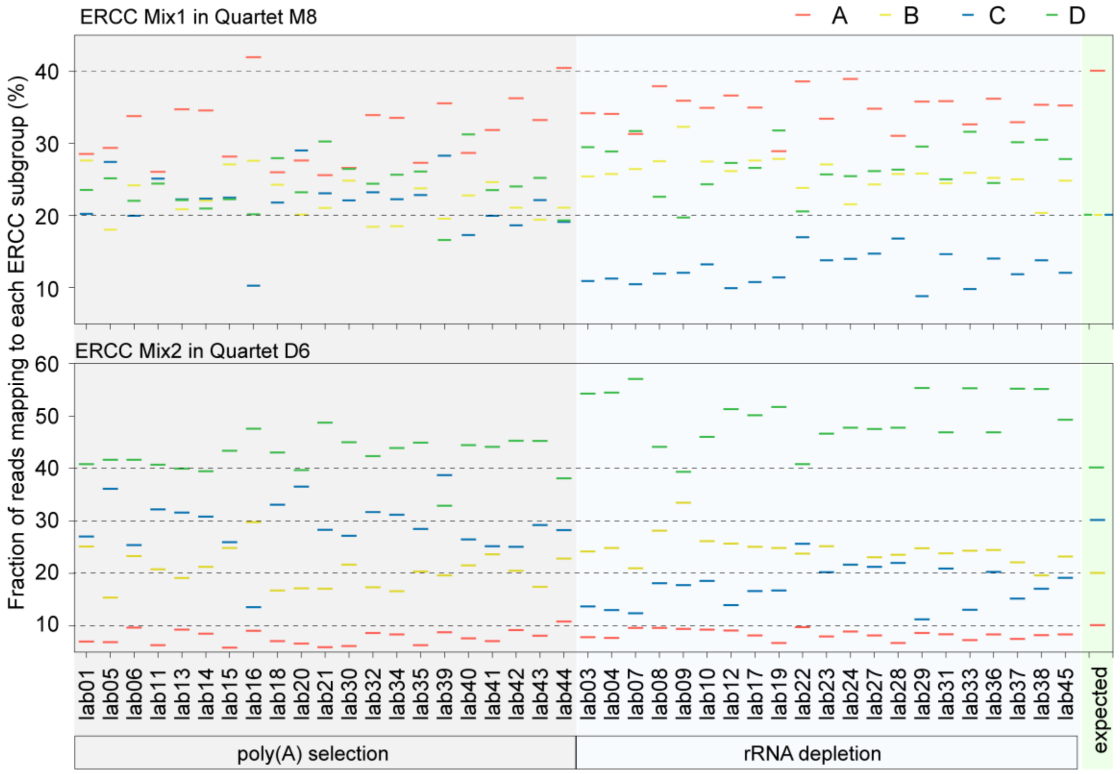

**Supplementary Figure 55. Percentage of reads mapped to four subgroups of ERCC genes.** Both ERCC Mix 1 and Mix 2 contain genes from four subgroups labeled as A, B, C, and D, each with 23 genes. The expected ratios of A, B, C, and D, are 2:1:1:1 and 1:2:3:4 in Mix 1 and Mix 2, respectively [11]. The poly(A) selection protocol associated with higher accuracy of recovering the intrinsic ratios.

### 2.4 The number of detected genes in Quartet and MAQC samples.

We compared the number of detected genes across all laboratories after applying the same threshold. Genes with at least one reads in all replicates were involved for analysis. The MAQC samples consistently exhibited a higher number of detected genes across laboratories, ranging from 18,146 to 53,743, while the Quartet samples had comparatively fewer genes, ranging from 16,401 to 46,883, and the mixed samples had the fewest genes, ranging from 16,064 to 45,771. The substantial variations in the number of detected genes among laboratories were mainly associated with specific gene types of interest and sequencing depth **(Supplementary Fig. 56)**. Noticeably, the relatively lower number of detected genes in some laboratories despite with high depth could be attributed to low read alignment rates, high duplication rates, and gene annotations PCR duplicates is undesirable since it reduces the effective sequencing depth of the experiment **(Supplementary Fig. 56)**. Similarly, increased sequencing depth contributed to the detection of more ERCC genes **(Supplementary Fig. 57)**.

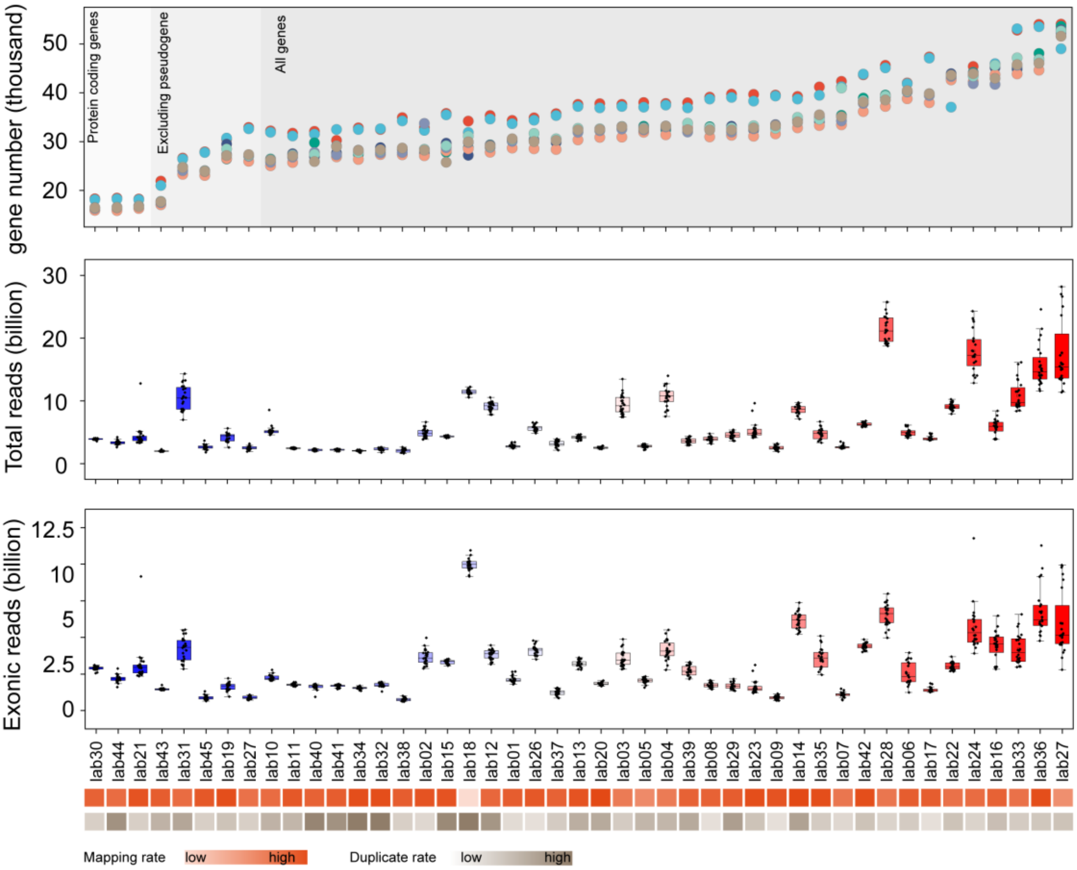

**Supplementary Figure 56. The number of genes detected by all laboratories.** The genes supported by at least one reads for all three replicates were included for analysis. Higher sequencing depth generally leads to the detection of more genes, although exceptions exist, such as when alignment rates are low or duplicate rates are high.

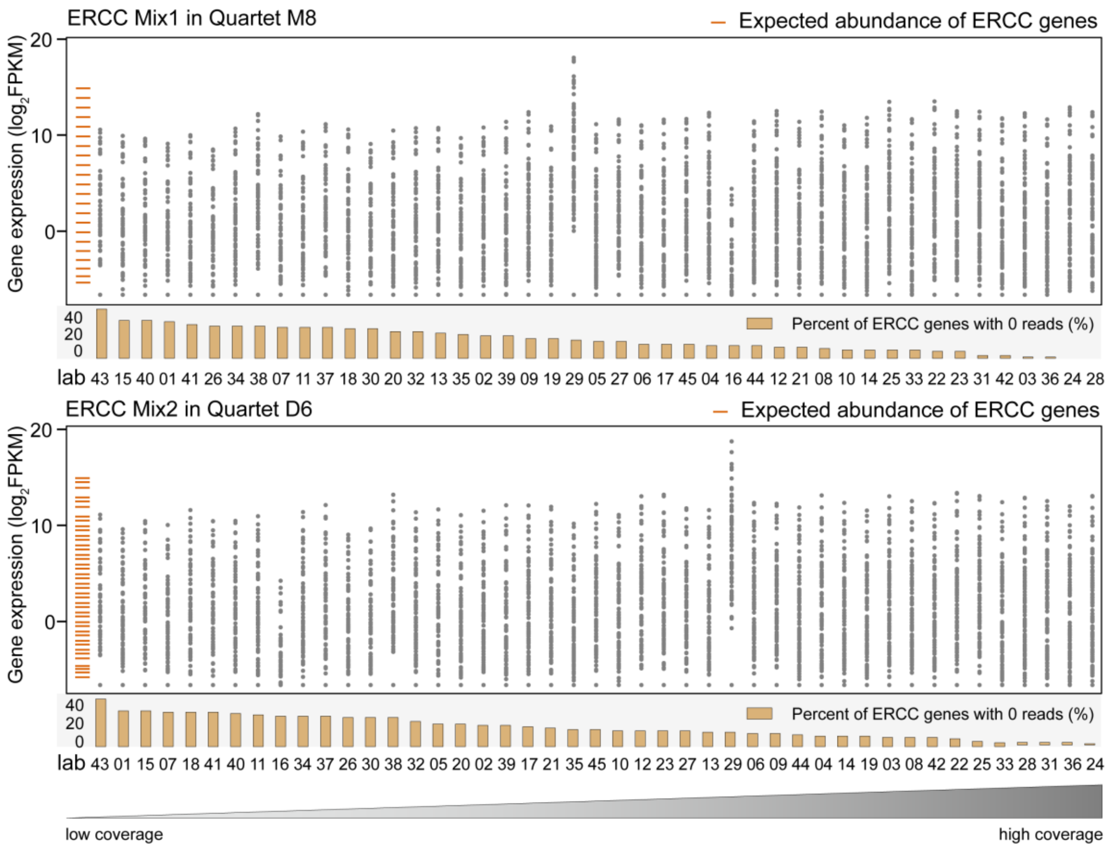

**Supplementary Figure 57. The number of ERCC genes detected by 42 laboratories.**

With the increase of sequencing depth, more ERCC genes were supported by at least one reads.
